# Smurfness-based two-phase model of ageing helps deconvolve the ageing transcriptional signature

**DOI:** 10.1101/2022.11.22.517330

**Authors:** Flaminia Zane, Hayet Bouzid, Sofia Sosa Marmol, Mira Brazane, Savandara Besse, Julia Lisa Molina, Céline Cansell, Fanny Aprahamian, Sylvère Durand, Jessica Ayache, Christophe Antoniewski, Nicolas Todd, Clément Carré, Michael Rera

**Affiliations:** Université Paris Cité, INSERM UMR U1284, 75004 Paris, France; Sorbonne Université, Institut de Biologie Paris Seine, 75005, Paris, France; Université Paris-Saclay, AgroParisTech, INRAE, UMR PNCA, 91120, Palaiseau, France; Metabolomics and Cell Biology Platforms, UMS AMMICa, Institut Gustave Roussy, Villejuif 94805, France; Centre de Recherche des Cordeliers, Equipe labellisée par la Ligue contre le cancer, Université de Paris, Sorbonne Université, INSERM U1138, Institut Universitaire de France, Paris 75006, France, Université Paris Cité, Institut Jacques Monod, CNRS UMR 7592, 75013 Paris, France; Eco-Anthropologie (EA), Muséum National d’Histoire Naturelle, CNRS, Université de Paris, Musée de l’Homme, Paris, France

**Keywords:** ageing, transcriptome, end of life, Smurfs, drosophila, lifespan increasing genetic intervention

## Abstract

Ageing is characterised at the molecular level by six transcriptional ‘hallmarks of ageing’, that are commonly described as progressively affected as time passes. By contrast, the ‘Smurf’ assay separates high-and-constant-mortality risk individuals from healthy, zero-mortality risk individuals, based on increased intestinal permeability. Performing whole body total RNA sequencing, we found that Smurfness distinguishes transcriptional changes associated with chronological age from those associated with biological age. We show that transcriptional heterogeneity increases with chronological age in non-Smurf individuals preceding the other five hallmarks of ageing, that are specifically associated with the Smurf state. Using this approach, we also devise targeted pro-longevity genetic interventions delaying entry in the Smurf state. We anticipate that increased attention to the evolutionary conserved Smurf phenotype will bring about significant advances in our understanding of the mechanisms of ageing.

**Graphical abstract:** The two-phase model of ageing allows to study separately the effect of chronological and physiological age. **(A)** Classic approaches for studying ageing tend to consider it as a black box affecting all individuals progressively from birth to death. Instead, the Smurf phenotype shows that life can be divided into two consecutive phases separated by an abrupt transition. **(B)** All individuals undergo this transition at a different moment in their life, prior to death. This allows us to switch from population based approaches, comparing bulks of age-matched individuals through time, to individuals-centred approaches relying on direct access to their transition status. **(C)** Such paradigm shift shows that hallmarks of ageing long thought to progressively change with age are actually mostly affected in a growing proportion of Smurfs, allowing for the identification of the chain of events accompanying ageing and death from natural causes. **(D)** By studying the behaviour of the ageing transcriptome as a function of chronological age and Smurfness separately, we demonstrate that the progressively changing transcriptional ageing signature, as described in Frenk & Houseley (2018), is in fact the convolution changes accompanying chronological age signature (increased transcriptional noise) and changes associated with Smurfness (or biological age) signature (increased stress response and inflammation, decreased expression of ribosomal and mitochondrial genes). We also identified a hallmark partially associated with only old Smurfs (ATH5), suggesting that chronological age can affect, late in life, the Smurf response.

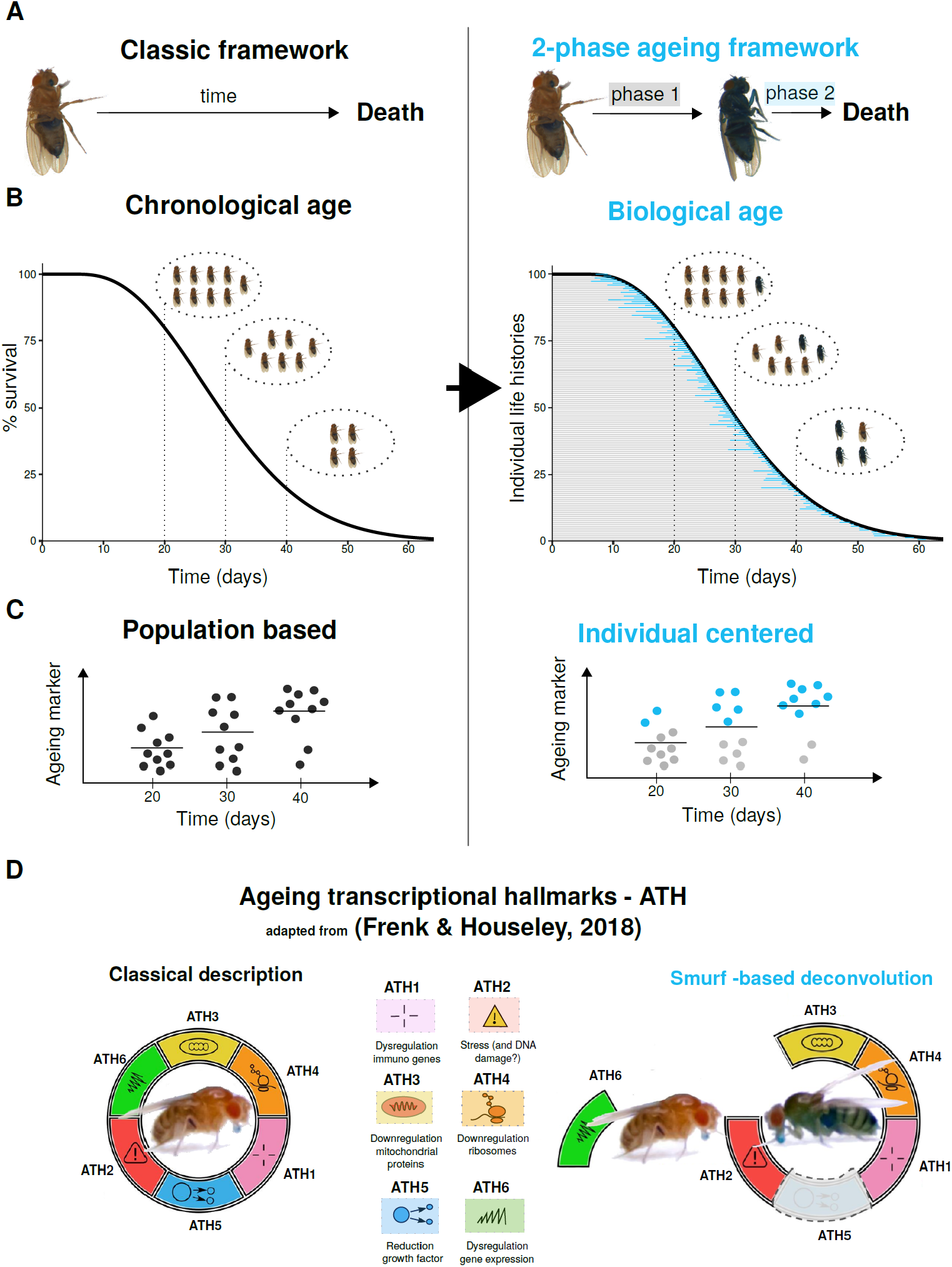

## Introduction

### Chronological age and physiological ageing

Ageing is commonly defined as a progressive decrease in functional efficiency associated with an age-related increasing vulnerability to death^1, 2^, although different modalities can be found across the livings^3^. In a given population, individuals of the same chronological age can yet experience different risks of mortality, showing that physiological ageing is not fully captured by chronological age. In humans, the notion of frailty - an unobserved individual modulator of the force of mortality - was introduced to explain this heterogeneity^4^. It was followed by the definition of specific frailty indexes, fixed sets of characteristics that can be used to predict an individual’s risk of death independently of its chronological age^5–7^. On the one hand, the use of such frailty indexes has now been extended to several model organisms^8–10^. On the other hand, efforts to define ageing at the cellular and molecular levels have led to the definition “hallmarks of ageing”^1, 11, 12^, evolutionary conserved molecular markers progressively affected in ageing individuals - and to the development of ageing clocks predicting biological age based on molecular markers. Ageing clocks based on 5-cytosine methylation of CpG sites^13–16^ work well in mammals but do not apply to model organisms such as *Caenorabditis elegans* or *Drosophilia melanogaster.* Nevertheless, recent work has identified a “universal” transcriptomic clock using *C. elegans*^17^, with the recent publication of the BiT age clock^18^, suggesting a possible conservation of critical biological age markers.

### The Smurf approach to ageing

The Smurf assay is an *in vivo* non-invasive assessment of increased intestinal permeability (IP) based on co-ingestion of the non-toxic blue food dye FD&C #1 (approx. 800Da). The dye, normally not absorbed by the digestive tract, spreads throughout the body in flies with altered IP, turning them blue^19, 20^, hence their name Smurfs. The Smurf assay was previously shown to be a powerful marker of biological age in *D. melanogaster*^19^ as well as other model organisms^21^. Maintaining a population on standard food containing the dye reveals that the proportion of Smurfs increases as a function of time^19^ and that all flies undergo the Smurf transition prior to death^19, 20^. Furthermore, Smurf flies present a low remaining life expectancy (T_50_ estimated at ∼ 2.04 days across different genetic backgrounds from the DGRP set^22^) that appears independent of their chronological age at Smurf transition^19, 20^. In a given population at any given age, the Smurfs are the only individuals showing high mortality risk, low energy stores, low motility, high inflammation and reduced fertility, making this subpopulation a characteristic frail subpopulation. We demonstrated, thanks to a simple two-phase mathematical model, that we are able to describe longevity curves using the age-dependent linear increase (approximation) of the Smurf proportion and the constant force of mortality in Smurfs^20^.

The above-mentioned studies led us to hypothesise that markers classically considered as progressively and continuously changing during ageing (the hallmarks of ageing) might actually accompany the Smurf transition and exhibit a biphasic behaviour (two-phase model of ageing^20, 23^. The age-dependent increase in mortality at the population-level should then be re-interpreted as the increasing proportion of Smurfs in the population of individuals still alive^20^. To test this hypothesis, we assessed the transcriptional changes occurring in flies as a function of both their Smurf status and chronological age. RNA-Sequencing (RNA-Seq) was performed on Smurf and non-Smurf individuals of different chronological ages after total RNA extraction from the whole body of mated female flies. Samples were collected at 20, 30 and 40 days after eclosion, corresponding to approximately 90%, 50% and 10% survival in the used line (*Drs*-GFP;Fig. S1-2).

## Results

### Smurfs have a stereotypical transcriptome

We first performed a Principal Component Analysis (PCA) to explore how our multiple samples did relate to each other. The first component (45% of variance) separates Smurfs and non-Smurfs samples (Fig. 1a). This component is significantly associated with Smurfness (R² ANOVA = 0.604, p-value = 1.67e^-07^), but not with age (p-value > 0.05). The second component (13%) segregates samples as a function of chronological age (Pearson ϱ = 0.717, p-value = 3.92e^-06^), with no significant association with Smurfness (p-value > 0.05). The fact that three 40 days Smurfs samples out of six clusters with same age non-Smurfs, a pattern confirmed using independent tSNE (t-distributed stochastic neighbour embedding) and hierarchical clustering on sample-to-sample distance (Fig. S3 and S4), indicates fewer differences between the transcriptomes of old Smurfs and non-Smurfs than between young ones.

**Figure 1.**
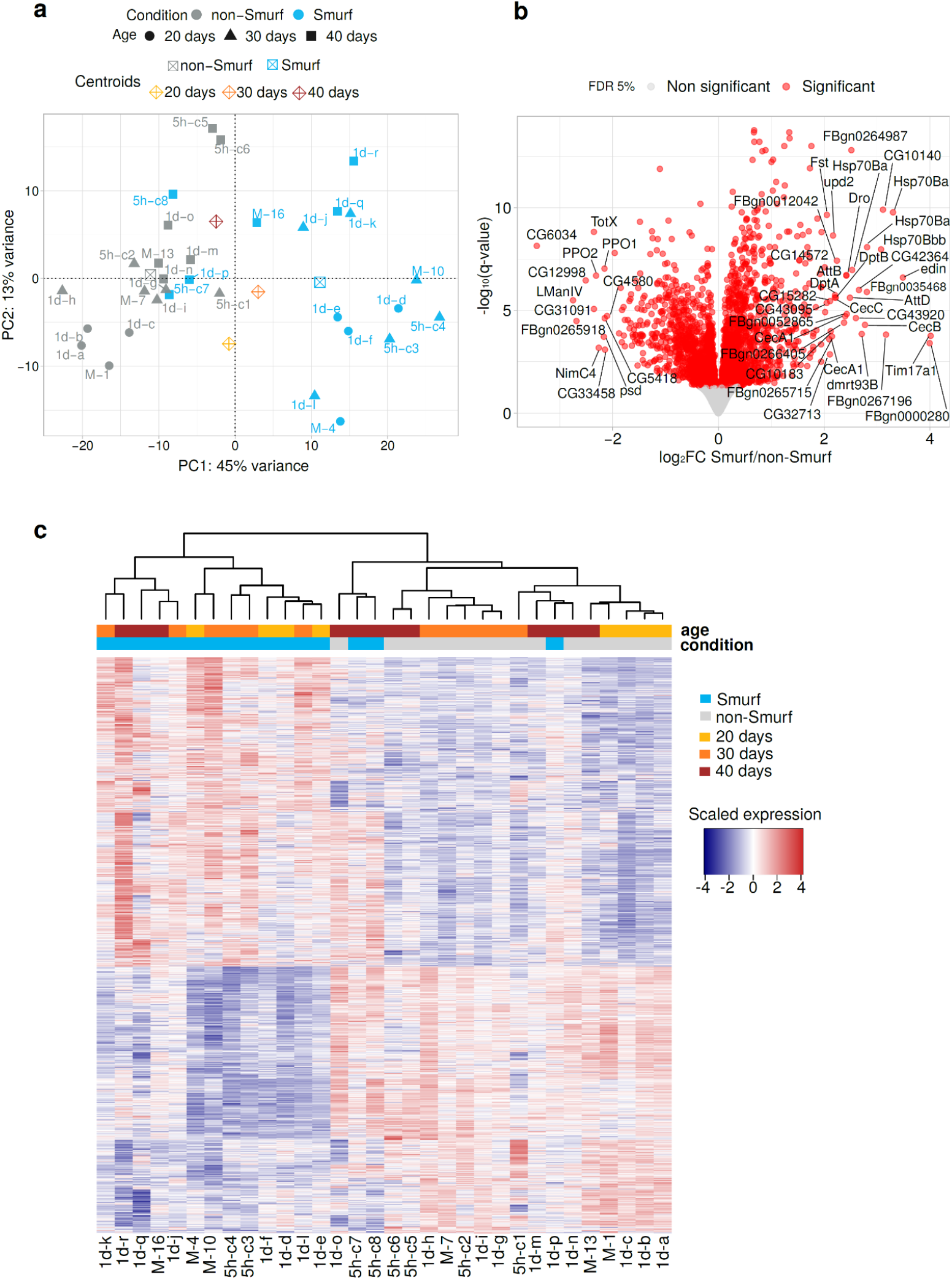
Smurfness is associated with a characteristic transcriptome. **a)** Samples plotted in the space of the first two PCA components. PCA performed on the 1000 top-variance genes results in a clear separation of Smurf (blue) and non-Smurf (grey) samples on PC1 while samples are distributed according to age on PC2. This shows that Smurfness explains most of the transcriptome variance in our dataset (45% for PC1), followed by age (13% for PC2). Shapes indicate the age as illustrated in the legend. Centroids coordinates for a specific group are the mean of the group coordinates. Each sample is associated with an acronym specifying the collection time after the transition (5h = 5 hours, 1d = 1 day and M = mixed - unknown time -) and a unique letter or number identifying the sample itself. **b) Volcano plot of the DEG analysis results.** The negative logarithm with base 10 of the FDR adjusted p-value (q-value) is plotted as a function of the shrinked (DESeq2 apeglm method^26^) fold change (base 2 logarithm) of the Smurf/non-Smurf expression ratio for each gene. The significant 3009 DEGs are represented in red. Upregulated Smurf genes (1618) plot on the right side of the graph, while downregulated genes (1391) on the left. Genes with a log_2_FC > |2| are labelled. Amongst the genes annotated as upregulated we can notice the presence of immune response genes (*Dro*, *AttB*, *AttC*, *DptA*, *DptB*, *CecA1*, *CecB*, *CecC*), confirming what already described in Smurfs^19^. **c) Smurf DEGs represent a Smurf specific signature.** Unsupervised hierarchical clustering on the samples by Smurf DEGs only divides Smurfs from non-Smurfs independently of their age, demonstrating that those genes are a Smurf specific signature. Non-Smurf samples tend to cluster by age, suggesting an age trend in the expression of Smurf DEGs in non-Smurf. The same three outliers of **(a)** are identified, indicating that those three samples indeed present a weaker expression pattern compared to the other Smurfs. Expression of genes in the heatmap is re-centered on the mean across samples, for easy visualisation of upregulated and downregulated genes.

We proceeded to quantify the differences between Smurfs and non-Smurfs through differential gene expression analysis (DESeq2^24^). Comparing the 16 Smurf and the 16 non-Smurfs samples, we identified 3009 differentially expressed genes (DEGs)(Fig 1b, DESeq2 results in Supplementary File 1). Confirming the PCA results, these genes represent a Smurf-specific signature that clusters the Smurfs samples (Fig 1c). Again, the effect of chronological age is less marked in Smurf samples than in non-Smurf ones. DESeq2 results were validated using the edgeR^25^ pipeline, which identified 2609 DEGs, 90% of which are overlapping with the DESeq2 output and present a strong correlation (Pearson ϱ = 0.99) for log_2_FC estimation (Fig. S5).

### Smurfness recapitulates the transcriptional signature of ageing

We used biological processes (BP) Gene ontology (GO)^27^ as gene sets in Gene Set Enrichment Analysis (GSEA)^28^ to characterise the Smurf signature. In order to fully examine the observed signal, we chose not to apply any filtering on the log_2_FC (FC: fold change). We mapped our results on the hallmarks of transcriptional ageing (ATH 1-6) described in Frenk et al.^29^ on the GSEA network (Fig. 2, Tab. S1). Genes upregulated in Smurfs are enriched in immune and stress response (ATH1), as previously reported in Smurfs^19^ as well as numerous ageing transcriptomic studies in Drosophila^30–35^ and other organisms^36–41^ including humans^42^. Here, the immune response is widely upregulated, with activation of both Toll (fungi and Gram-positive response)^43^ and Immune deficiency (Imd, Gram-negative response)^44, 45^ pathways. Antimicrobial peptides (AMPs), which are surrogates of inflammation in flies, are strongly upregulated (*CecA1*, *CecA2*, *CecB*, *CecC*, *DptA*, *Def*, *Dpt*, *Drs*, average log_2_FC=2.33) with their upstream regulators *Rel* (Imd pathway, log_2_FC=0.61) and *dl* (Imd pathway, log_2_FC=0.27) also upregulated. Stress responses (ATH2) such as protein folding and unfolded protein response (UPR, with upregulation of *Xbp1* and *Ire1*) are over represented in our dataset. Smurfs present a significant induction of 22% of Drosophila chaperons and co-chaperons (Flybase^46^ annotation, version FB2022_04), with a broad upregulation of the Hsp70 family (6 out of 7 genes detected are upregulated, average log_2_FC=2.60), as previously described in ageing^33, 47^. We detect a significant upregulation of 51% of the annotated cytosolic Glutathione S-transferases (Gst), a family of genes involved in detoxification and oxidative stress response.

**Figure 2.**
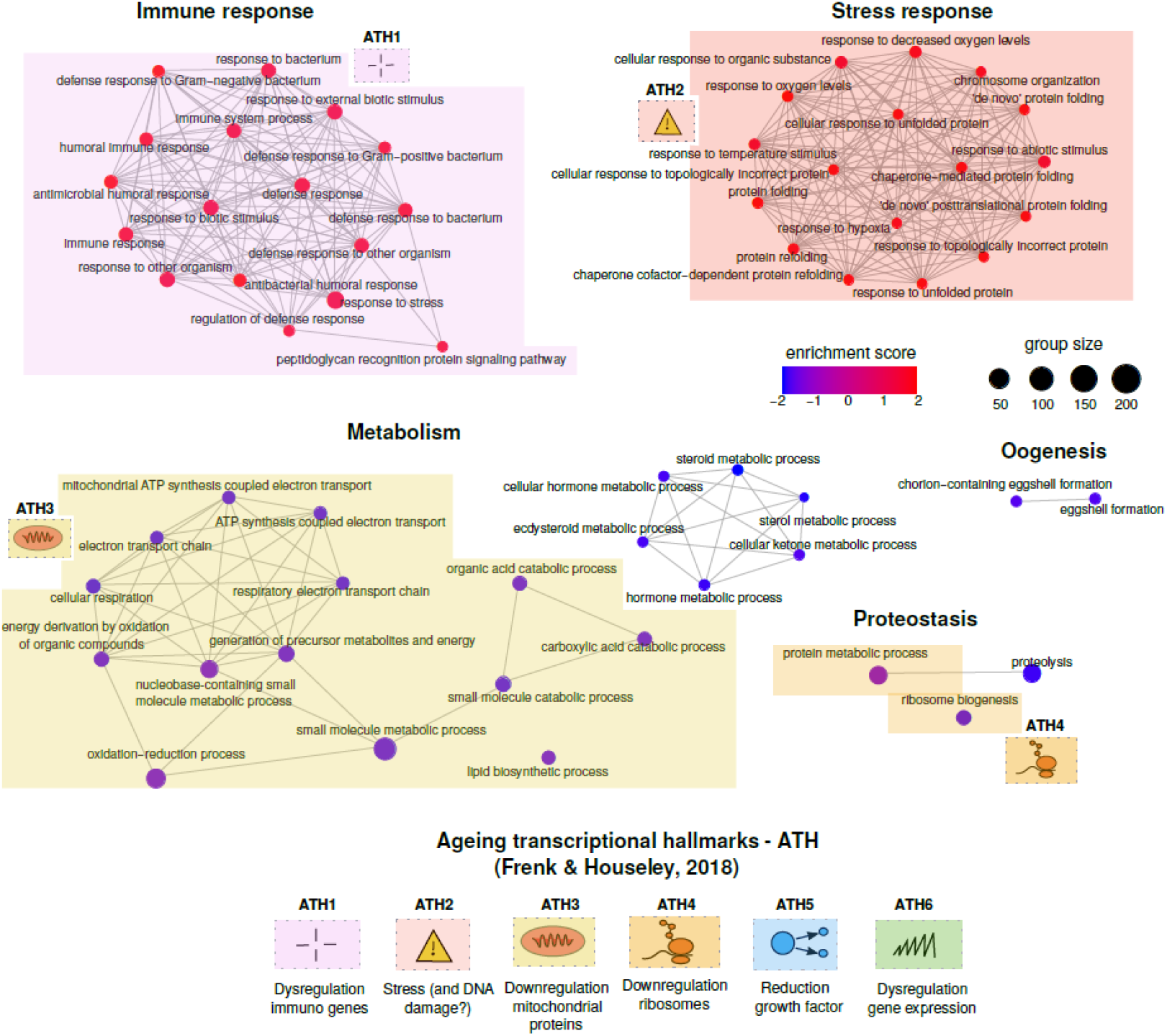
GSEA analysis (GO biological process categories) of Smurf specific genes. GSEA results are represented as a network, where nodes are significantly enriched categories (deregulation colour code as in legend) and edges are connected categories with overlapping genes. From the 59 significant categories, we identified and manually annotated five hubs: immune response, stress response, metabolism, proteostasis and oogenesis. Hallmarks of transcriptional ageing, as enunciated in Frenk & Houseley^29^ (bottom of figure). The hallmarks present in the Smurf specific signature (ATH1-4) are mapped close to the related categories. Overall, in the Smurfs specific genes we detect four hallmarks of transcriptional ageing. Note that the DNA damage response (ATH4) is indicated with a question mark in Fig. 2 following the conflicting data presented by Frenk & Houseley. No category maps to ATH5 (reduction in growth factor, downregulation of cell cycle genes) and ATH6 (increased transcriptional hetereogeneity, DNA and RNA dysregulation).

Downregulated genes show a broad enrichment in metabolism-related categories (ATH3). The decreased expression of genes involved in fatty acid biosynthesis, such as *FASN1* (log_2_FC=-0.61), *ACC* (log_2_FC=-0.31) and *eloF* (log_2_FC=-0.41) corroborates the decreased triglycerides content previously described in Smurfs^19^. The mitochondrial electron transport chain (ETC) also shows a broad downregulation (ATH3). In order to provide a quantification of the ETC downregulation, we mapped the Smurf DEGs on ETC complexes Flybase annotation, and computed the percentage of downregulated genes. Through all the complexes, all the genes detected as DEGs are downregulated (no upregulation observed) (Complex I: 17 genes, 38% of the Complex I, average log2FC = -0.18; Complex II: 2 genes, 33% percent of Complex II, average log2FC = -0.17; Complex III: 4 genes, 29% of the Complex III, average log2FC = -0.21; Complex IV: 4 genes, 19% of Complex IV, average log2FC = -0.18; Complex V: 7 genes, 41% of Complex V, average log2FC = -0.19. Percentage refers to the number of genes detected in our dataset for the specific complex). Despite the minor fold changes, the ETC components’ persistent downregulation may indicate that the aerobic metabolism they mediate is also downregulated. In addition, the upregulation of lactate dehydrogenase gene (*Ldh*) (log_2_FC=0.95) could suggest a compensatory anaerobic metabolism replacing a probable dysfunction of the aerobic ETC path, or an altered pyruvate intake into the mitochondria. Consistently, *Idh3A*, *Idh3B*, *Mdh1*, *Mdh2* and *Fum1*, involved in the tricarboxylic acid (TCA) cycle are downregulated, with fold changes similar to the ones reported above.

Genes involved in ecdysone biosynthesis (*sad*, *spo* and *phm*) and egg formation (*Vm26Aa*, *Vm26Ab*, *Vml* and *psd* are downregulated (log_2_FC is respectively -2.67, -2.63, -2.51, -2.49), giving a molecular hint for explaining the previously reported decrease in fertility in Smurf females and males^48^. A few categories related to proteostasis are also present amongst the ones deregulated in Smurfs. The ribosome biogenesis category (GO:0042254), mapping to ATH4, contains 190 genes out of which 46 are significantly deregulated, most of them, 96%, being downregulated. Regarding the proteolysis category, we detected the downregulation of 10 trypsin-like endopeptidases and 14 Jonah genes (serine endopeptidases family).

The Smurf signal overlaps with numerous changes that were described so far as ageing-related, mapping to four out of six ATH (ATH 1-4).

We compared our results with proteomic and metabolomic data obtained from Smurf and non-Smurf mated females from the same genetic background. Enrichment analysis on significantly differentially represented proteins (ANOVA p-value < 0.05, for complete results see Supplementary File 2) confirms our results of a decreased fatty acid catabolism, mitochondrial respiration and ribosomal proteins (Fig. S6). Response to stress (including genes such as *cact, Hsp70* and *Cat*) is upregulated, in line with what described in our transcriptome study.

Quantitative enrichment analysis on metabolites concentrations in Smurfs and non-Smurfs (Supplementary file 3) confirms the molecular separation between the two phases (Fig. S7) and the metabolic transcriptional signature observed. We detected deregulation of fatty acid biosynthesis and degradation pathways (KEGG^49^, with palmitic acid [log_2_FC=-1.37] and myristic acid [log_2_FC=-1.69], Fig. S8) and pyruvate metabolism (which includes metabolites from the TCA cycle) (Table S2). Regarding glucose metabolism, the overexpression of *Ldh* is confirmed by a significant (Wilcoxon test, p-value < 0.05) lactic acid increase in Smurfs (log_2_FC=0.90) (Fig. S9). The TCA cycle displays a significant general decrease at a transcriptomic level, and a general impairment at a metabolomic level,though the only metabolite significant to Wilcoxon test is succinate, log_2_FC=1.28) (Fig. S10).

These results indicate that the transcriptional dysregulation observed in Smurfs has a functional impact.

### Old Smurfs carry additional age-related changes

Our analysis (Fig. 1a, Fig. S3 and S4) suggested transcriptional differences between the old and young Smurfs. We therefore applied a DEG analysis restricted to Smurfs. Only 4 DEGs were identified when comparing 20 and 30-day Smurfs (FDR cut-off at 5%) while the 40 days Smurfs present 2320 DEGs compared to 20-day Smurfs (1385 upregulated and 935 downregulated) (DESeq2 results in Supplementary File 4). GSEA identified 125 deregulated GO BP categories (Fig. 3 and Table S3). The majority of the detected categories are associated with RNA processing, transcription, chromatin organisation, DNA replication and repair (ATH6). In the case of old Smurfs, we find downregulation of genes involved in histone methylation (*trr, Cfp1, Dpy-30L1, Smyd5, NSD, CoRest, Lpt*, average log_2_FC∼-0.26), amongst which genes of the Polycomb Repressive Complex 2 (*esc, E(z), Su(z)12I,* average log_2_FC∼-0.24). We also detect the downregulation of the histone deacetylase *HDAC1* (log_2_FC=-0.18) and genes involved in histone acetylation (as *CG12316, Ing3, Ing5, Taf1, Atac3, Brd8, Spt20, mof,* average log_2_FC∼-0.30). Chromatin-related genes are thus modestly (0 < |log_2_FC| < 1) but broadly decreased in old Smurfs. Interestingly, our proteome analysis shows a significant decrease of H3.3B (log_2_FC=-0.43) and H4 (log_2_FC=-0.54) in Smurfs suggesting a “loss of heterochromatin”^50^. Another interesting signal is the DNA repair nodes (“GO:0006302 double-strand break repair”, “GO:0006281 DNA repair”), where we retrieve 12% of the detected genes as significantly downregulated (average log_2_FC=-0.24). We also retrieved nodes associated with downregulation of genes involved in cell cycle (as cyclins), or their regulators (as *E2f2*, log_2_FC∼-0.17), which map to the ATH5 (growth factor and regulation of cell cycle). Genes involved in spindle organisation during mitosis are also found downregulated (as *Mtor* - log_2_FC ∼ -0.28- and *Chro* - log_2_FC ∼ -0.19-) suggesting a broad dysregulation of cell proliferation processes.

**Figure 3.**
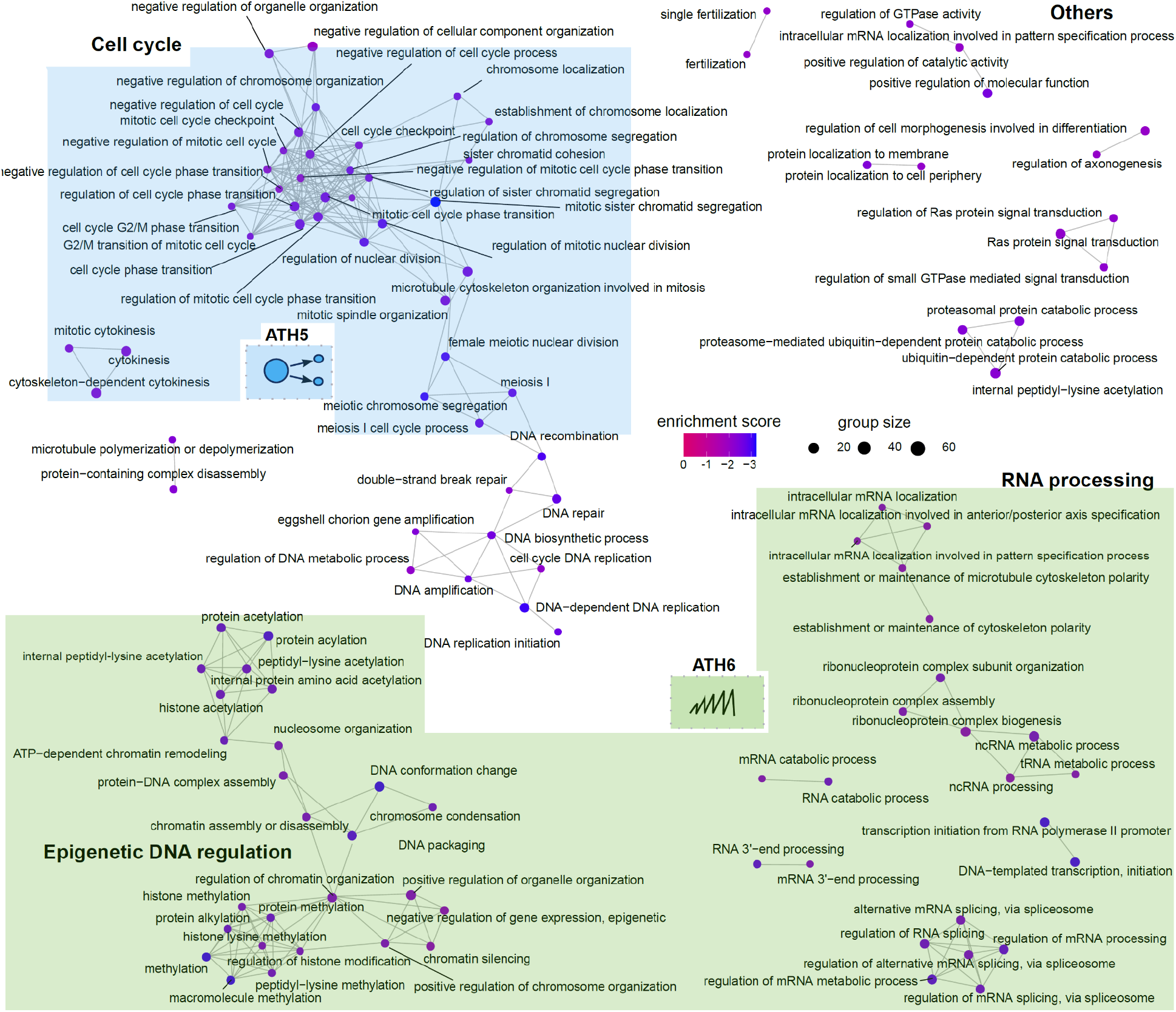
Old Smurfs carry an ageing-related signal amongst downregulated genes. Results of the GSEA analysis are represented as in Fig 3. Only downregulated nodes presenting at least one interconnection are represented here. Complete list of deregulated categories can be found in Supplementary Table S8. GSEA analysis identifies 115 downregulated GO BP categories, which are mostly related to DNA regulation, RNA processing and cell cycle regulation. A few nodes are associated with DNA repair. Interestingly, the signal carried by the old Smurfs maps (at least partially) to the “dysregulation in gene expression” (in green, ATH6) and the “reduction in growth factors” (ATH5) transcriptional ageing markers that were not detected in the Smurf specific signature. In addition, the DNA damage nodes show downregulation of genes involved in DNA repair, which has also been discussed as an ageing marker. Interestingly, there are no hubs in the network overlapping with the Smurf specific signature of Fig.2, showing that the core Smurf signal is not affected by chronological age. However, the old Smurfs do carry an additional signature compared to their younger counterparts, suggesting the existence of a “chronological-age burden” that might increase the probability of entering the Smurf pre-death phase, without however being necessary or sufficient for it.

The old Smurf signature therefore partially carries ATH5 and ATH6, the two hallmarks of transcriptional ageing that we did not detect in the Smurf specific signature. It is important to highlight that we do not find Smurf-related categories in the GSEA output, confirming that young Smurf and old Smurfs indeed do carry the same Smurf signature illustrated in Fig. 2. However, our analysis shows that the old Smurfs carry additional transcriptional changes, which mostly relate to transcription and DNA regulation. To investigate if those are time-dependent changes, which are weakly carried by old individuals and then enhanced in the Smurf stage of their life, we fitted a per-gene regression model on all samples, including as explanatory variables Smurfness, time and an interaction term amongst the two. We then performed GSEA on the list of genes presenting significant coefficients (F-statistic). The RNA processing categories (as well as the “chromosome organization”) were detected as significantly affected by time, suggesting that the deregulation trends for such processes may already be present in the non-Smurfs.

### Removing the Smurf-specific signature unveils the transcriptional effects of chronological age

In order to confirm the Smurf-specificity of the signature described above, we removed Smurf samples from the study and compared the non-Smurfs over time. Only 526 DEGs were found when comparing 20 and 40 days old non-Smurfs (and 57 when comparing 20 and 30 days old non-Smurfs) (DESeq2 results in Supplementary File 5). 59% of these genes are overlapping with Smurf-specific DEGs. 22 GO BP deregulated categories were found by GSEA (Fig. 4a and Table S4). Overall, the genes that are known as being downregulated with ageing are actually downregulated mostly in Smurfs (Fig 5b, point i), with little to no effect associated with chronological age (Fig. 4b, point ii). The largest overlap is observed for the immune response pathways (ATH1, increased inflammation). Out of the overlapping genes (20), 50% are AMPs, produced downstream the pathway. We do not find significant deregulation of the *dl* transcription factor (Smurf significant log_2_FC=0.27), while *rel* is upregulated (log_2_FC=0.42, while for the Smurfs we detected a log_2_FC of 0.61). These results suggest that the immune response is active in the old non-Smurf but to a lower extent than in Smurfs.

**Figure 4.**
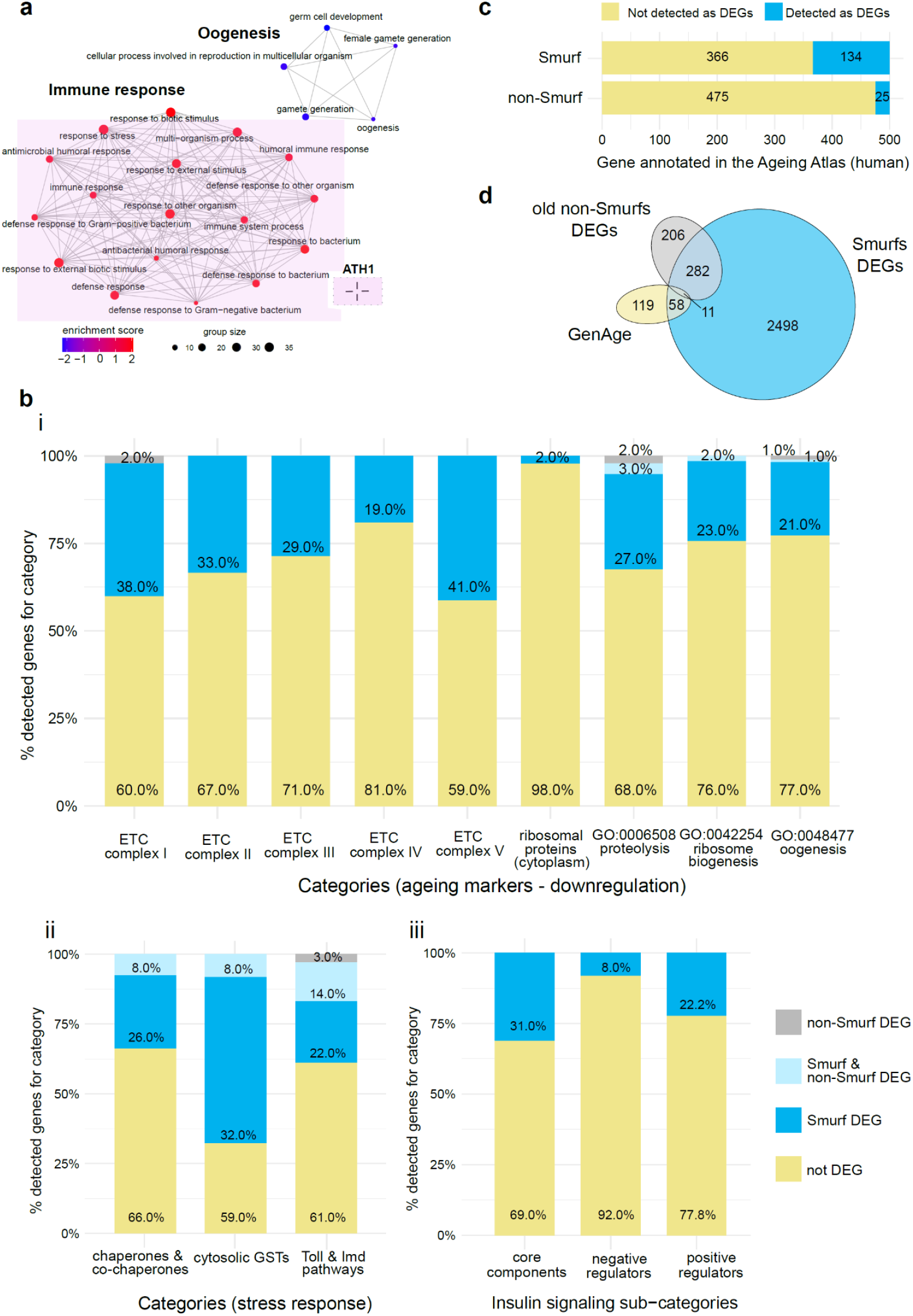
Smurfness is a better predictor of transcriptional ageing markers than chronological age. **a) GSEA analysis (GO BP categories) on old non-Smurf specific genes.** Results are represented as in Fig. 2. GSEA analysis identifies 22 deregulated GO BP categories, related to immune response (upregulation, in red) and oogenesis (downregulation in blue). The analysis carried on chronological age can therefore detect only one hallmark of transcriptional ageing^29^ (ATH1, for representation of transcriptional hallmarks, see Fig. 2). **b) Manual mapping of Smurf and old non-Smurf DEGs on ageing processes.** For each process, the histograms represent the percentage of genes mapping to it but not detected as DEGs in our analysis (yellow), detected as Smurf DEGs (blue), detected as both Smurf and non-Smurf DEGs (light blue), or only detected in the old non-Smurf DEGs (grey). When not stated otherwise, the gene lists are retrieved from Flybase. Genes described as downregulated with ageing **(i)** are mostly detected only in Smurfs, with the exception of structural ribosomal proteins, whose downregulation is not significant in Smurfs. For the processes described as upregulated with ageing **(ii)**, the Smurf samples do retrieve more information than the non-Smurfs, with the last however carrying more signal than in the case of the downregulated genes, especially for the immune response (as already showed in **(a)**). Similarly, the IIS pathway displays deregulation in the Smurfs, while no gene is detected as deregulated when looking only at chronological age **(iii)**. **c) Mapping of Smurf and non-Smurf DEGs to human ageing-related genes (annotated in the Ageing Atlas).** The Ageing Atlas annotates 500 human ageing-related genes. All of those have orthologs in Drosophila, which are all present in our dataset. By studying the Smurf phenotype, we can detect 134 genes out of the annotated 500. The number of detected genes drops to 25 when using chronological age only as an ageing marker. **d) Longevity genes and Smurfness.** The Venn diagram shows the overlap between the annotated longevity genes in Drosophila (GenAge), the Smurf DEGs and the non-Smurf DEGs. While Smurf-centred analysis retrieves ∼37% of the longevity genes, the non-Smurf centred analysis only retrieves ∼6%, not adding information to what was already detected by the Smurf analysis.

**Figure 5.**
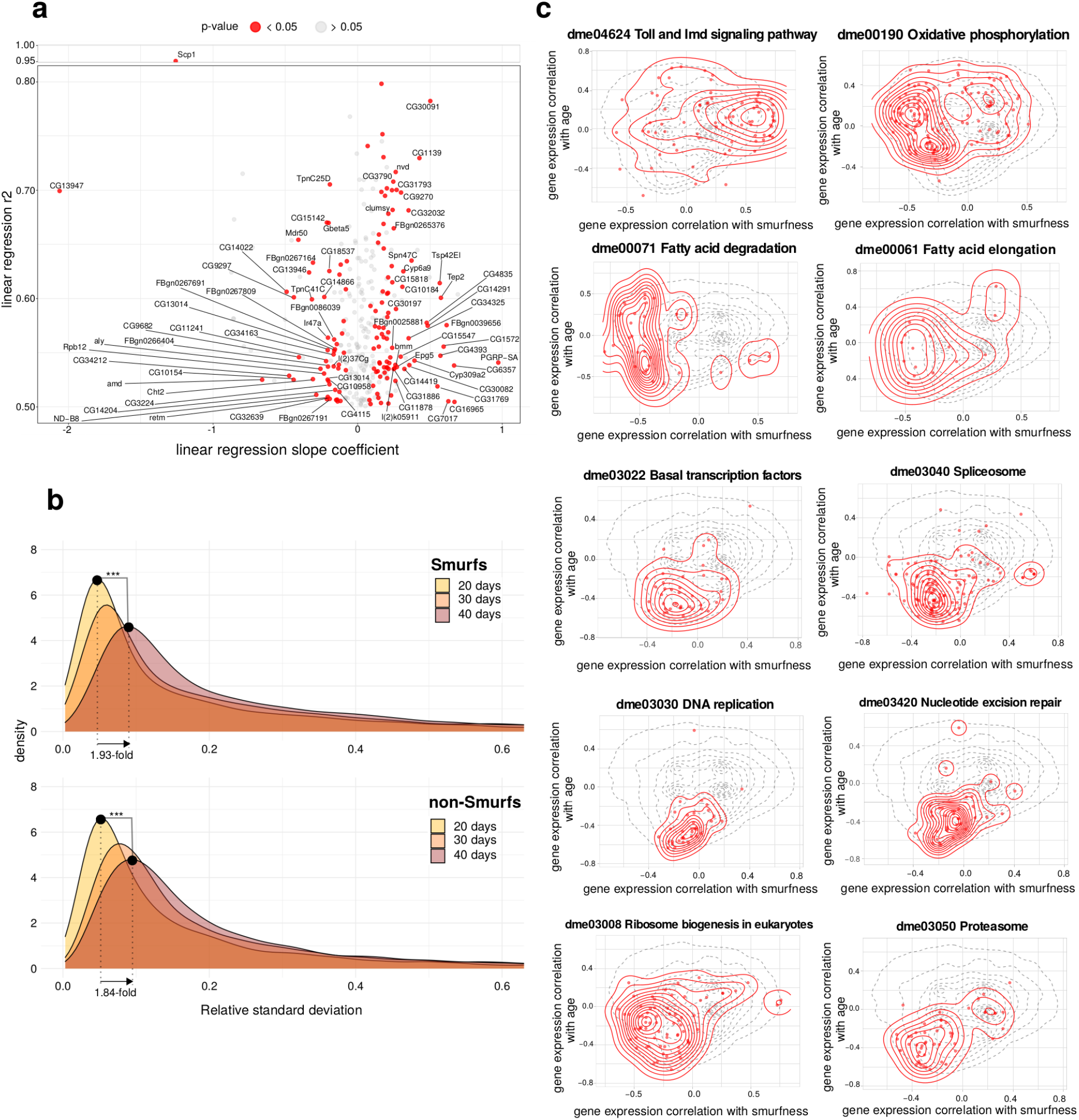
Chronological age and Smurfness respective effects on the transcriptome. **a) Linear regression of gene expression in non-Smurfs over time.** The r^2^ of the applied linear model is plotted as a function of the slope coefficient. Only genes non differentially expressed in Smurfs are plotted, in order to focus on a possible weak age-related non-Smurf signal. Genes presenting a significant slope are plotted in red. **b) Chronological age effect on transcriptional heterogeneity.** The RSD densities are plotted for the different ages group (Smurf and non-Smurf). The tail of the distribution is cut at RSD = 0.6 for illustration purposes. Smurfs and non-Smurfs present a similar behaviour, with the peak of the distribution showing an almost 2-fold increase from 20 days to 40 days (peak_S20_ = 0.046, peak_S40_ = 0.089, peak_NS20_ = 0.051, peak_NS40_ = 0.094), showing the effect of chronological age on transcriptional noise. ***p-value < 10^-16^ (KS statistic). **c) Effect of Smurfness and chronological age on biological pathways.** Smurfness and chronological age both affect the biology of the individual. Here we show how some pathways are affected by age and Smurfness respectively. Dotted line in the background corresponds to the density of all the genes analysed. Red points and density correspond to the genes mapping to the pathway (KEGG database) of interest. The statistics was assessed using the Fasano-Franceschini test (FDR adjusted p-value). Toll and Imd pathways (r_smurf_ = 0.248, r_age_ = 0.080, p-value = 5.2e-06), oxidative phosphorylation (ETC genes, r_smurf_ = -0.217, r_age_ = 0.088, p-value = 4.5e-15), fatty acid degradation (r_smurf_ = -0.388, r_age_ = -0.063, p-value = 4.3e-09) and fatty acid elongation (r_smurf_ = -0.255, r_age_ = -0.031, p-value = 3.8e-03) are mostly correlating with smurfness; spliceosome (r_smurf_ = - 0.124, r_age_ = -0.288, p-value = 1.5e-17), basal transcription factors (r_smurf_ = -0.096, r_age_ = -0.318, p-value = 3.1e-08), DNA replication (r_smurf_ = -0.070, r_age_ = -0.393, p-value = 2.2e-09) and repair (Nucleotide excision repair, r_smurf_ = -0.073, r_age_ = -0.338, p-value = 1.2e-10) are mostly correlating with age; Ribosome biogenesis (r_smurf_ = -0.203, r_age_ = -0.159, p-value = 4.0e-10) and proteasome (r_smurf_ = -0.166, r_age_ = -0.276, p-value = 3.5e-09) apper to occupy a zone of similar correlation with both Smurfness and age (with the peak of the density for the ribosomial pathway occupying a zone of high correlation with Smurfness, as expected given the results obtain in our analysis -Fig. 2 and Fig. 3-).

Regarding the genes mapping to the insulin-like receptor signalling (IIS) pathway (Fig 5b, point iii), we do not detect any deregulation in the non-Smurfs, with IIS core components being affected only in Smurfs. While no significant change is detected for the *Ilp* genes (insulin-like peptides activating the pathway), we find low but significant upregulation of *Inr* (receptor, log_2_FC=0.42), *chico* (first kinase of the cascade, log_2_FC 0.23) and the kinase *Akt1* (log_2_FC=0.18). *Inr* and *chico* are well-described longevity genes in Drosophila, positively affecting ageing when negatively modulated^51, 52^. No significant changes are detected for the Drosophila mTOR genes *Tor* and *raptor,* nor *foxo*. However, we find significant upregulation of *Thor*, coding for the homologous mammalian translation initiation factor 4E-BP, a *foxo* target of which the upregulation was already described at the protein level in Smurfs^19^.

Our dataset contains all the orthologs of the 500 human genes associated with ageing present in the Ageing Atlas^53^ (Table S5 and S6). We find that 26.8% of these genes are present in the Smurf list (121 Drosophila genes corresponding to 134 human genes), while only 4% are present in the old non-Smurfs (24 Drosophila genes corresponding to 25 human genes) (Fig. 4c).

Over the past 40 years, numerous genes have been shown to modulate ageing when artificially deregulated. We explored whether our list of DEGs is overlapping these “longevity genes”. Out of the 201 Drosophila longevity genes annotated in GenAge^54^, 188 are present in our dataset. Smurfs DEGs allow the detection of 37% of them, while the old non-Smurf DEGs detect only 6% (Fig. 4d and Tables S7 and S8). Furthermore, all the longevity genes present in the non-Smurf DEGs are also present in the Smurf DEGs.

Taken together, the results show that Smurfness predicts ageing-associated changes described in the literature better than chronological age.

### Identifying weak chronological age-dependent signature

In light of the evidence that most of the transcriptional alterations described as age-related are Smurf-specific, with only a small part of the signal retrieved in old non-Smurfs (Fig. 4), we wondered whether weaker but relevant age-related changes might be present in non-Smurfs but missed by the DESeq2 approach. We therefore regressed gene expression data on chronological age (20, 30, 40 days) in the non-Smurfs using a linear model. After filtering for significance to F-test (p-value < 0.05) and R^2^ (> 0.5) we identified 301 genes (207 showing an increasing expression with time, 94 decreasing) (Table S9). 51.6% of these genes also belong to the Smurf DEGs. We focused on the 146 remaining genes (93 with positive slope, 53 negative). Results are presented in Fig. 5a. No enrichment in GO categories was found (GOrilla enrichment^55^, using the whole set of detected genes as background), suggesting that once the Smurf signal is removed, no strong coherent deregulation can be detected in the non-Smurfs in our dataset. Nevertheless, figure 2a shows the old non-Smurf samples to cluster with old Smurf samples. This is supported by the decreasing number of detected DEGs between age-matched Smurf and non-Smurfs with chronological age (Fig. S11).

Ageing has been reported as increasing the gene expression heterogeneity in a variety of organisms, tissues and cell types^56–63^ (ATH6). We computed the relative standard deviation (RSD) of each gene for each group (Smurfness and age), plotted the distributions of the RSD across groups and compared them using the Kolmogrov-Smirnov (KS) statistic (Fig. 5b). All genes are affected, independently of their expression levels (Figure S12). In both Smurfs and non-Smurfs, the peak of the RSD distribution shifts towards the right with age (1.93-fold increase for the Smurfs, and 1.84-fold for the non-Smurfs) suggesting that gene expression increases in heterogeneity as a function of chronological age with no further changes at the Smurf transition.

In brief, our results show that four out of six transcriptional ageing markers (ATH1-4) are specific to the Smurf phenotype, independently of their chronological age (Fig. 2). On the other hand, the alteration in chromatin-related genes and mRNA processing, as well as cell cycle genes (together with a weaker DNA repair signal) appear to be exclusively carried by the old Smurfs (ATH 5-6) (Fig. 3). We could not identify biological processes strictly related to the old non-Smurfs compared to their young counterparts (Fig. 4). However, the increased heterogeneity in gene expression (ATH6) appears to be primarily affected by chronological age (Fig. 5b). In order to visually represent the relative effect of both the chronological and biological age we computed the correlation of individual gene expression with each.We identified 113 annotated KEGG pathways where at least 10 genes present in our dataset are mapped. We finally obtained 48 correlating (Fasano-Franceschini test^64^, FDR for p-value correction) with Smurfness (Table S10) and 38 correlating with chronological age (Table S11). Fig. 5c shows the Toll and Imd pathways mostly displaying positive correlation with Smurfness; the ETC (oxidative phosphorylation pathway) and fatty acid degradation/elongation mostly negatively correlates with Smurfness, while showing a lower correlation with age. Interestingly, transcription-related pathways (spliceosome and basal transcription factors) as well as DNA amplification and repair pathways show a higher negative correlation to chronological age compared to Smurfness. Finally, the proteasome and ribosome biogenesis seem equally affected by chronological age and Smurfness.

### Using Smurfness to identify new “longevity genes”

We decided to investigate whether altered expression of transcription factors (TFs) could explain the transcriptional signature of Smurfs. We identified 102 TFs showing altered expression in Smurfs (77 upregulated, 25 downregulated, Table S12) out of the 629 annotated in Flybase. In order to reduce the potential functional redundancy in this list, we used i-cisTartget^65, 66^ to predict putative upstream regulators of the Smurf-deregulated TFs. We selected the hits presenting a score above 4 (3 being the recommended minimum threshold). Second, to avoid limiting our selection criteria only to TFs, we applied the same i-cisTarget algorithm to genes showing at least a 4-fold difference (|log_2_FC|>2). Results are shown in Table S13. We selected 17 TFs of interest for functional validation amongst the best i-cisTarget scores or high deregulation (Table 1).

**Table 1.** List of TFs selected for experimental validation. 17 TFs were selected for functional validation: 8 were found in the i-cisTarget analysis, 3 in both DESeq2 and i-cisTarget analysis and 6 in the DESeq2 analysis alone, chosen for their strong deregulation.

| Gene symbol | Selection method | Deregulation |
| --- | --- | --- |
| *Adf1* | i-cisTarget | putative regulator TFs up in Smurf |
| *Aef1* | i-cisTarget | putative regulator TFs up in Smurf |
| *CG4360* | i-cisTarget | putative regulator TFs up in Smurf |
| *FoxP* | DESeq2 & i-cisTarget | up in Smurf & putative regulator TFs up in Smurf |
| *Hsf* | i-cisTarget | putative regulator genes up in Smurf |
| *Trl* | i-cisTarget | putative regulator TFs up in Smurf |
| *dmrt93B* | DESeq2 | up in Smurf |
| *Ets21C* | DESeq2 | up in Smurf |
| *Hey* | DESeq2 | up in Smurf |
| *kay* | DESeq2 | up in Smurf |
| *Mef2* | DESeq2 & i-cisTarget | up in Smurf & putative regulator TFs up in Smurf |
| *rib* | DESeq2 | up in Smurf |
| *Ets96B* | DESeq2 | down in Smurf |
| *GATAd* | i-cisTarget | putative regulator TFs down in Smurf |
| *GATAe* | i-cisTarget | putative regulator TFs down in Smurf |
| *NF-YB* | DESeq2 & i-cisTarget | up in Smurf & putative regulator TFs up in Smurf |
| *srp* | i-cisTarget | putative regulator TFs down in Smurf |

To assess their effect on mean lifespan (ML), we proceeded with their knock-down (KD) and/or overexpression (OX) using GeneSwitch^67, 68^ (GS). This technique, widely used in Drosophila, allows spatially and temporally tuned KD or OX in individuals of the same genetic background. Since our candidate genes were selected from whole body data, we used the ubiquitous daughterless-GS (*da*GS) driver. When transgenic lines were available we performed both KD and OX during the adulthood of the fly (i.e. after eclosion) or during its whole life (development and adulthood)(Fig. 6a). Five different concentrations of RU486 (0 µg/mL -control, 10 µg/mL, 50 µg/mL, 100 µg/mL, 200 µg/mL) were used to explore a broad range of inducing conditions, as in ref ^69^. During development, we lowered the concentrations by a factor 10 in order to avoid potential toxic effects, as suggested by Osterwalder et al.^67^ and performed in Rera et al.^70^.

**Figure 6.**
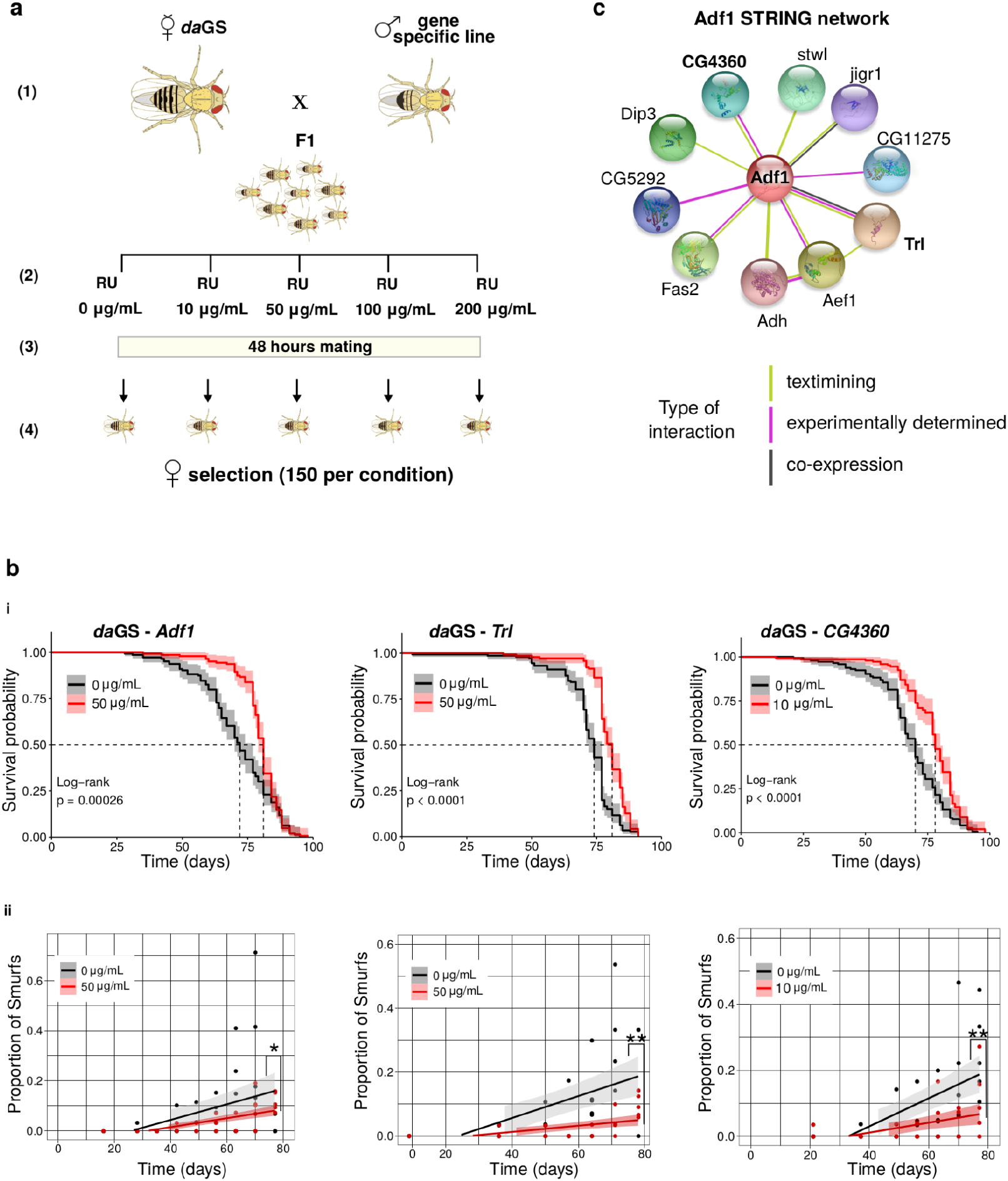
Identification of new longevity genes using the Smurf phenotype. **a) Gene expression alteration through GeneSwitch (GS).** KD and/or overexpression of the target gene in the whole body of Drosophila were performed by crossing virgins females of the ubiquitous *daughterless*-GS (*da*GS) driver with males carrying the UAS transgene (Step 1). The F1 was reared either on food without the inducer RU486 (adult only induction), either with food presenting the following RU486 gradient: 0 µg/mL -control-, 1 µg/mL, 5 µg/mL, 10 µg/mL, 20 µg/mL (whole life induction). At the moment of eclosion, flies are transferred onto food with the following RU486 concentrations: 0 µg/mL -control-, 10 µg/mL, 50 µg/mL, 100 µg/mL, 200 µg/mL. Flies are randomly distributed if not developed on drug, otherwise they are distributed according to the developmental drug condition (Step 2). Flies are left mating for 48 hours (Step 3) and subsequently 150 females per concentration (divided on 5 vials/30 females each) are randomly selected for the longevity experiment (Step 4). **b) Effect of *Adf1*, *Trl* and *CG4360* KD on longevity and on the Smurf dynamics in the population. (i)** The KD of *Adf1* (+11.8%, ML_RU0_ = 71.0, ML_RU50_ = 79.5) and *Trl* (+10.5%, ML_RU0_ = 72.2, ML_RU50_ = 79.8) in the whole body during adulthood significantly extend lifespan, as well as for the KD during the whole life of *CG4360* (+12.4%, ML_RU0_ = 68.5, ML_RU10_ = 77.0). **(ii)** The proportion of Smurfs for the corresponding control and treated populations are plotted as a function of time. The proportion of Smurfs is computed as the number of Smurfs over the total number of flies alive (Smurfs + non-Smurfs). Data are fitted using a linear approximation^19, 20^. In all cases, the populations show a significant increase with time of the Smurf proportion (F-statistic) (*Adf1*: slope_RU0_ = 0.0055, p-value_RU0_ = 4.72e-03, slope_RU50_ = 0.0018, p-value_RU50_ = 4.17e-07; *Trl*: slope_RU0_ = 0.0044, p-value_RU0_ = 6.53e-04, slope_RU50_ = 0.0009, p-value_RU50_ = 5.39e-04; *CG4360*: slope_RU0_ = 0.0042, p-value_RU0_ = 6.58e-05, slope_RU10_ = 0.0015, p-value_RU10_ = 6.51e-03). Furthermore, the slope of the control population is significantly different from the one of the treated (F-statistic), which displays a slower increase in the Smurf proportion with time. P-values indicated in figure: * < 0.05; ** < 0.01. **(c) Adf1 interaction network from STRING database.** The three TFs identified as new longevity genes have been retrieved from i-cisTarget as putative regulators of upregulated Smurf TFs. The annotated interactions in the STRING database show how those genes have been already described together. Adf1 and Trl displayed stronger evidence (text mining, co-expression and proved interaction in Drosophila *in vitro*), while the evidence for CG4360 and Adf1 interaction comes from text mining and interaction shown between homologous in *C. elegans).* We decided to assign to CG4360 the gene name of *Sag1* (Smurf Associated Gene 1) given its potential involvement in the Smurf phase. Regarding the remaining nodes of the network, they show weaker evidence (see Fig. S19). CG11275 and CG5292 have been shown to interact with Adf1 on two-yeast hybrid assay on the FlyBI project (https://flybi.hms.harvard.edu/).

The longevity experiments are summarised in Fig. S13 and Table S14. Four TFs presented a positive effect on ML when knocked-down in at least one RU486 condition during adulthood *Trl* + 9.5%, *Adf1* +7.6%, *CG4360* +7.3%, *Ets96B* +6.6%) and one when overexpressed during adulthood and development (*Hsf* +10.3%). A second independent experiment confirmed the effect of *Trl*, *Adf1*, *CG4360* (Fig. 6b, point i). A third experiment validated the ML extension of CG4360 when downregulated during adulthood only (Fig. S14), as the first two experiments showed contrasting results for the longevity effect when downregulation was performed during the whole life. We confirmed the knockdowns through RT-qPCR for each line (Fig. S15), and validated that RU486 alone has no effect on ML (Fig. S16).

We then tested whether the identified ML extension was due to delayed entry in Smurf state (Fig. 6b, point ii) by fitting two linear regression models. First, in order to test the effect of chronological age, we regressed the proportion of Smurfs on chronological age separately in both the controls and KD individuals. Secondly, in order to investigate the difference between the two populations, we regressed the proportion of Smurfs on chronological age and RU486 concentration (as a categorical variable), allowing for an interaction between chronological age and RU486 concentration.

The results show that the conditions leading to ML extension also lead to a slower increase in Smurf’s prevalence (Fig. 6b, point ii). This was not the case for conditions not leading to different ML (Fig. S17). These results suggest that the KD of the studied genes increases the mean lifespan by extending the non-Smurf period of life, possibly because these genes modulate early steps of ageing. Interestingly, the three genes we validated for their role in longevity are reported to possibly interact based on the STRING database^71^ (Fig. 6c). We failed to identify any significant increase of mean lifespan on males (Fig. S18).

## Discussion

This study describes how the long defined transcriptional signature of ageing and associated “ageing transcriptional hallmarks”, instead of accompanying chronological age with continuous and progressive changes, actually behave in a biphasic manner. We identified this previously hidden behaviour thanks to the Smurf, two-phase, model of ageing.

The detection of living individuals showing an increased intestinal permeability to a very small (800Da), non-toxic, blue food dye previously allowed us to propose a model of ageing with two consecutive and necessary phases. Although a recent article by Bitner and colleagues^72^ suggest that only a low proportion of flies undergo the Smurf transition, our extensive characterization of the phenotype using female flies from lines of different genetic backgrounds characterized by significantly different life expectancies ranging from 20 to 80 days (DGRP, *Drs*GFP, w^1118^, Canton-S, Oregon-R and w^Dahomey^) as well as in F1 individuals, monitored individually or in groups, unequivocally show that every female Drosophila dies as a Smurf. In addition, the Smurf phenotype also accompanies ageing in Drosophila males as shown by us and others. Failure by Bitner et al. to reproduce our results is likely due to their non-standard protocol.

The mathematical model we developed^20^ fitted reasonably well survival curves while allowing for a direct interpretation of the parameters. In addition, the hypothesis that every individual turns Smurf prior to death allowed us to predict the relatively constant remaining lifespan of Smurfs irrespective of their chronological age, which we then validated using multiple DGRP lines. Given our previous description of physiological hallmarks of ageing - loss of fertility, mobility, energy stores - segregating with Smurf individuals (and defining Smurfness as an objective indicator of frailty), we decided here to explore the behaviour of the transcriptional hallmarks of ageing in light of the Smurf state of individuals and their chronological age.

Distinguishing these two subpopulations allowed us to observe the gene expression noise doubling between young and old individuals, making the transcriptional noise (ATH6) the only transcriptional hallmark of ageing to display a time-dependent behaviour in our study. This increase of noise in the gene expression level, often associated with transcriptional drift^57^, is concomitant with the time-dependent increasing risk for an individual to enter the second and last phase of life, the Smurf phase. Interestingly, interventions decreasing it were already shown to extend lifespan in nematodes^73^. ^75 76^. Old non-Smurfs also show some of ATH1 suggesting that inflammation could precede the Smurf transition at least in old individuals although we cannot exclude that at an advanced age, the likelihood of sampling pre-Smurf or early Smurfs is high and this signal could be due to such individuals contaminating the non-Smurf samples. Then, individuals in the Smurf phase undergo a dramatic shift in gene expression with over 3000 genes differentially expressed compared to age-matched non-Smurf individuals. More importantly, these genes span across the six ageing transcriptional hallmarks, systemic inflammation (ATH1), active stress response (ATH2), decreased mitochondrial/energy metabolism (ATH3) and altered protein translation (ATH4). Old Smurf individuals also show a worsening of their DNA repair pathways, cell cycle regulation pathways (ATH5), chromatin regulation and RNA processing (ATH6). Recently, David Gems and João Pedro de Magalhães questioned the position of the hallmarks of ageing as a paradigm. Our results here seem to support this questioning^74^.

Indeed, the hallmarks are defined as 1) manifesting in an age-related fashion, 2) their accentuation accelerating ageing and 3) intervention on them leading to delay, reverse or stop ageing^1^. However, rather than causative of the process they appear to be markers of a terminal and, so far, non-reversible phase of life except for the dysregulation of gene expression. Further characterization of the chain of events might allow to discriminate between major theories of ageing such as “inflammageing”, “genome maintenance” or “oxidative damage”. Can an evolutionary conserved hallmark of ageing characteristic of the Smurf phase of life be a driver of ageing? On the other hand, if ageing is not programmed, how can such a late-life phase be so much evolutionarily conserved and molecularly stereotyped?

In addition, here we show how most of the pro-longevity genetic interventions identified so far involve genes affected by the Smurf transition. Our longevity experiments in Drosophila demonstrate that it is possible to significantly increase lifespan by tuning the expression of TFs likely to explain the Smurf-associated transcriptional signature (*Trl*, *Adf-1* and CG4360/*Sag1*) and delay the time of entrance in the Smurf phase. Although moderate, these increases of health and lifespan were consistent across inducing conditions and independent experiment, while of a similar extent to longevity studies properly controlling for genetic background using the gene switch system. The fact that we do not detect an increase in lifespan in males does not invalidate the results obtained on females. Those results are in line with the physiological sexual dimorphism of Drosophila longevity, an issue which has been recently more investigated^77–79^ but they could also be due to a sex-specificity of the transcriptional signature presented in this article or due to the weaker inducibility of the daGS driver^69^.

Even though the aim of the paper was not to characterise the events occurring at the intestinal level - the intestinal permeability is merely a marker of the last phase of life in our model - we detected alterations of cell junction components RNA as well as the JAK/STAT pathway and ECM remodelling proteins, suggesting a broad restructuring of tissues at the scale of the whole organism. This is reminiscent of the overall alteration of controlled epithelial permeability broadly affecting living organisms during ageing^80^. The recent demonstration that the Smurf phenotype is due to increased intestinal permeability but also to decreased Malpighian tubules activity^81^ is supportive of organismal functional failure occurring in the Smurf phase. Whether it is what is called multivisceral failure in humans is under investigation, but it might highlight the use of the Smurf model of ageing for the study of other barriers, especially the BBB.

By questioning the place of the hallmarks of ageing within the ageing process, our study highlights the high relevance of using the Smurf phenotype in ageing studies across multiple model organisms thanks to its strong evolutionary conservation. The absence of Smurf classification in the experimental design indeed results in a non-negligible confounding factor altering the interpretability of the results. Taking into consideration the Smurf phenotype in ageing studies is key to taking into account the interindividual heterogeneity. As schematized in our graphical abstract, looking at age-related phenotypes without the Smurf phenotype can lead to misinterpretation, attributing to advancing age what is actually due to an increased proportion of Smurf individuals. Based on our results, we anticipate that the Smurf phenotype will become a standard parameter in ageing research, not as a measurement of intestinal permeability but rather as a marker for frail individuals in the last phase of their life. Its broad evolutionary conservation as well as the distinct molecular changes occurring in the two phases of ageing will certainly allow a deep reexamination of the evolutionary mechanisms at stake in the wide presence of ageing through living organisms.

## Supplementary materials

All codes and associated processed data are available at https://github.com/MichaelRera/SmurfsTrsc Raw RNAseq data are available at NCBI Geo https://www.ncbi.nlm.nih.gov/geo/query/acc.cgi?acc=GSE219063

Supplementary figures are available at Supplementary_figures.pdf

Supplementary tables are available at Supplementary_tables.pdf

Supplementary files:

- Supplementary file 1: SupFile1_res_DESeq2_Smurf_nonSmurf.xlsx, DESeq2 results Smurf vs non-Smurf;
- Supplementary file 2: SupFile2_Proteomic_results_Smurf_nonSmurf.xlsx, proteome analysis results;
- Supplementary file 3: SupFile3_Metabolomic_data_Smurf_nonSmurf.xlsx, metabolomic processed data, used as an input for MetaboAnalyst;
- Supplementary file 4: SupFile4_res_DESeq2_40daysS_20daysS.xlsx, DESeq2 results 40 days Smurf vs 20 days Smurf;
- Supplementary file 5: SupFile5_res_DESeq2_40daysNS_20daysNS.xlsx, DESeq2 results 40 days non-Smurf vs 20 days non-Smurf

## Material and methods

### RNA-seq: experimental design

A synchronous isogenic population of *drosomycin*-GFP (*Drs*-GFP) Drosophila line was used for the RNA-sequencing experiment (40 vials of 30 mated female flies). For the longevity recording, flies were transferred on fresh food and deaths scored on alternative days. Flies were sampled for the sequencing experiment at day 20 (80% survival), day 30 (50% survival) and day 40 (10% survival). Each sample is a mixture of 8 flies. The sampling protocol for Smurfs and age-matched non-Smurfs is the following: all flies - the ones used for longevity and the ones used for sampling - are transferred on blue food overnight; at 9 a.m. 1 Smurf sample and age-matched non-Smurf are collected (Mixed samples), and all the remaining Smurfs are discharged; five hours later, 2 Smurf and non-Smurf samples are collected (5 hours Smurfs), and all the remaining Smurfs are discharged; twenty-four hours later, 3 Smurf and non-Smurf samples are collected. Note that at 90% no 5 hours Smurfs could be collected due to the low probability of flies turning Smurf at this age. After sampling, flies were immediately frozen in liquid N_2_ and stored at -80°C up to RNA extraction. Each time-point has a minimum of three biological replicates.

### RNA-seq: pre-processing

Sequencing was externalised to Intragen. Library preparation was done using ‘TruSeq Stranded mRNA Sample Prep Illumina’ kit and conducted on HiSeq4000 Illumina sequencer (paired-end sequencing). Data preprocessing was performed on Galaxy^82^ server. Quality control was performed using FastQC^83^, and resulted in no reads filtering. Reads were aligned with Hisat2^84^ on the reference *D. melanogaster* genome BDGP6.95. Reads count was performed with featureCounts^85^, resulting in a raw counts matrix of 15364 genes.

### RNA-seq: analysis

Unless stated otherwise, all analysis were performed on R 3.5.3 and plots generated with ggplot2 3.3.5. PCA was performed using package DESeq2 1.22.2. Association of components with Smurfness and age was computed using the functions PCA and dimdesc from FactoMineR 2.4. tSNE was performed on package Rtsne 0.15. Sample-to-sample distance heatmap was computed using function dist from stats 3.5.3, and plotted using heatmap 1.0.12. PCA, tSNE and clustering analyses were performed using normalized counts additionally transformed with the *vst* DESeq2 function to stabilize the variance. For the tSNE analysis, the perplexity parameter was set to 10. Additional ed details on the analyses can be found in the Github repository. The main DEGs analysis was performed on DESeq2 1.22.2, while validation analysis on edgeR 3.24.3. Enrichmend analysis was performed with the Bioconductor package clusterProfiler 3.10.1, which calls fgsea 1.8.0; analysis was ran with the following parameters: nPerm = 15000, minGSSize = 10, maxGSSize = 600. Enrichment plot was generated with the function emmaplot from the same package. Venn diagram (Fig. 4D) was generated using eulerr Rshiny app. Pearson correlation for analysis in Fig. 5C was computed with the cor() R function.

### Proteomic data collection and analysis

*Drs*GFP Smurfs (8 hours) and non-Smurfs were sampled at 80 and 10% survival in quadruplicates of 10 females. Flies were quickly homogenised in 96µL NU-PAGE 1X sample buffer containing antiproteases and quickly spun to precipitate debris. 40µL of samples were then loaded on a NU-PAGE 10% Bis-Tris gel prior to being sent for label free proteomics quantification.

### Metabolomic data collection and analysis

*Drs*GFP Smurfs and non-Smurfs were sampled at 50% survival. Each sample corresponds to a mixture of 20/30 individuals, for a total of 7 Smurf and 7 non-Smurf samples. Drosophila were weighted to reach around 30 mg in a 2 mL-homogenizer tube with ceramic beads (Hard Tissue Homogenizing CK28, 2.8 mm zirconium oxide beads; Precellys, Bertin Technologies, France). Then, 1 mL of ice-cold CH_3_OH/water (9/1, -20°C, with internal standards) was added to the homogenizer tube. Samples were homogenised (3 cycles of 20 s/ 5000 rpm; Precellys 24, Bertin Technologies) and homogenates were then centrifuged (10 min at 15000 g, 4°C). Supernatants were collected and several fractions were split to be analysed by different Liquid and Gaz chromatography coupled with mass spectrometers (LC/MS and GC/MS)^86^.Widely targeted analysis by GC-MS/MS was performed on a coupling 7890A gas chromatography (Agilent Technologies) Triple Quadrupole 7000C (Agilent Technologies) and was previously described in^87^. Polyamines, nucleotides, cofactors, bile acids and short chain fatty acids analyses were performed by LC-MS/MS with a 1260 UHPLC (Ultra-High Performance Liquid Chromatography) (Agilent Technologies) coupled to a QQQ 6410 (Agilent Technologies) and were previously described in^87^. Pseudo-targeted analysis by UHPLC-HRAM (Ultra-High Performance Liquid Chromatography – High Resolution Accurate Mass) was performed on a U3000 (Dionex) / Orbitrap q-Exactive (Thermo) coupling, previously described in^87, 88^. All targeted treated data were merged and cleaned with a dedicated R (version 4.0) package (@Github/Kroemerlab/GRMeta). 202 metabolites were detected. All the analysis presented (fold change estimation, Wilcoxon test and quantitative enrichment analysis) were done using MetaboAnlyst^89^. One Smurf sample was removed from the analysis as generated starting from 8 individuals only, resulting in a total N of 7 non-Smurfs and 6 Smurfs. Samples were normalised by weight. Gene expression and metabolites representation KEGG maps were generated using pathview 1.2^90^ (R package).

### Longevity experiments

All the flies are kept in closed vials in incubators at controlled temperature, humidity and 12 hours light cycle. Experiments are carried at 26°C. Longevity experiments (included the one from where flies were sampled for the RNAseq) were run on the following food composition: 5.14% (w/v) yeast, 2.91\% (w/v) corn, 4.28% (w/v) sugar, 0.57% (w/v) agar and Methyl 4-hydroxybenzoate (Moldex) at a final concentration of 5.3 g/L to prevent fungi contamination. Just after eclosion, flies are collected in tubes with food and RU486 (Fig. 5a). Males and females are left together to mate for 48 hours. After that time, males or females (depending on the experiment) are sorted in a number of 30 per vial, with 5 vials for each RU concentration (total N per concentration is 150). Flies are transferred to new vials with fresh food and scored three times per week (Monday, Wednesday, Friday). An exception are the first two weeks of the experiment, when females undergo an additional transfer on Saturday or Sunday due to the fertilised eggs altering the food composition. The food is prepared the day before the scoring (1.25 mL per vial) and stored at room temperature.

### Lines used

*daGS* driver (provided by Tricoire laboratory, Université de Paris). <u>Bloomington stock (with</u> <u>associated targeted gene if GS)</u>: *Drs-*GFP 55707, *dmrt93B*, 27657; *Ets21C*, 39069; *Hey*, 41650; *kay*, 27722; *Mef2*, 28699; *rib*, 50682; *Ets96B*, 31935; *GATAd*, 34625; *GATAe*, 33748; *srp*, 28606; *NF-yB*, 57254; *Aef1*, 80390; *CG4360*, 51813; *FoxP*, 26774; *Hsf*: 41581; *Trl* 41582. <u>FlyORF stock (with associated</u> <u>targeted genes):</u> *NF-yB*, F001895; *CG4360*, F000063; *dmrt93B*, F000445; *Ets96B*, F000142; *Ets21C*, F000624; *srp*, F000720; *GATAd*, F000714; *Hsf*, F000699. VRDC stock (with associated gene): *Adf1*, 4278.

### Smurf assay recording

Flies were transferred to food containing the blue dye FD&C #1 at 2.5\% (w/v) 24 hours prior to Smurfs counting. The dye is added as the last component in the food preparation, and dissolved in it. At the moment of the counting, flies were transferred back on normal food. All the flies are therefore spending the same amount of time on blue food, in order not to introduce bias in the counts. Note that with the following method we are not having information about the time at which the Smurfs are becoming such. However, as the Smurfs spend on average the same amount of time in this phase^20^, recording the presence of a “mixed” Smurf population provides a good estimation of their appearance in the population. Smurf counting was performed every two weeks while the population was in the survival plateau, and every week once it exited it.

### RNA extraction and qPCR quantification

Extraction of RNA was performed using the Trizol protocol as in^91^, adapted to the amount of tissue used. Each sample corresponds to a mixture of 3 flies for the RT-qPCR experiments and 8 flies for the RNA-Seq. For the RT-qPCRs, RNA was retro-transcribed using the Applied Biosystems cDNA Reverse Transcription Kit. RT-qPCR was subsequently performed using the Applied Biosystem PowerTrack SYBR Master Mix on Biorad CFX 96. Primers were designed on Benchling. *Adf1* Fw: ACAGCCCTTCAACGGCA, *Adf1* Rw: CGGCTCGTAGAAGTATGGCT; *CG4360* Fw: CAGCAGAGCACCCTTACCAA, *CG4360* Rw: GGAGCGGGCATTGAGTGAT; *Trl* Fw: TCCTATCCACGCCAAAGGCAAA, Trl Rw: TAGCAAATGGGGCAAGTAGCAGG; *Act* Fw: CCATCAGCCAGCAGTCGTCTA, *Act* Rw: ACCAGAGCAGCAACTTCTTCG.

## Acknowledgements

We thank Camille Garcia from the proteomic platform of Institut Jacques Monod (ProtéoSeine) for producing and pre-processing the proteomics data presented in the manuscript. We thank for his helpful comments on the manuscript.

## Contributions

M.R. conceived the presented idea and model. F.Z and M.R. conceived, planned and performed the analysis and experiments as well as wrote the manuscript. H.B. and M.B performed the RT-qPCR experiments and analysed the results. S.S.M. and J.L.M. participated in the longevity experiments. S.B. provided technical support for the analysis. C.C., F.A. and S.D. performed and analysed the metabolomics experiments. J.A. and M.R. performed and analysed the proteomics experiments. C.A. helped with the RNAseq analysis. All authors discussed the results and contributed to the final manuscript.

## Supplementary Figures

**S1.**
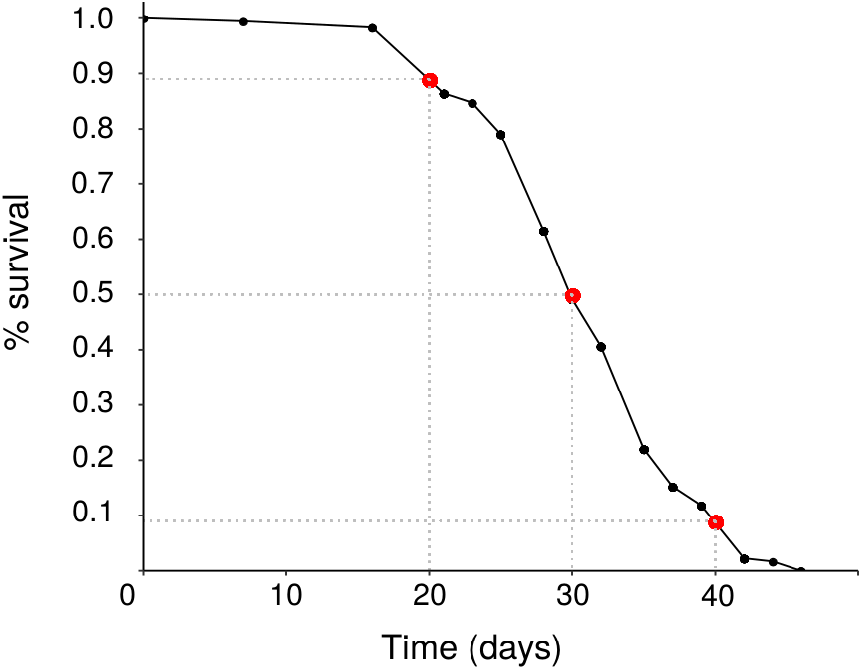
Survival curve of the *Drs*GFP population used for the RNA-seq experiment. Red dots (and axis intersecting dotted lines) highlight the sampling time points (20 days - 90% survival, 30 days - 50% survival, 40 days - 10% survival).

**S2.**
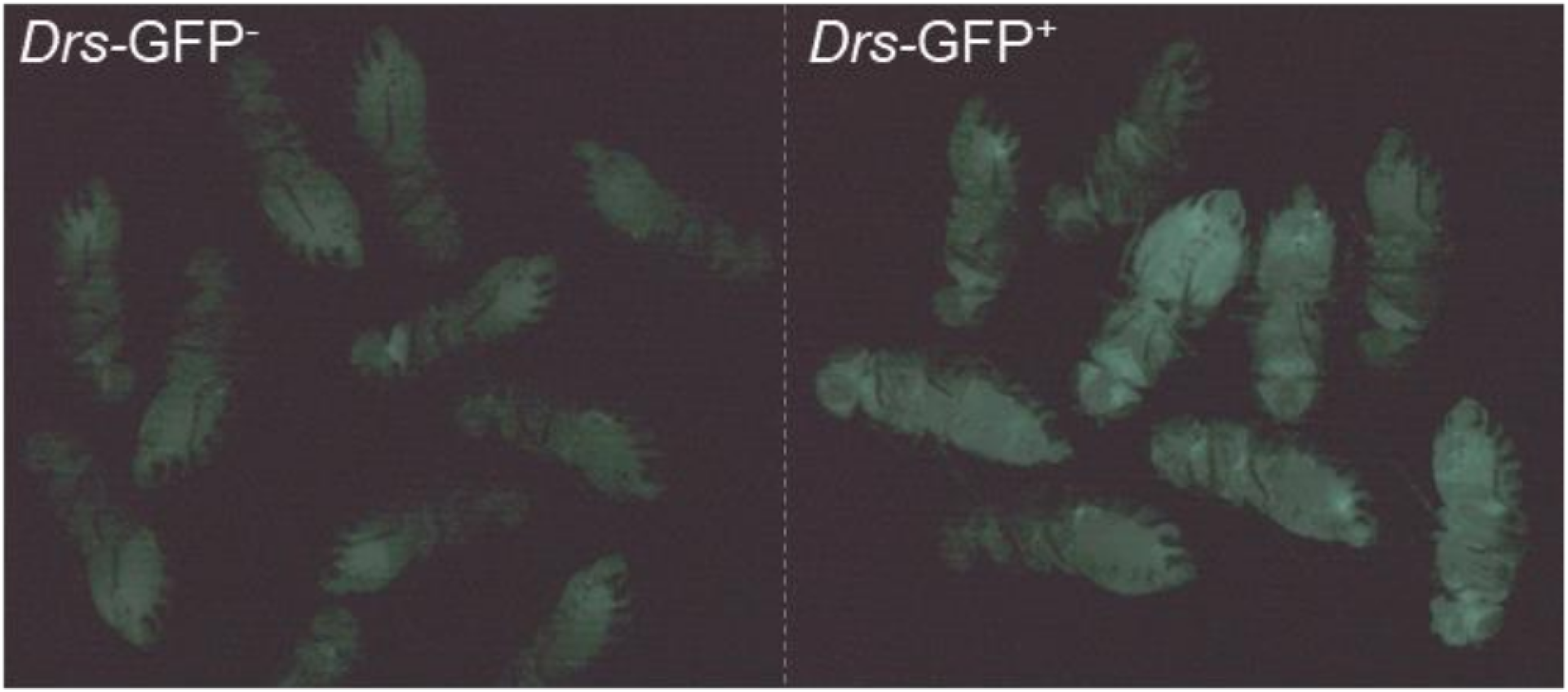
GFP^+^ and GFP^-^ flies from *Drs*-GFP line. Flies separated based on the GFP activation status (GFP^-^ on the left and GFP^+^ on the right), a possible alternative method for Smurf selection.

**S3.**
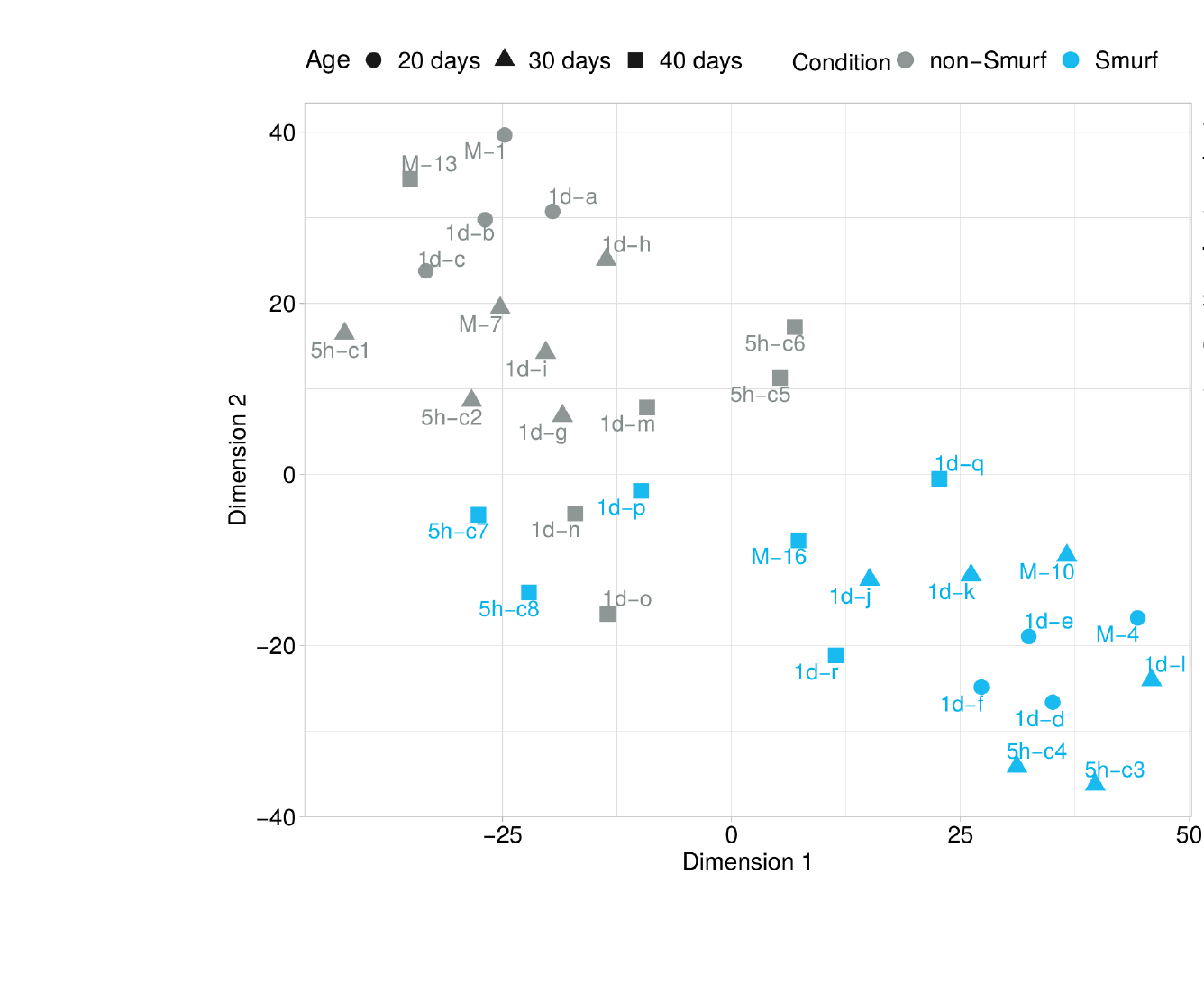
tSNE (perplexity = 10) on RNA-seq samples. tSNE is computed on all genes. Colour indicates Smurf status, symbols the age (as in legend). Similarly to the PCA results in Fig. 1a, Smurf and non-Smurfs samples form two groups. In the non-Smurf groups we can notice the samples segregating by age, while the Smurf group appear more mixed. The same “mixed” behaviour for the 40 days samples as in the PCA are identified.

**S4.**
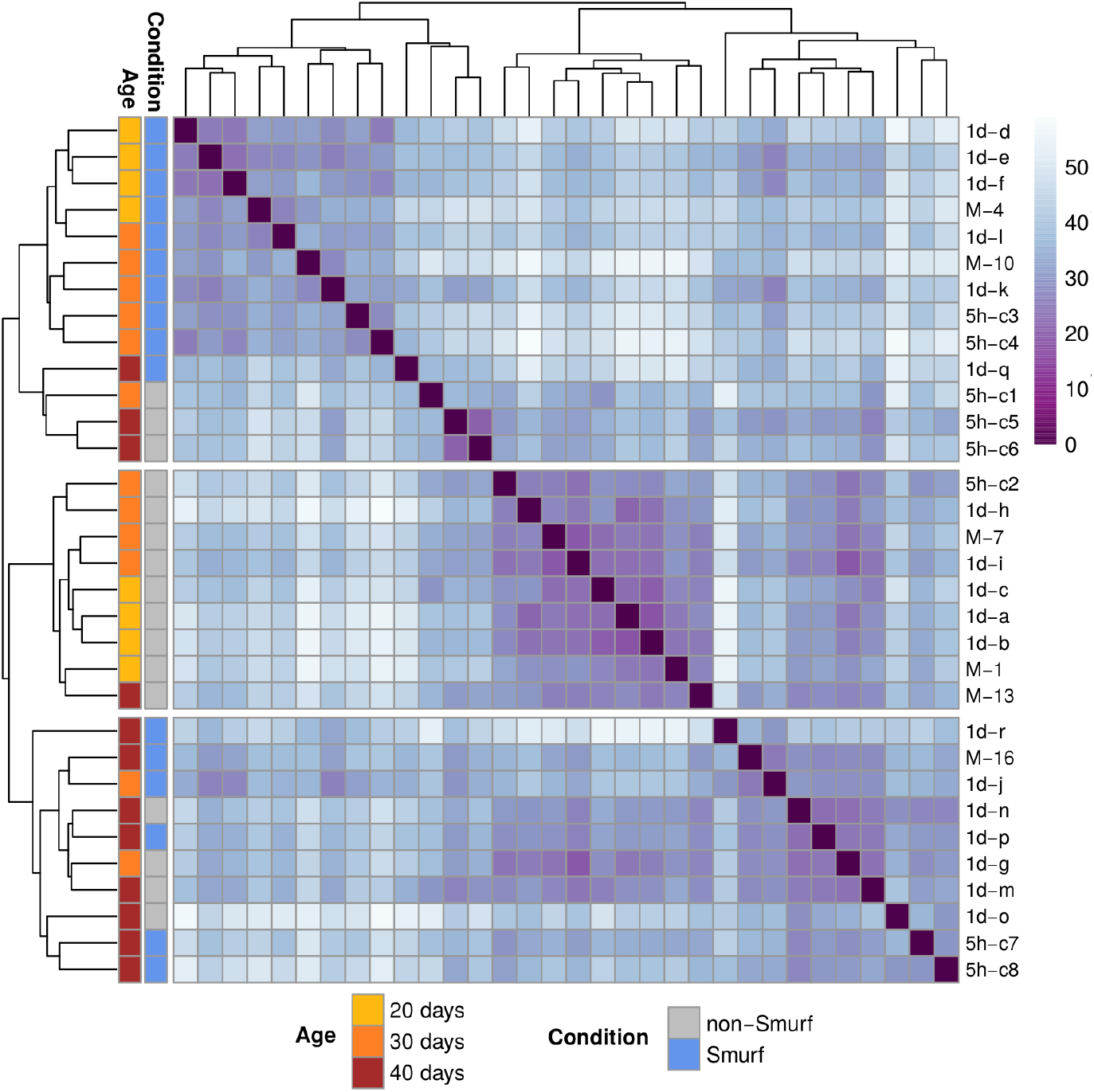
Unsupervised hierarchical clustering on sample-to-sample distance. Distance matrix (euclidean distance) is computed using all genes. Three main clusters are identified, showing good separation between Smurfs and non Smurfs but for the 40 days samples, which appear to either correlate with one of the two groups or form a third cluster independently of the Smurf status.

**S5.**
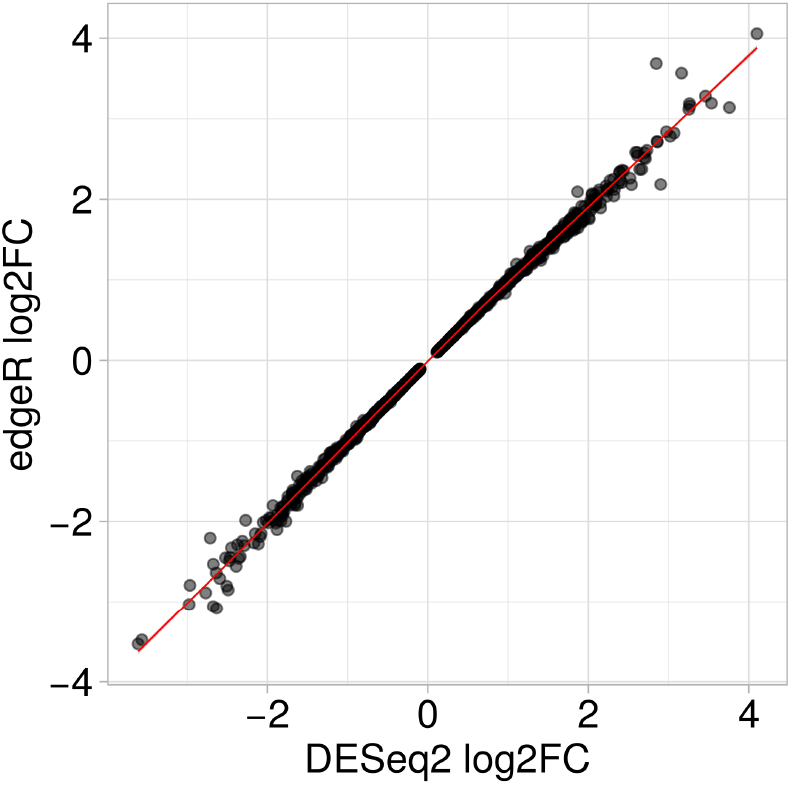
edgeR pipeline validates DESeq2 analysis. Each of the commonly 2362 DEGs identified by the two pipelines is plotted as a function of the estimated fold changes. Estimated pearson correlation between the two is 0.99.

**S6.**
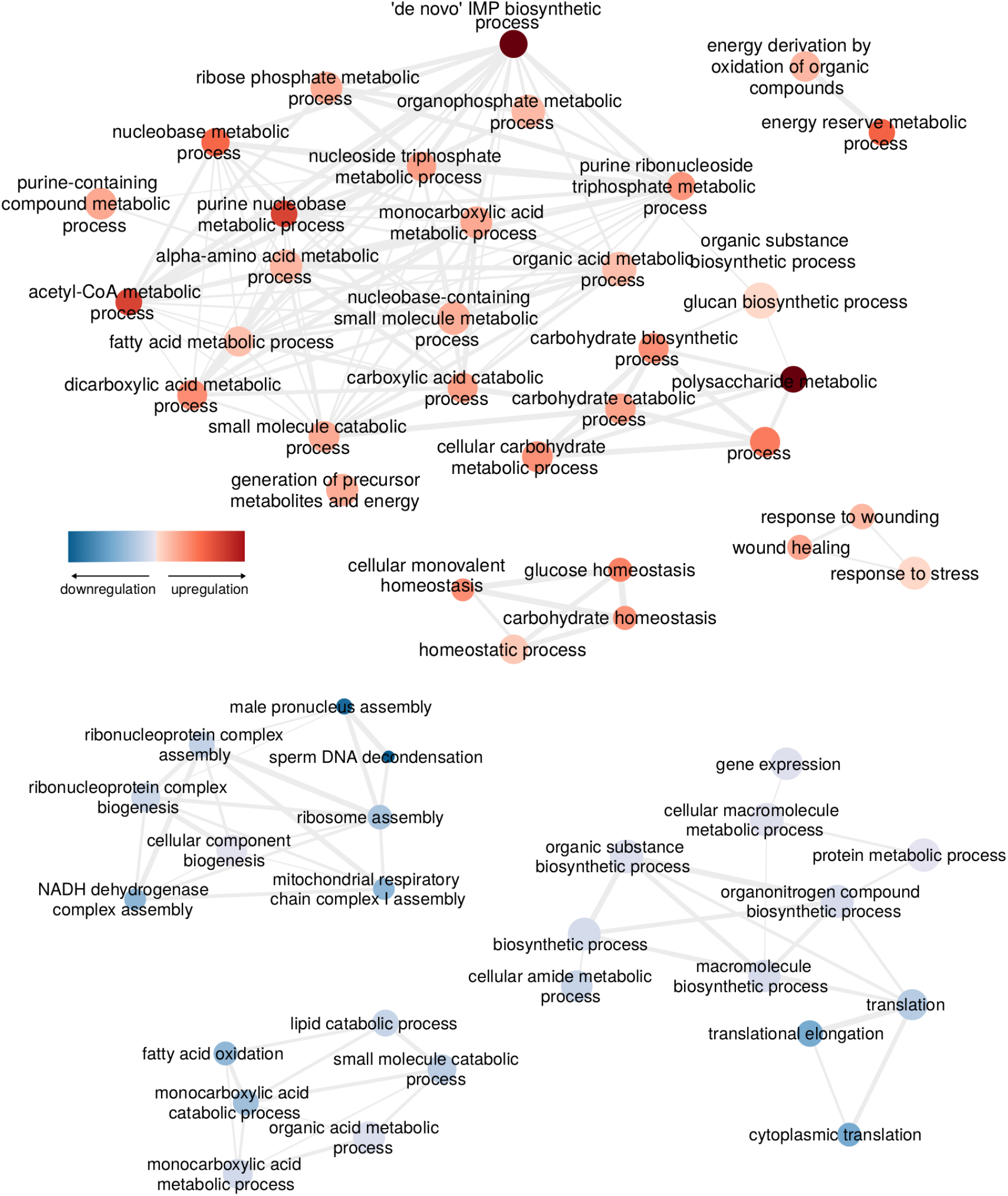
Enrichment analysis on differentially expressed proteins, Smurf vs non-Smurfs. Interconnected GO BP significant categories are here represented as a network. The color indicates the level of deregulation (Panther Fold Change estimation) - http://www.pantherdb.org/ -. The node size provides an approximate indication of the GO BP category size. Amongst the upregulated categories we mostly observe response to stress and proteins involved in metabolism (with a strong signal coming from the IMP biosynthesic process category - associated to purine metabolism, not observed in the transcriptome). The downregulated categories mostly map to ribosomal proteins, mitochondrial respiratory chain (complex I), metabolism (with the lipid catabolic process confirming what is observed in the Smurf transcriptome). The gene expression categories include numerous ribosomal proteins and should therefore not be interpreted as a signal regarding transcriptional regulation.

**S7.**
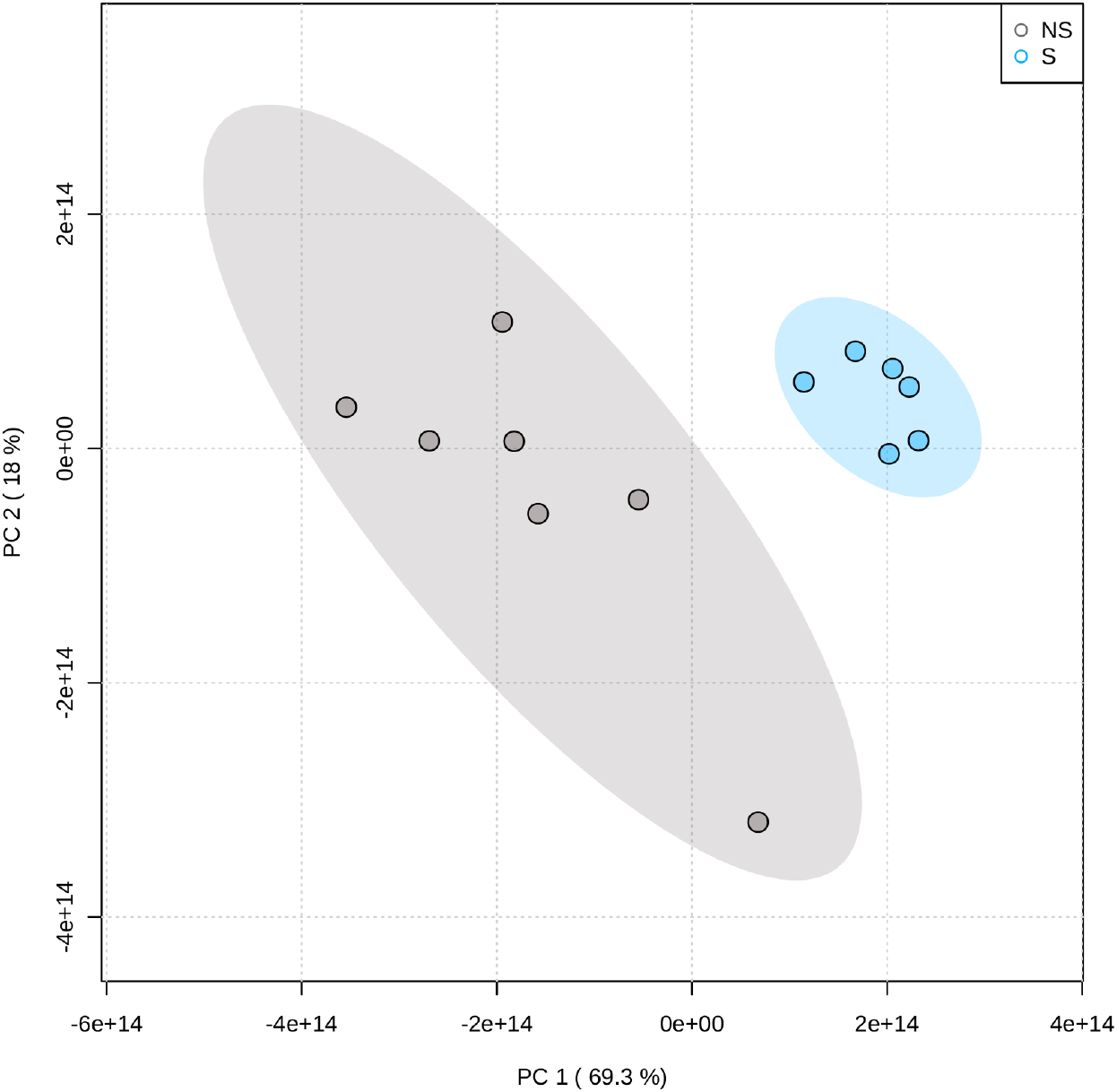
PCA performed on metabolomic data. Similarly to what occurs for th transcriptomic, the PCA on the quantification of 202 metabolites clearly separates Smurf and non-Smurf samples. PCA performed through MetaboAnalyst online platform.

**S8.**
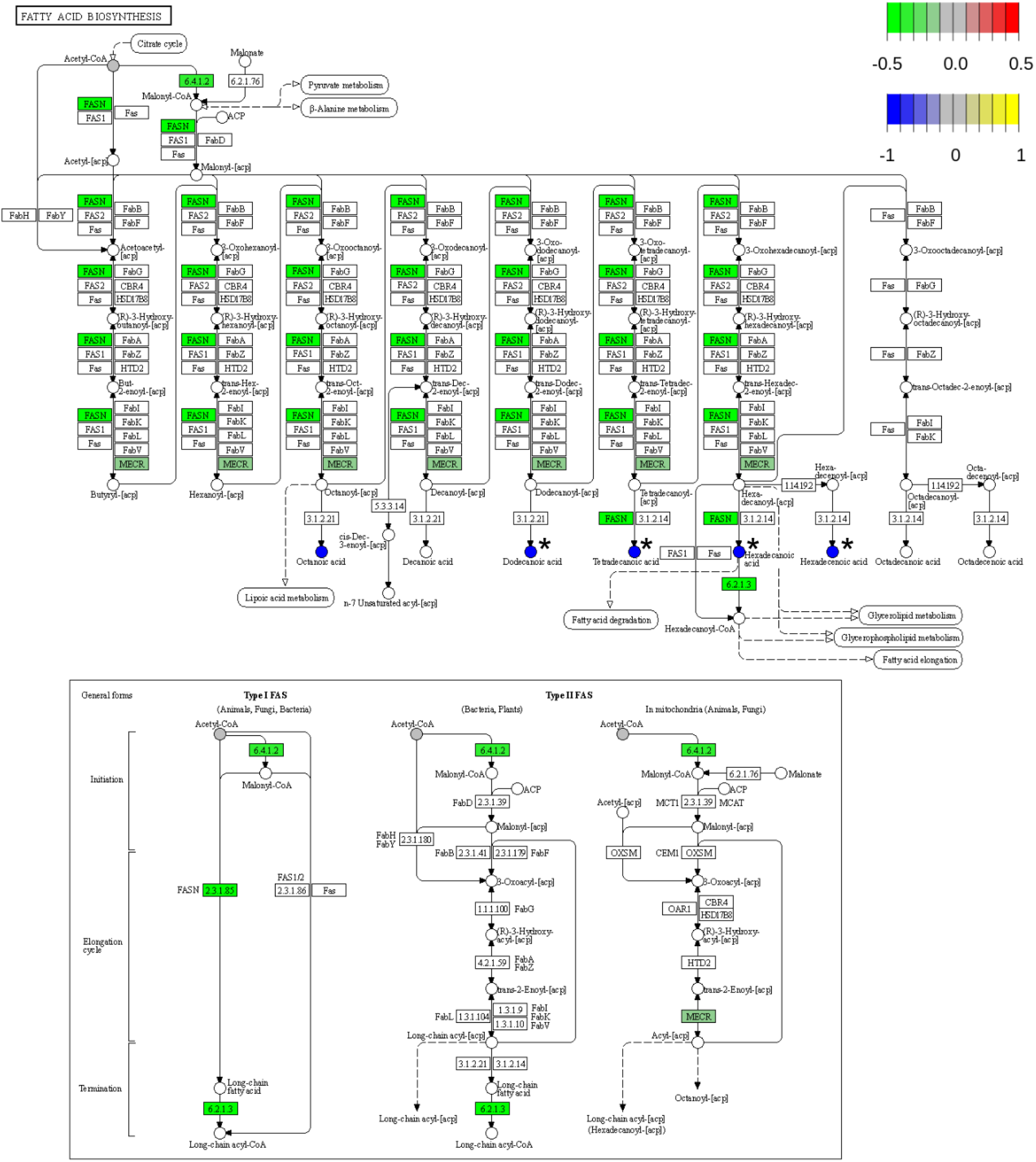
Smurf DEGs and metabolites FC on KEGG fatty acid biosynthesis pathway (dm00061). The pathview R package is used to map the genes identified as DEGs in the Smurf/non-Smurf comparison and belonging to the KEGG fatty acid pathway (dm00061). The log_2_FC estimated by the DESeq2 analysis is represented by the color scale. The detected metabolites are colored according Smurf/non-Smurf log_2_FC, and associated to a * when significant to Wilcoxon test (p-value < 0.05). The downregulation of biosynthesis-mediating enzymes is associated by a decreased presence in Smurfs of the final fatty acid products, suggesting that the transcriptional signature is functional. Enzymatic complexes are annotated through unique identification code, while genes are automatically annotated with the humane symbol. To retrieve the Drosophila gene symbols from pathway’s nodes, go to the online version and place the pointer on the gene on interest (https://www.genome.jp/pathway/dme00061).

**S9.**
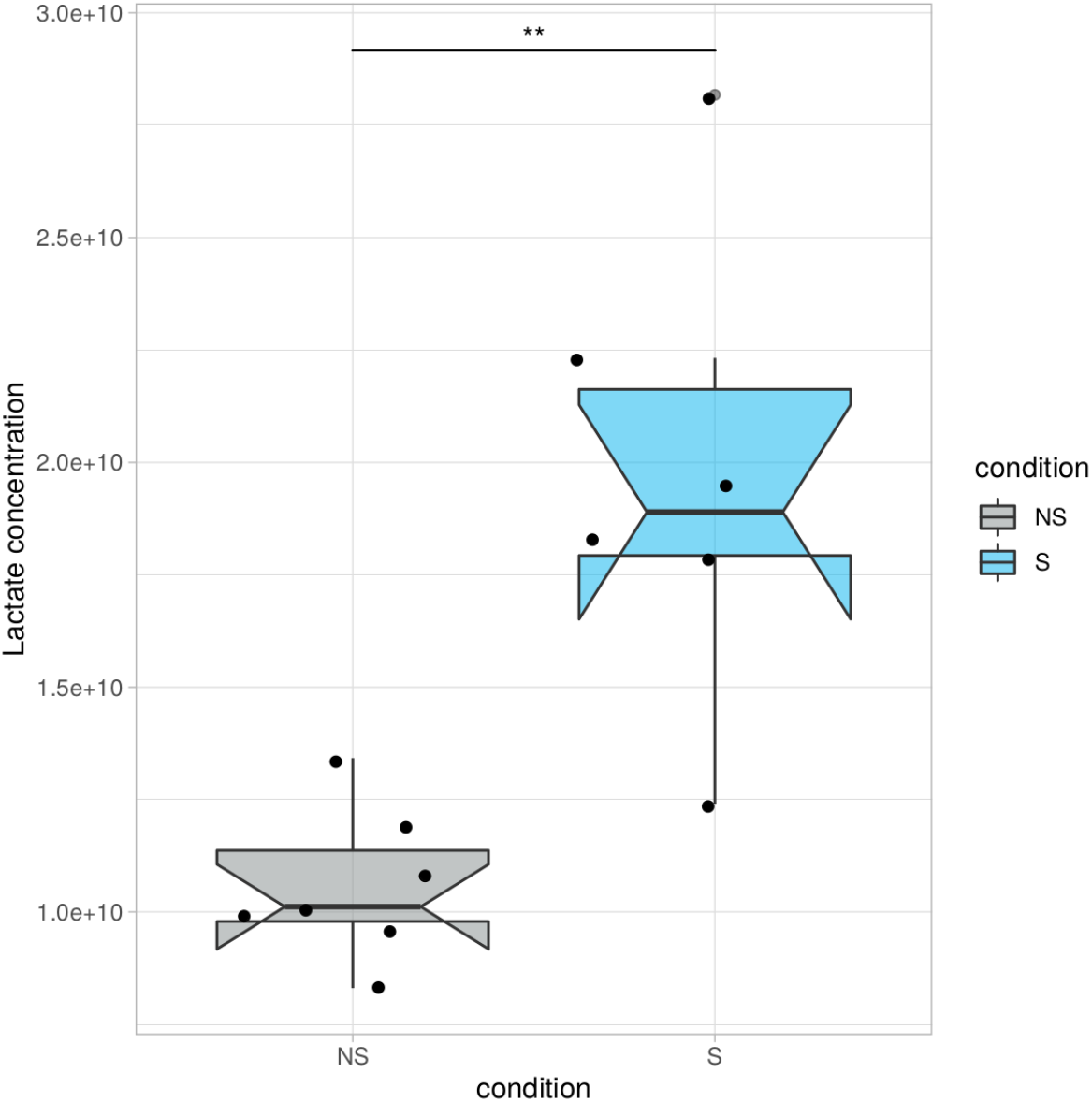
Lactic acid levels are significantly higher in Smurfs. Smurf present a significant increase in lactic acid level (log_2_FC = 0.90; **p-value < 0.001) compared to non-Smurfs. This confirms that the transcriptional upregulation of *Ldh* in Smurfs is functional. Smurfs might rely more on fermentation after glycolysis (compared to the non-Smurfs) given the general impairment experience in mitochondria.

**S10.**
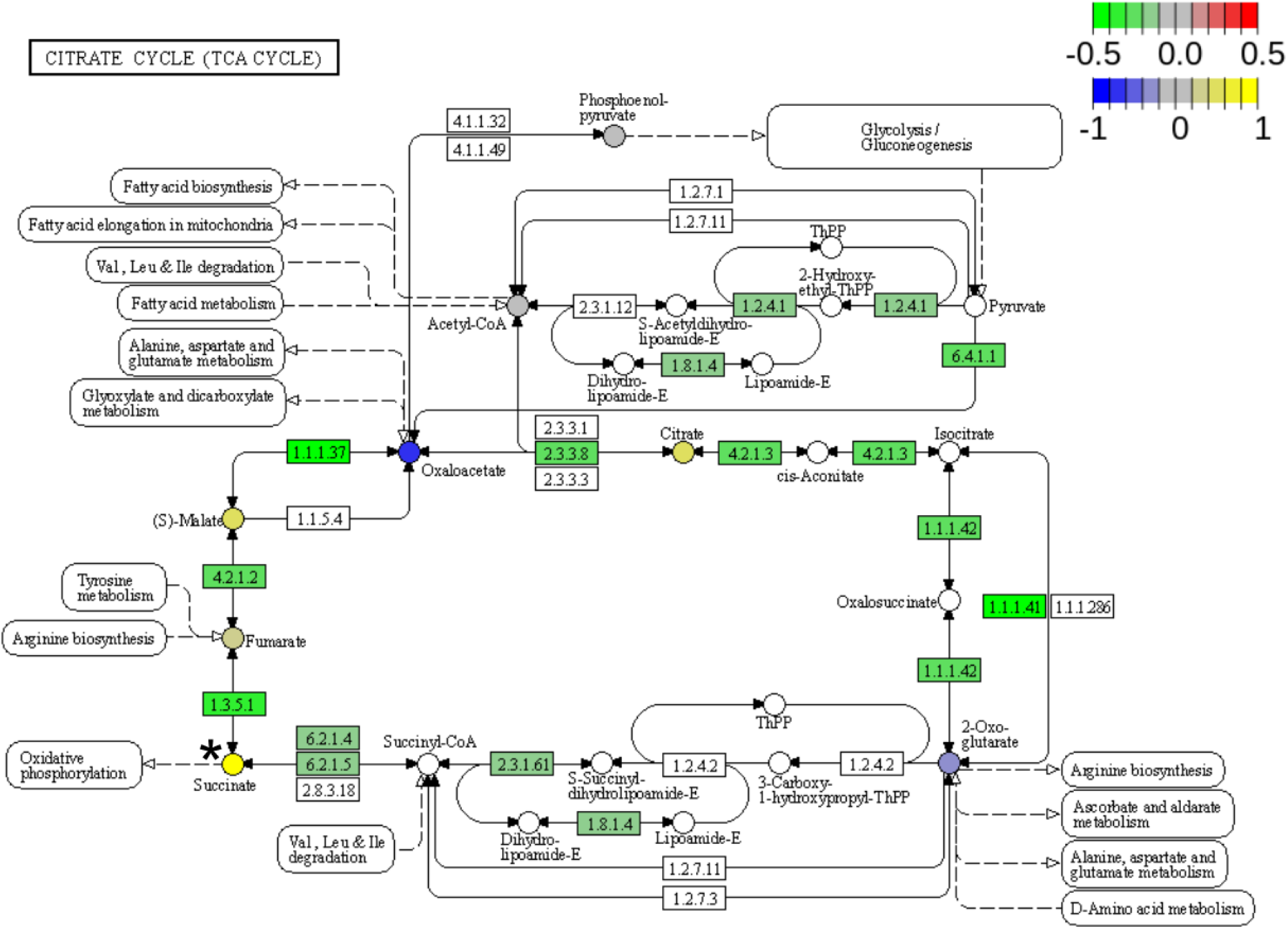
Smurf DEGs and metabolites FC on KEGG TCA cycle pathway (dm00061). The pathview R package is used to map the genes identified as DEGs in the Smurf/non-Smurf comparison and belonging to the KEGG fatty acid pathway (dm00061). The log_2_FC estimated by the DESeq2 analysis is represented by the color scale. The detected metabolites are colored according Smurf/non-Smurf log_2_FC, and associated to a * when significant to Wilcoxon test (p-value < 0.05). As already discussed in the Smurf transcriptome characterization, the TCA cycle displays wide downregulation. At a metabolomic level, the pathway missed the threshold for significance in the quantitative enrichment analysis (FDR = 0.13), and only succinate is significant to Wilcoxon test (log_2_FC = 1.28, p-value < 0.05). However, given the general impairment of mitochondrial metabolism observed at a transcriptomic and proteomic level, we believe the trend observed in the metabolomic data could still be interesting and serve as hypothesis generator for further analysis. Enzymatic complexes are annotated through unique identification code, while genes are automatically annotated with the humane symbol. To retrieve the Drosophila gene symbols from pathway’s nodes, go to the online version and place the pointer on the gene of interest (https://www.genome.jp/pathway/map00020).

**S11.**
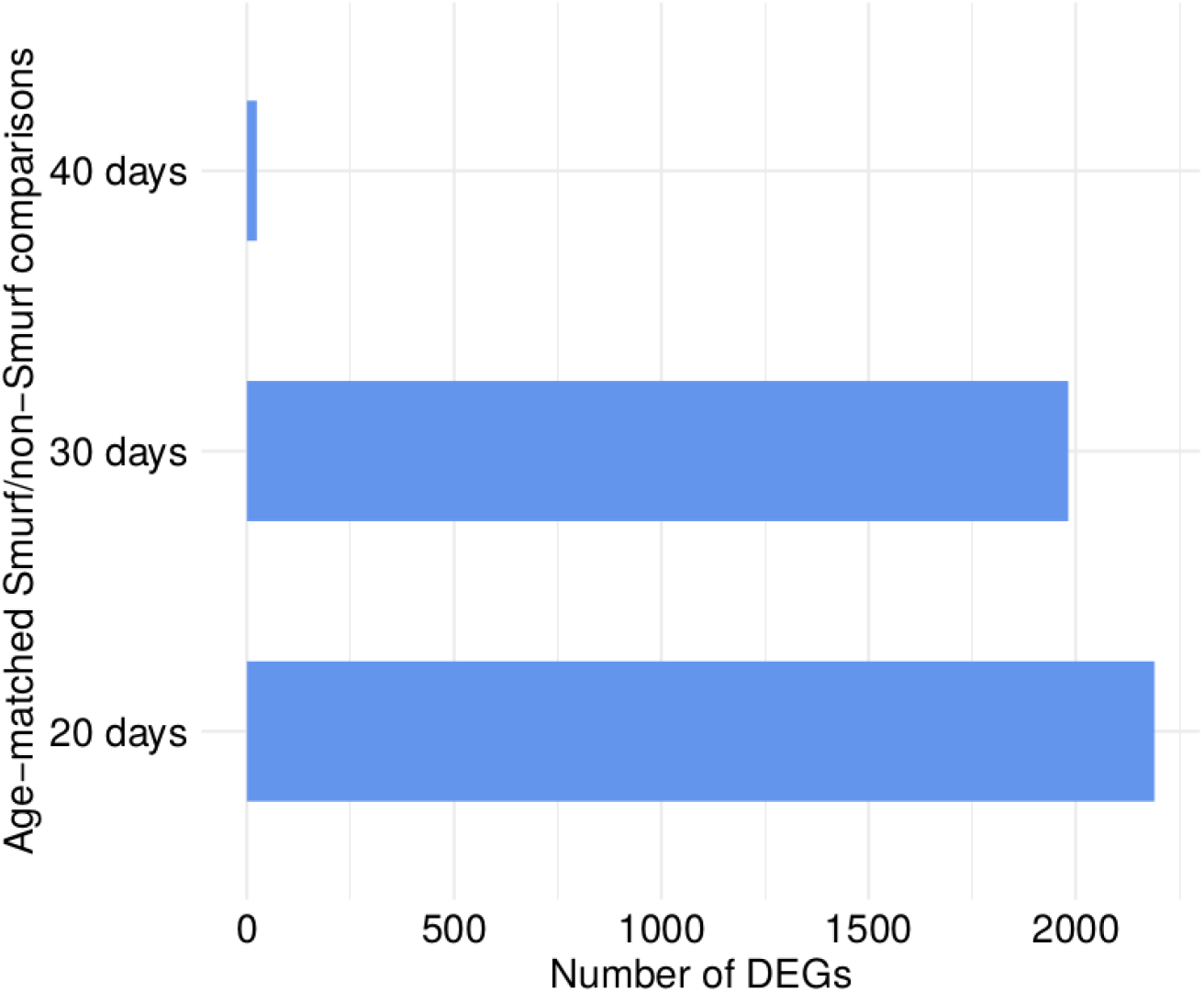
The number of DEGs in age-matched Smurf/non-Smurf comparisons decreases with chronological age. When comparing age-matched Smurfs and non-Smurfs, different number of DEGs are retrieved (DEGs_20_ = 2190, DEGs_30_ = 1982, DEGs_40_ = 24). The dramatic drop of DEGs at 40 days suggests that the transcriptome of old Smurfs and non-Smurfs are more similar than at younger ages. This was already suggested by the PCA (Fig. 1a) and might suggest that the old non-Smurfs samples, collected in the old population, are enriched in pre-Smurfs compared to their younger counterparts.

**S12.**
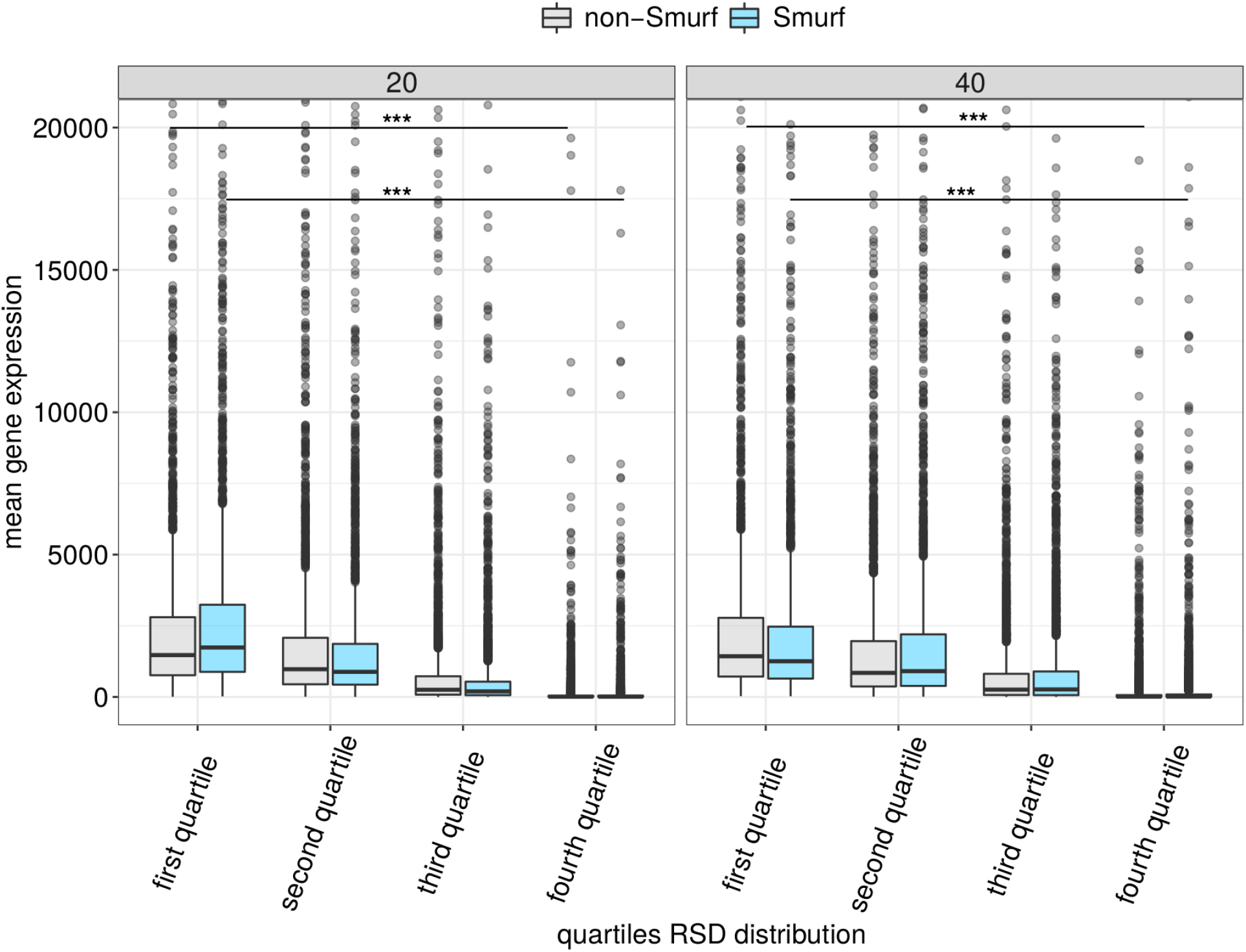
Higher relative standard deviation (RSD) in gene expression in our dataset is associated to lower counts. We divided the RSD distributions in Fig. 4b into the four quartiles (x axis) and plotted the mean gene expression of the associated genes (y axis) for Smurfs and non-Smurfs at 20 and 40 days. The mean gene expression shows a decreasing trend over the four group, proved by the significant difference between the mean gene expression of the first and fourth quartile for both Smurf and non-Smurf at 20 and 40 days (wilcoxon test, p-value < 10^e-16^).

**S13.**
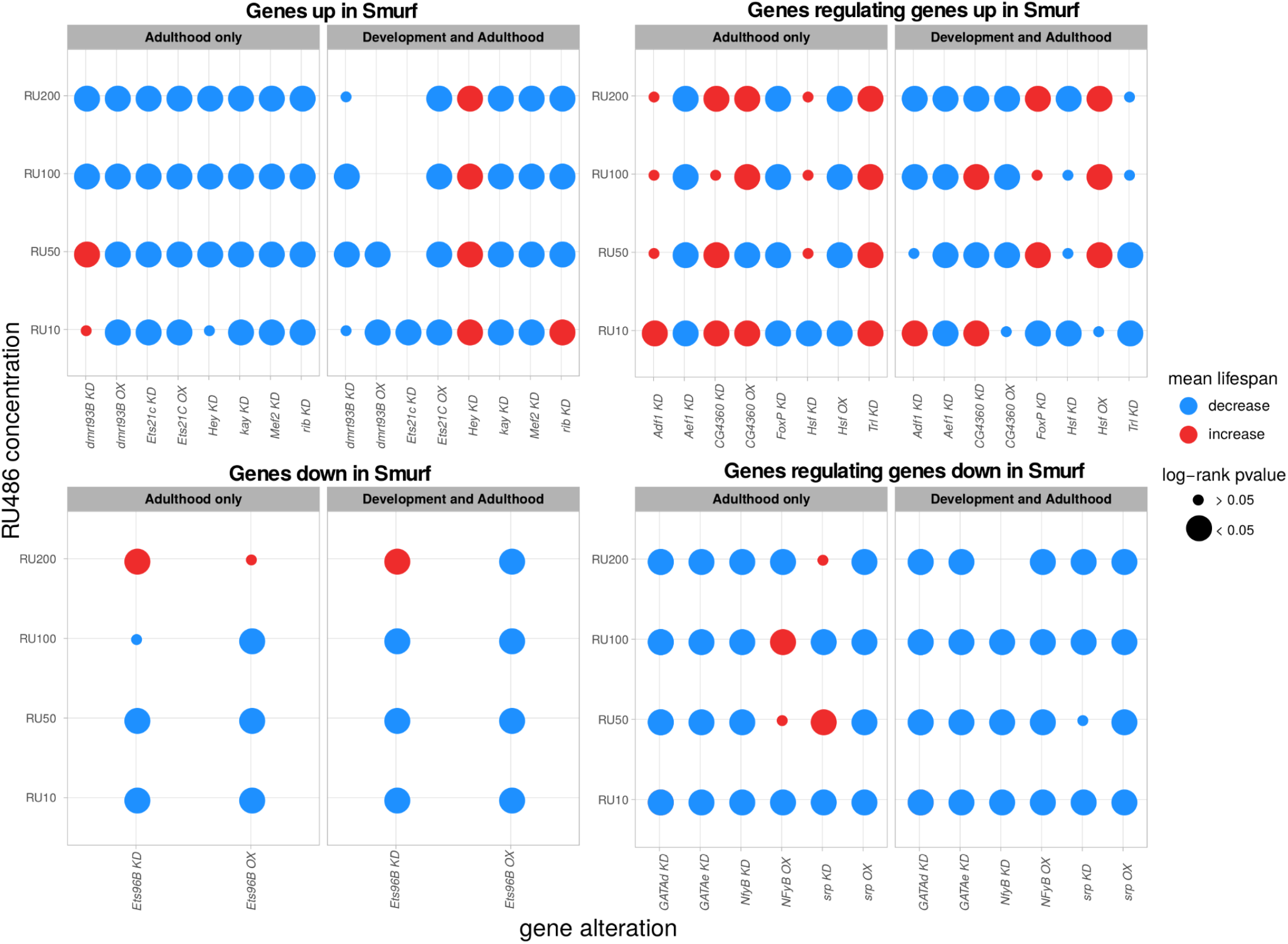
Longevity screening results. Summary of the results of the longevity screen carried out on the genes listed in Table 1. For each experiment, the 4 RU486 treatments and the two experimental setting (“adulthood only” and “development & adulthood”) are listed. The controls are not represented as they are the reference for the statistical test (log-rank) and computation of the mean lifespan change. The size of the the point indicates the significance of the difference in the longevity curve (treatment compared to control), while the colour indicates the direction of the change - decrease or increase of mean lifespan. In most cases we detected a significant difference with negative effect on the populations’ lifespan (blue large points). Interestingly most of the positive hits (red large points) map to the group of genes found by i-cisTarget as putative regulators of TFs up in Smurfs.

**S14.**
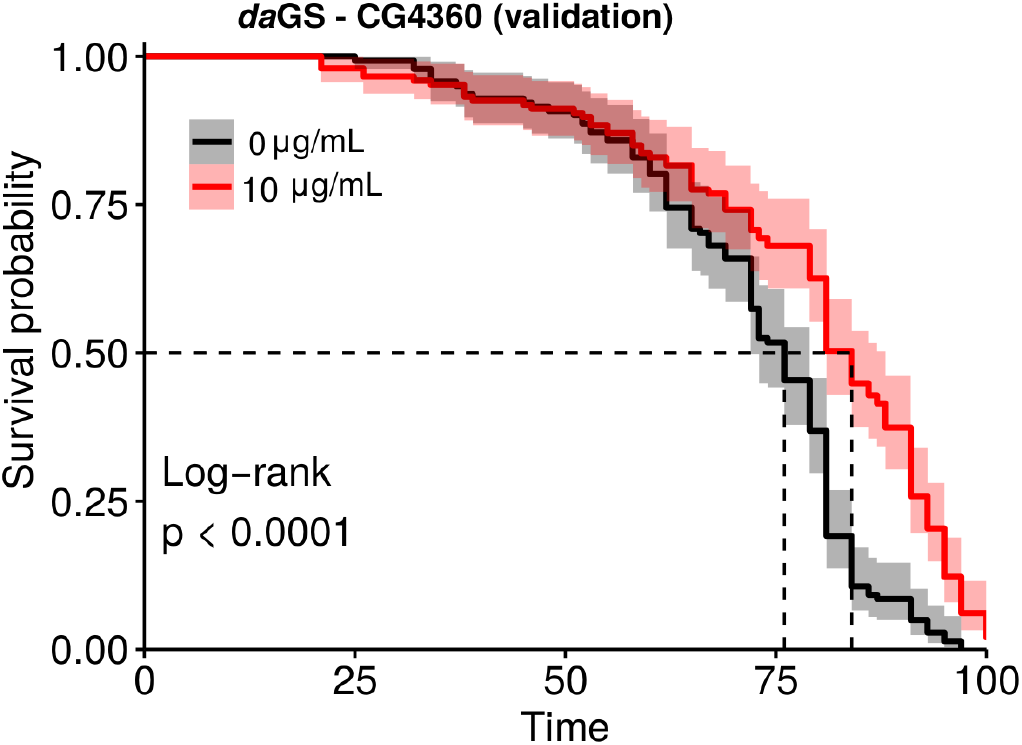
CG4360 KD (adulthood & development) validation. The effect showed in Fig.7 on the CG4360 KD (adulthood & development setting), RU10 μg/mL, is confirmed by a third independent experiment. The effect is not observed on the “adulthood only” setting. The dotted line point at the median lifespan of the populations. The effect on the mean lifespan (ML) is + 9.5% (ML_RU0_ = 71.5, ML_RU10_ = 78.5).

**S15.**
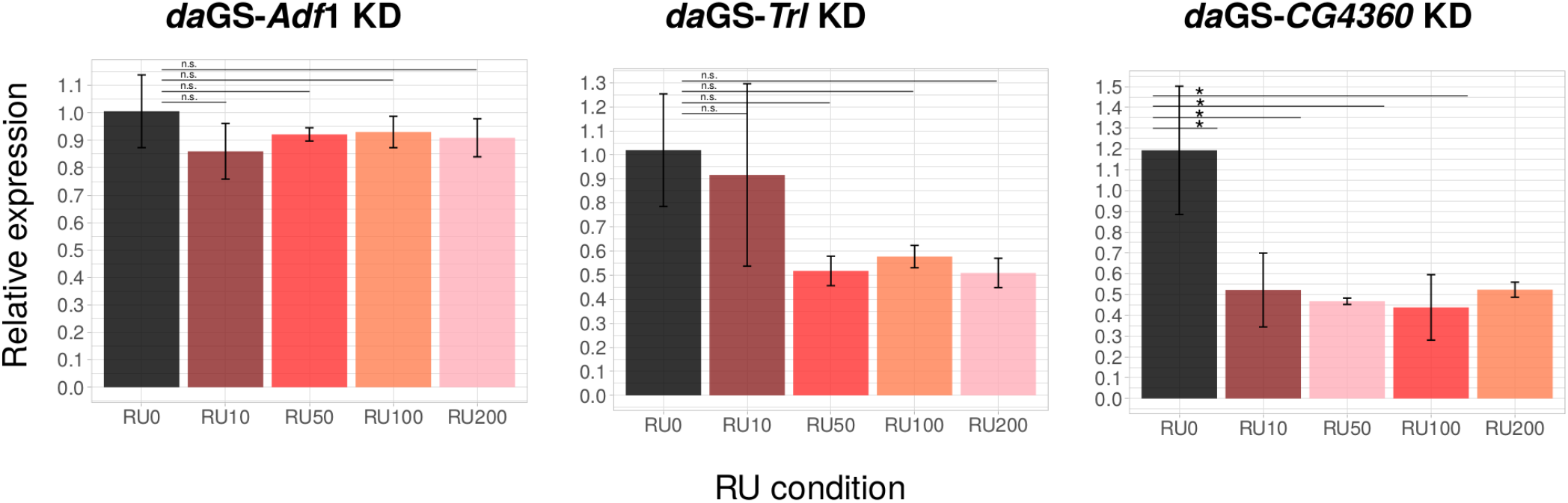
Relative gene expression quantification on KD lines (*da*GS- *Adf1*, *Trl*, *CG4360*). qPCR results for quantification of Adf1, Trl and CG4360 expression in the respective KD lines (whole body RNA extraction). In no case we observe the expected gradient, suggesting that depending on the line the RU486 induction is more or less strong independently of the amount of drug. A significant downregulation (Wilcoxon test) is detected for CG4360 at all RU486 concentrations. However, even if a trend is noticeable for *Trl* and *Adf1*, the difference in relative expression is not significant to Wilcoxon test. p-value: * = 0.05, n.s. > 0.05. *Adf1*: average 2^-ΔΔCt RU0 = 1.006, RU10 = 0.860, RU50 = 0.921, RU100 = 0.930, RU200 = 0.909; SD (standard deviation) RU0 = 0.132, RU10 = 0.101, RU50 = 0.024, RU100 = 0.057, RU200 = 0.069. *Trl*: average 2^-ΔΔCt RU0 = 1.019, RU10 = 0.916, RU50 = 0.518, RU100 = 0.577, RU200 = 0.509; SD RU0 = 0.234, RU10 = 0.380, RU50 = 0.061, RU100 = 0.046, RU200 = 0.060; *CG4360*: average 2^-ΔΔCt RU0 = 1.051, RU10 = 0.522, RU50 = 0.468, RU100 = 0.438, RU200 = 0.523; SD: RU0 = 0.379, RU10 = 0.177, RU50 = 0.015, RU100 =,0.157, RU200 = 0.036. N = 3 sample for each RU concentration, where 1 sample is the mixture of 3 flies.

**S16.**
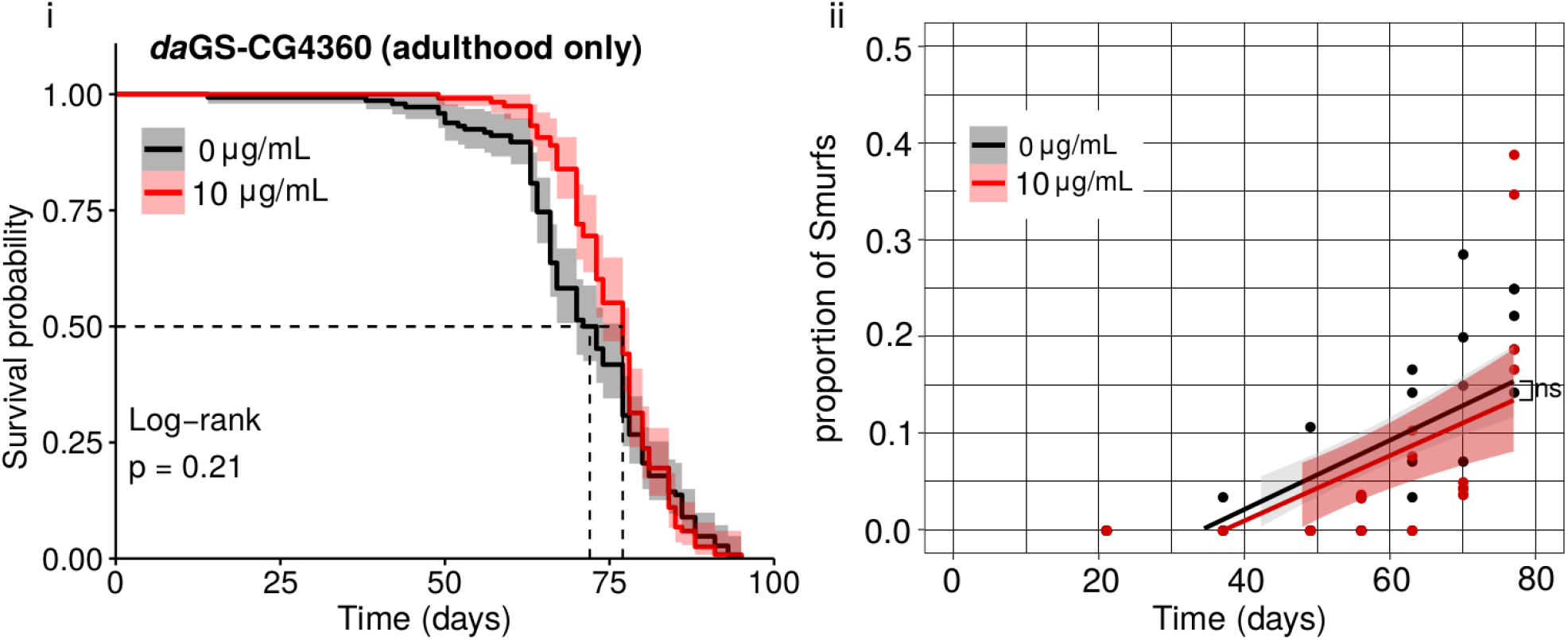
RU486 treatment does not affect lifespan. In order to confirm that the RU486 treatment alone does not affect lifespan, we performed GS longevity experiment with the daGS driver inducing *w* KD (white KD does not affect longevity). We induced the GS with RU 200 μg/mL, corresponding to the highest treatment used in our longevity experiments. No significant difference in the longevity curves is detected in the “adulthood only” setting (ML_RU0_ = 37.4, ML_RU200_ = 37.9, p-value in figure). A significant difference is detected in the “adulthood & development” setting (ML_RU0_ = 37.0, ML_RU200_ = 38.7, p-value in figure). However, the modest effect (+4.5%), together with the overlap of the confidence intervals of the curves, suggest that the effect is not biologically relevant.

**S17.**
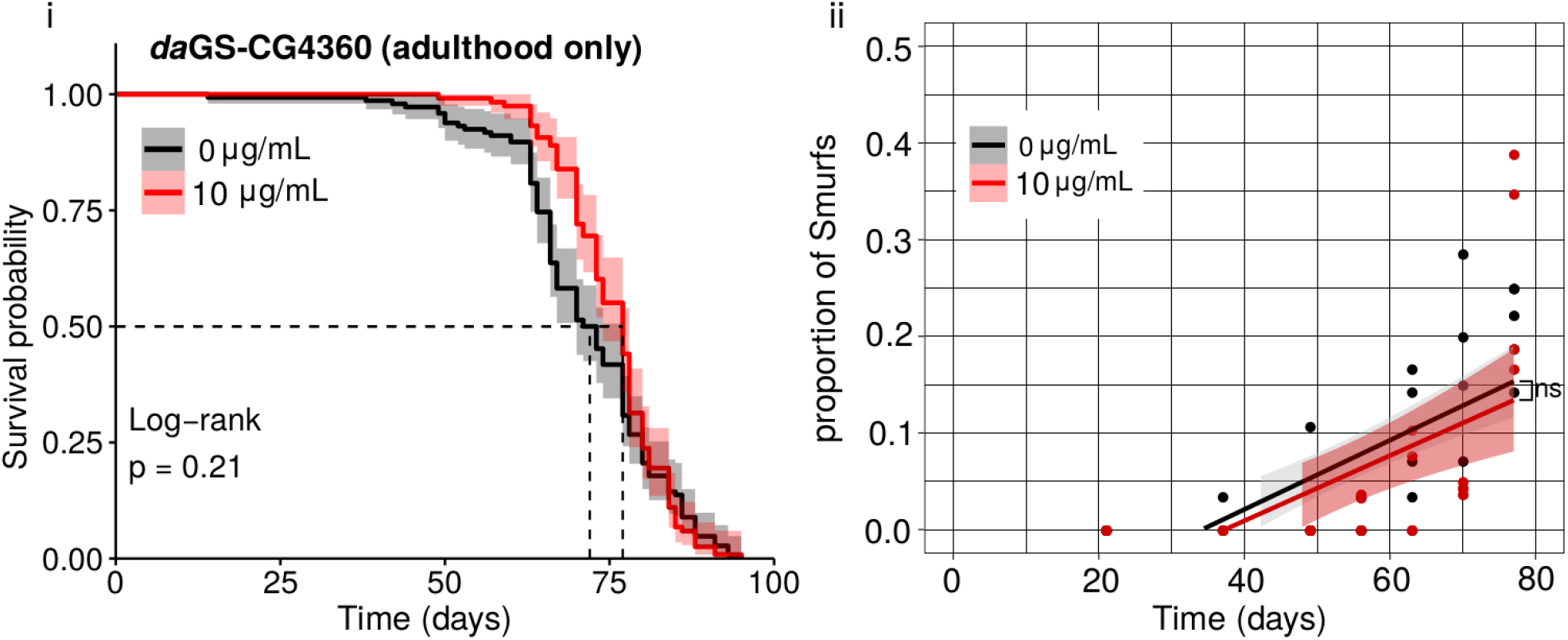
Two populations with non-significantly different lifespan experience the same Smurf proportion increase over time: the example of *CG4360* KD (adulthood only). (i) Longevity experiment. *CG4360* does not extend lifespan when knocked-down during adulthood only (ML_RU0_ = 71.6, ML_RU10_ = 75.5, log-rank p-value = 0.21). **(ii) Smurf proportion evolution over time.** The Smurf proportion significantly increases over time in the populations (slope_RU0_ = 0.0036, p-value_RU0_ = 1.50e-06, slope_RU10_ = 0.0034, p-value_RU10_ = 6.12e-04). However, no significant difference is detected between the slope of the control and the treated population (p-value = 0.84), contrary to what observed when the populations have significantly different lifespan (Fig. 6b).

**S18.**
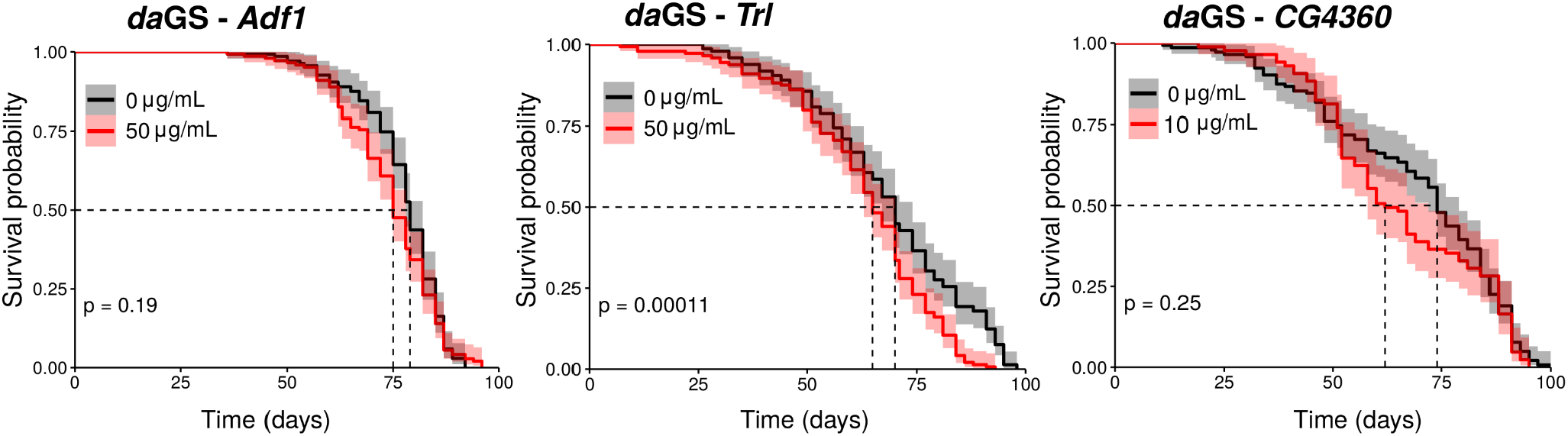
Longevity experiments on males. In order to investigat if the longevity effect found on females applies to males, we performed the experiment on males from the same GS line. Results are reported for the condition extending lifespan on females (RU50 μg/mL, adulthood only, for *Adf1* and *Trl*; RU10 μg/mL, development & adulthood, for *CG4360*). No significant effect is detected for *Adf1* and *CG4360* (log-rank p-values reported in figure; *Adf1*: ML_RU0_ = 77.1, ML_RU50_ = 74.4; *CG4360*: ML_RU0_ = 68.7, ML_RU10_ = 65.8). A significant negative effect is detected for *Trl* KD (*Trl*: ML_RU0_ = 68.5, ML_RU50_ = 62.9, -8.1%). However, the longevity curves are evolving similarly and the confidence intervals are diverging only after the T_50_; this suggests that the results need to be interpreted carefully, as the significance might not imply biological relevance.

**Fig S19.**
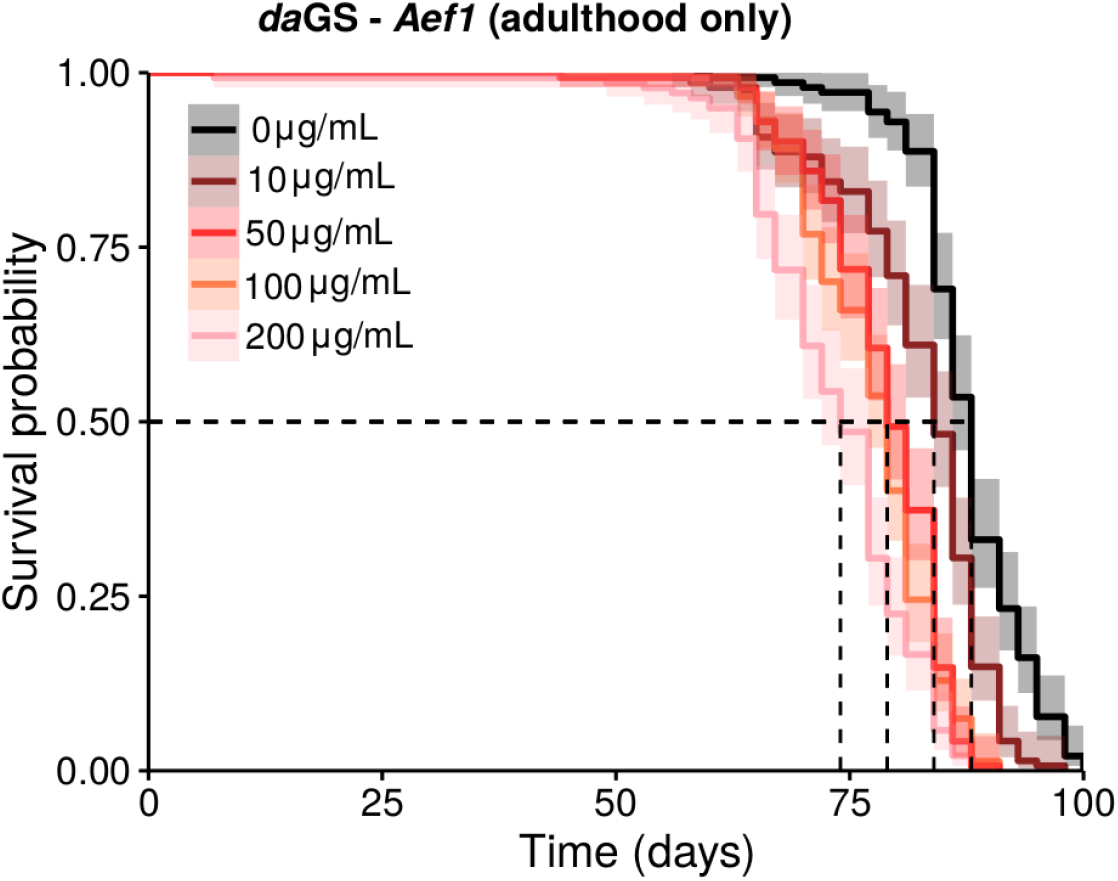
*Aef1* KD negatively affects life expectancy following the treatment gradient. *Aef1* KD negatively affects lifespan at all doses (ML_RU0_ = 87.6, ML_RU10_ = 82.2, ML_RU50_ = 78.6, ML_RU100_ = 77.4, ML_RU200_ = 73.3; p-value < 0.00001 for the log-rank test, details in Table S14). Dashed lines in figure indicate the median lifespan. The dose-dependent trend suggested by the ML values is confirmed when comparing the longevity curves of the treated populations, with only the RU50 and RU100 showing no significant difference (RU10-RU50: p-value = 3e-10; RU50-RU100: p-value = 0.2; RU100-RU200: p-value = 2e-04). Such trend suggest an effect on longevity of *Aef1* rather than a toxic effect of the KD.

## Supplementary Tables

**Table S1.** GSEA results, Smurf/non-Smurf analysis. List of the 59 significant deregulated GO BP categories (adjusted p-value < 0.05) from the GSEA analysis on the list of Smurf DEGs. Results are illustrated in Fig. 2 of the main text. GO BP category: ID and description of the biological process category; size: number of genes annotated in the category; NES: normalized enriched score; p.adjust: FDR correction on the p-value, * < 0.05, ** < 0.01.

| **GO BP category** | **size** | **NES** | **p.adjust** |
| --- | --- | --- | --- |
| GO:0050829 defense response to Gram-negative bacterium | 39 | 2.343910 | ** |
| GO:0019731 antibacterial humoral response | 29 | 2.335396 | ** |
| GO:0070482 response to oxygen levels | 35 | 2.322513 | ** |
| GO:0006457 protein folding | 44 | 2.273671 | ** |
| GO:0019730 antimicrobial humoral response | 53 | 2.272312 | ** |
| GO:0006458 'de novo' protein folding | 15 | 2.253483 | ** |
| GO:0051084 'de novo' posttranslational protein folding | 15 | 2.253483 | ** |
| GO:0051085 chaperone cofactor-dependent protein refolding | 15 | 2.253483 | ** |
| GO:0061077 chaperone-mediated protein folding | 25 | 2.249745 | ** |
| GO:0006986 response to unfolded protein | 16 | 2.241839 | ** |
| GO:0035966 response to topologically incorrect protein | 16 | 2.241839 | ** |
| GO:0034620 cellular response to unfolded protein | 15 | 2.234994 | ** |
| GO:0035967 cellular response to topologically incorrect protein | 15 | 2.234994 | ** |
| GO:0042026 protein refolding | 15 | 2.116671 | ** |
| GO:0006959 humoral immune response | 63 | 2.092197 | ** |
| GO:0001666 response to hypoxia | 23 | 2.082786 | * |
| GO:0036293 response to decreased oxygen levels | 26 | 2.057510 | ** |
| GO:0042742 defense response to bacterium | 80 | 2.042555 | ** |
| GO:0051276 chromosome organization | 61 | 2.027083 | ** |
| GO:0050830 defense response to Gram-positive bacterium | 32 | 2.011920 | * |
| GO:0031347 regulation of defense response | 33 | 1.971860 | * |
| GO:0006955 immune response | 107 | 1.951750 | ** |
| GO:0061057 peptidoglycan recognition protein signaling pathway | 10 | 1.935157 | * |
| GO:0006952 defense response | 138 | 1.905159 | ** |
| GO:0098542 defense response to other organism | 103 | 1.869388 | * |
| GO:0009617 response to bacterium | 97 | 1.844119 | * |
| GO:0009266 response to temperature stimulus | 48 | 1.789309 | * |
| GO:0002376 immune system process | 137 | 1.789089 | ** |
| GO:0009628 response to abiotic stimulus | 125 | 1.749653 | * |
| GO:0071310 cellular response to organic substance | 75 | 1.745146 | * |
| GO:0009607 response to biotic stimulus | 138 | 1.732527 | ** |
| GO:0043207 response to external biotic stimulus | 138 | 1.732527 | ** |
| GO:0051707 response to other organism | 138 | 1.732527 | ** |
| GO:0006950 response to stress | 356 | 1.731664 | ** |
| GO:0019538 protein metabolic process | 534 | -1.395434 | * |
| GO:0055086 nucleobase-containing small molecule metabolic process | 117 | -1.614257 | * |
| GO:0055114 oxidation-reduction process | 225 | -1.618842 | * |
| GO:0008610 lipid biosynthetic process | 68 | -1.728761 | * |
| GO:0015980 energy derivation by oxidation of organic compounds | 52 | -1.797256 | * |
| GO:0006091 generation of precursor metabolites and energy | 73 | -1.811853 | * |
| GO:0016054 organic acid catabolic process | 54 | -1.817399 | * |
| GO:0046395 carboxylic acid catabolic process | 54 | -1.817399 | * |
| GO:0044281 small molecule metabolic process | 314 | -1.850554 | * |
| GO:0007304 chorion-containing eggshell formation | 30 | -1.878965 | * |
| GO:0042254 ribosome biogenesis | 46 | -1.902000 | * |
| GO:0042180 cellular ketone metabolic process | 20 | -1.904478 | * |
| GO:0042775 mitochondrial ATP synthesis coupled electron transport | 25 | -1.911838 | * |
| GO:0030703 eggshell formation | 31 | -1.924518 | * |
| GO:0022904 respiratory electron transport chain | 26 | -1.933564 | * |
| GO:0042773 ATP synthesis coupled electron transport | 26 | -1.933564 | * |
| GO:0044282 small molecule catabolic process | 72 | -1.937346 | * |
| GO:0022900 electron transport chain | 27 | -1.956907 | * |
| GO:0034754 cellular hormone metabolic process | 13 | -1.986510 | * |
| GO:0045455 ecdysteroid metabolic process | 13 | -1.986510 | * |
| GO:0045333 cellular respiration | 39 | -1.998705 | * |
| GO:0006508 proteolysis | 267 | -2.029165 | * |
| GO:0042445 hormone metabolic process | 21 | -2.039538 | * |
| GO:0016125 sterol metabolic process | 11 | -2.125654 | ** |
| GO:0008202 steroid metabolic process | 19 | -2.179092 | ** |

**Table S2.** Quantitative enrichment analysis on metabolites profile (S/NS), significant hits. Quantitative enrichment analysis on metabolites quantification (from MetaboAnalyst) results in 13 significant KEGG pathways. The TCA cycle missed the 5% significant threshold (FDR = 0.13), but most of the associated metabolites are present in the pyruvate metabolism pathway. In confirmation of what is seen with the transcriptomic, we find fatty acid metabolism associated pathways. A signal from amino acids metabolism is also detected. Metabolite set: KEGG pathway; Total: number of metabolites in the pathway; FDR: adjusted p-value.

| Metabolite Set | Total | Hits | FDR |
| --- | --- | --- | --- |
| Valine, leucine and isoleucine degradation | 40 | 3 | 3.4984E-4 |
| Biosynthesis of unsaturated fatty acids | 36 | 5 | 5.067E-4 |
| alpha-Linolenic acid metabolism | 13 | 2 | 5.067E-4 |
| Linoleic acid metabolism | 5 | 1 | 5.067E-4 |
| Fatty acid biosynthesis | 47 | 4 | 0.0020546 |
| Fatty acid degradation | 39 | 3 | 0.0020546 |
| Fatty acid elongation | 38 | 2 | 0.0020546 |
| Pyruvate metabolism | 22 | 6 | 0.0071512 |
| Valine, leucine and isoleucine biosynthesis | 8 | 2 | 0.0078342 |
| Phenylalanine metabolism | 10 | 3 | 0.043592 |
| Phenylalanine, tyrosine and tryptophan biosynthesis | 4 | 2 | 0.043592 |
| Thiamine metabolism | 7 | 1 | 0.051592 |
| Cysteine and methionine metabolism | 33 | 5 | 0.059126 |

**Table S3.** GSEA analysis on old Smurfs/young Smurfs. List of the 125 deregulated GO BP categories (adjusted p-value < 0.05) from the GSEA analysis on the list of old Smurf DEGs. Results are partially illustrated in Fig. 4 of the main text. GO BP category: ID and description of the biological process category; size: number of genes annotated in the category; NES: normalized enriched score; p.adjust: FDR correction on the p-value, 0.01 < * < 0.05.

| GO BP category | size | NES | p.adjust |
| --- | --- | --- | --- |
| GO:0010038 response to metal ion | 15 | 1.983838 | * |
| GO:0010035 response to inorganic substance | 22 | 1.782223 | * |
| GO:0006030 chitin metabolic process | 25 | 1.778918 | * |
| GO:1901071 glucosamine-containing compound metabolic process | 27 | 1.775833 | * |
| GO:0006022 aminoglycan metabolic process | 29 | 1.750849 | * |
| GO:0006040 amino sugar metabolic process | 28 | 1.737450 | * |
| GO:0007606 sensory perception of chemical stimulus | 24 | 1.622532 | * |
| GO:0055114 oxidation-reduction process | 134 | 1.526567 | * |
| GO:0005975 carbohydrate metabolic process | 68 | 1.515730 | * |
| GO:0017144 drug metabolic process | 64 | 1.513947 | * |
| GO:1901988 negative regulation of cell cycle phase transition | 17 | -1.979813 | * |
| GO:1901991 negative regulation of mitotic cell cycle phase transition | 17 | -1.979813 | * |
| GO:0034470 ncRNA processing | 26 | -1.998833 | * |
| GO:0051129 negative regulation of cellular component organization | 37 | -2.000688 | * |
| GO:0045930 negative regulation of mitotic cell cycle | 19 | -2.006150 | * |
| GO:0043254 regulation of protein complex assembly | 28 | -2.009431 | * |
| GO:0022613 ribonucleoprotein complex biogenesis | 34 | -2.026966 | * |
| GO:0010769 regulation of cell morphogenesis involved in differentiation | 23 | -2.043760 | * |
| GO:0006342 chromatin silencing | 25 | -2.044283 | * |
| GO:0045814 negative regulation of gene expression, epigenetic | 25 | -2.044283 | * |
| GO:0030951 establishment or maintenance of microtubule cytoskeleton polarity | 13 | -2.050560 | * |
| GO:0050770 regulation of axonogenesis | 14 | -2.057130 | * |
| GO:0060810 intracellular mRNA localization involved in pattern specification process | 14 | -2.082873 | * |
| GO:0060811 intracellular mRNA localization involved in anterior/posterior axis specification | 14 | -2.082873 | * |
| GO:0043085 positive regulation of catalytic activity | 24 | -2.084854 | * |
| GO:0007338 single fertilization | 11 | -2.090751 | * |
| GO:0009566 fertilization | 11 | -2.090751 | * |
| GO:1990778 protein localization to cell periphery | 14 | -2.098238 | * |
| GO:0031400 negative regulation of protein modification process | 15 | -2.104281 | * |
| GO:0030952 establishment or maintenance of cytoskeleton polarity | 15 | -2.111130 | * |
| GO:0007265 Ras protein signal transduction | 34 | -2.117713 | * |
| GO:0051052 regulation of DNA metabolic process | 18 | -2.127928 | * |
| GO:0046578 regulation of Ras protein signal transduction | 17 | -2.145951 | * |
| GO:0044786 cell cycle DNA replication | 12 | -2.148274 | * |
| GO:0006399 tRNA metabolic process | 19 | -2.161945 | * |
| GO:0008298 intracellular mRNA localization | 15 | -2.164617 | * |
| GO:0033047 regulation of mitotic sister chromatid segregation | 11 | -2.168912 | * |
| GO:0006402 mRNA catabolic process | 12 | -2.170716 | * |
| GO:0051056 regulation of small GTPase mediated signal transduction | 18 | -2.179228 | * |
| GO:0007093 mitotic cell cycle checkpoint | 11 | -2.194288 | * |
| GO:0010638 positive regulation of organelle organization | 35 | -2.202552 | * |
| GO:0043087 regulation of GTPase activity | 12 | -2.225064 | * |
| GO:0000075 cell cycle checkpoint | 16 | -2.228099 | * |
| GO:0022618 ribonucleoprotein complex assembly | 20 | -2.239742 | * |
| GO:0031124 mRNA 3'-end processing | 11 | -2.253050 | * |
| GO:0000281 mitotic cytokinesis | 22 | -2.253149 | * |
| GO:1902275 regulation of chromatin organization | 33 | -2.259971 | * |
| GO:0061640 cytoskeleton-dependent cytokinesis | 35 | -2.267458 | * |
| GO:0000910 cytokinesis | 36 | -2.271666 | * |
| GO:2001252 positive regulation of chromosome organization | 20 | -2.272650 | * |
| GO:0006511 ubiquitin-dependent protein catabolic process | 38 | -2.274556 | * |
| GO:0006302 double-strand break repair | 10 | -2.286608 | * |
| GO:0010948 negative regulation of cell cycle process | 22 | -2.295652 | * |
| GO:0034968 histone lysine methylation | 12 | -2.296995 | * |
| GO:0010639 negative regulation of organelle organization | 24 | -2.302245 | * |

**Table S3. GSEA analysis on old Smurfs/young Smurfs.**
|  |  |  |  |
| --- | --- | --- | --- |
| GO:0032984 protein-containing complex disassembly | 15 | -2.313278 | * |
| GO:0000086 G2/M transition of mitotic cell cycle | 11 | -2.327275 | * |
| GO:0044839 cell cycle G2/M phase transition | 11 | -2.327275 | * |
| GO:0051303 establishment of chromosome localization | 11 | -2.327275 | * |
| GO:0065004 protein-DNA complex assembly | 22 | -2.333056 | * |
| GO:0007062 sister chromatid cohesion | 11 | -2.343420 | * |
| GO:0000380 alternative mRNA splicing, via spliceosome | 25 | -2.345997 | * |
| GO:0000381 regulation of alternative mRNA splicing, via spliceosome | 25 | -2.345997 | * |
| GO:0034728 nucleosome organization | 25 | -2.347394 | * |
| GO:0043161 proteasome-mediated ubiquitin-dependent protein catabolic process | 29 | -2.349185 | * |
| GO:0031109 microtubule polymerization or depolymerization | 11 | -2.355110 | * |
| GO:0071826 ribonucleoprotein complex subunit organization | 23 | -2.366604 | * |
| GO:0016571 histone methylation | 13 | -2.374419 | * |
| GO:0031056 regulation of histone modification | 13 | -2.374419 | * |
| GO:0007307 eggshell chorion gene amplification | 10 | -2.376503 | * |
| GO:0006333 chromatin assembly or disassembly | 22 | -2.378977 | * |
| GO:0045786 negative regulation of cell cycle | 29 | -2.382539 | * |
| GO:0072657 protein localization to membrane | 19 | -2.385931 | * |
| GO:0007052 mitotic spindle organization | 31 | -2.394738 | * |
| GO:0034660 ncRNA metabolic process | 37 | -2.397254 | * |
| GO:0010498 proteasomal protein catabolic process | 32 | -2.399087 | * |
| GO:0033045 regulation of sister chromatid segregation | 15 | -2.401296 | * |
| GO:1901987 regulation of cell cycle phase transition | 30 | -2.405488 | * |
| GO:1901990 regulation of mitotic cell cycle phase transition | 30 | -2.405488 | * |
| GO:0050000 chromosome localization | 12 | -2.405691 | * |
| GO:0006470 protein dephosphorylation | 18 | -2.445764 | * |
| GO:0007098 centrosome cycle | 15 | -2.448630 | * |
| GO:0018022 peptidyl-lysine methylation | 14 | -2.461362 | * |
| GO:0044772 mitotic cell cycle phase transition | 35 | -2.468455 | * |
| GO:0044770 cell cycle phase transition | 36 | -2.487258 | * |
| GO:0051983 regulation of chromosome segregation | 16 | -2.490788 | * |
| GO:0002181 cytoplasmic translation | 23 | -2.494502 | * |
| GO:0030261 chromosome condensation | 17 | -2.506977 | * |
| GO:0043484 regulation of RNA splicing | 28 | -2.514039 | * |
| GO:0048024 regulation of mRNA splicing, via spliceosome | 28 | -2.514039 | * |
| GO:1902850 microtubule cytoskeleton organization involved in mitosis | 35 | -2.514714 | * |
| GO:0006401 RNA catabolic process | 17 | -2.518757 | * |
| GO:0044093 positive regulation of molecular function | 32 | -2.524733 | * |
| GO:0006475 internal protein amino acid acetylation | 25 | -2.530423 | * |
| GO:0016573 histone acetylation | 25 | -2.530423 | * |
| GO:0018393 internal peptidyl-lysine acetylation | 25 | -2.530423 | * |
| GO:0018394 peptidyl-lysine acetylation | 25 | -2.530423 | * |
| GO:0071897 DNA biosynthetic process | 19 | -2.564043 | * |
| GO:0043044 ATP-dependent chromatin remodeling | 19 | -2.572069 | * |
| GO:2001251 negative regulation of chromosome organization | 15 | -2.578708 | * |
| GO:0006479 protein methylation | 16 | -2.592930 | * |
| GO:0008213 protein alkylation | 16 | -2.592930 | * |
| GO:0061982 meiosis I cell cycle process | 25 | -2.638776 | * |
| GO:0006473 protein acetylation | 27 | -2.647222 | * |
| GO:0006277 DNA amplification | 12 | -2.652135 | * |
| GO:0050684 regulation of mRNA processing | 31 | -2.666468 | * |
| GO:0006270 DNA replication initiation | 12 | -2.680627 | * |
| GO:0007127 meiosis I | 17 | -2.706746 | * |
| GO:0007088 regulation of mitotic nuclear division | 26 | -2.717261 | * |
| GO:0031123 RNA 3'-end processing | 18 | -2.732982 | * |
| GO:1903311 regulation of mRNA metabolic process | 33 | -2.740358 | * |

**Table S3. GSEA analysis on old Smurfs/young Smurfs.**
|  |  |  |  |
| --- | --- | --- | --- |
| GO:0006281 DNA repair | 31 | -2.753201 | * |
| GO:0051783 regulation of nuclear division | 27 | -2.798919 | * |
| GO:0006323 DNA packaging | 34 | -2.824701 | * |
| GO:0043543 protein acylation | 31 | -2.886683 | * |
| GO:0007143 female meiotic nuclear division | 19 | -2.953082 | * |
| GO:0006352 DNA-templated transcription, initiation | 27 | -2.965346 | * |
| GO:0006367 transcription initiation from RNA polymerase II promoter | 27 | -2.965346 | * |
| GO:0006310 DNA recombination | 17 | -2.968695 | * |
| GO:0032259 methylation | 22 | -3.031190 | * |
| GO:0043414 macromolecule methylation | 22 | -3.031190 | * |
| GO:0071103 DNA conformation change | 38 | -3.039650 | * |
| GO:0045132 meiotic chromosome segregation | 20 | -3.053654 | * |
| GO:0006261 DNA-dependent DNA replication | 34 | -3.079054 | * |
| GO:0000070 mitotic sister chromatid segregation | 34 | -3.244982 | * |

**Table S4.** GSEA analysis on old/young non- Smurfs. List of the 22 significant deregulated GO BP categories (adjusted p-value < 0.05) from the GSEA analysis on the list of old non-Smurf DEGs. Results are illustrated in Fig. 3a of the main text. GO BP category: ID and description of the biological process category; size: number of genes annotated in the category; NES: normalized enriched score; p.adjust: FDR correction on the p-value, * < 0.05, ** < 0.01.

| GO BP category | size | NES | p.adjust |
| --- | --- | --- | --- |
| GO:0009617 response to bacterium | 44 | 2.077570 | ** |
| GO:0009607 response to biotic stimulus | 52 | 2.010078 | ** |
| GO:0043207 response to external biotic stimulus | 52 | 2.010078 | ** |
| GO:0051707 response to other organism | 52 | 2.010078 | ** |
| GO:0019731 antibacterial humoral response | 16 | 1.954582 | ** |
| GO:0042742 defense response to bacterium | 35 | 1.953953 | ** |
| GO:0009605 response to external stimulus | 67 | 1.942054 | ** |
| GO:0050830 defense response to Gram-positive bacterium | 20 | 1.926474 | ** |
| GO:0050829 defense response to Gram-negative bacterium | 15 | 1.925644 | ** |
| GO:0098542 defense response to other organism | 41 | 1.893139 | ** |
| GO:0006952 defense response | 53 | 1.889830 | ** |
| GO:0002376 immune system process | 43 | 1.822360 | ** |
| GO:0051704 multi-organism process | 75 | 1.800343 | ** |
| GO:0006950 response to stress | 90 | 1.783484 | ** |
| GO:0019730 antimicrobial humoral response | 23 | 1.760254 | * |
| GO:0006955 immune response | 35 | 1.755081 | * |
| GO:0006959 humoral immune response | 26 | 1.739773 | * |
| GO:0007292 female gamete generation | 11 | -2.120223 | ** |
| GO:0048477 oogenesis | 11 | -2.120223 | ** |
| GO:0022412 cellular process involved in reproduction in multicellular organism | 15 | -2.142952 | * |
| GO:0007276 gamete generation | 16 | -2.202907 | * |
| GO:0007281 germ cell development | 13 | -2.273476 | * |

**Table S5.**
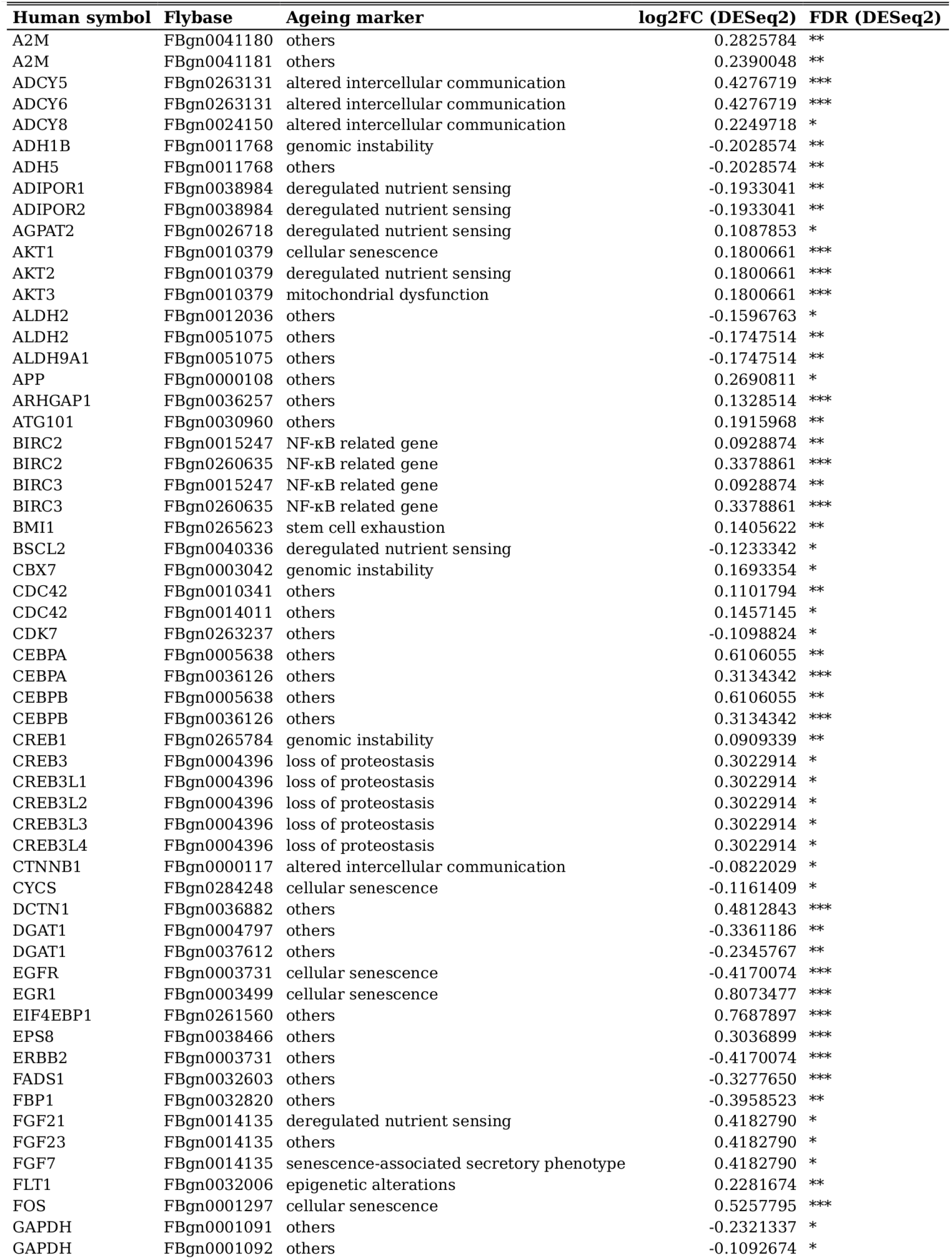
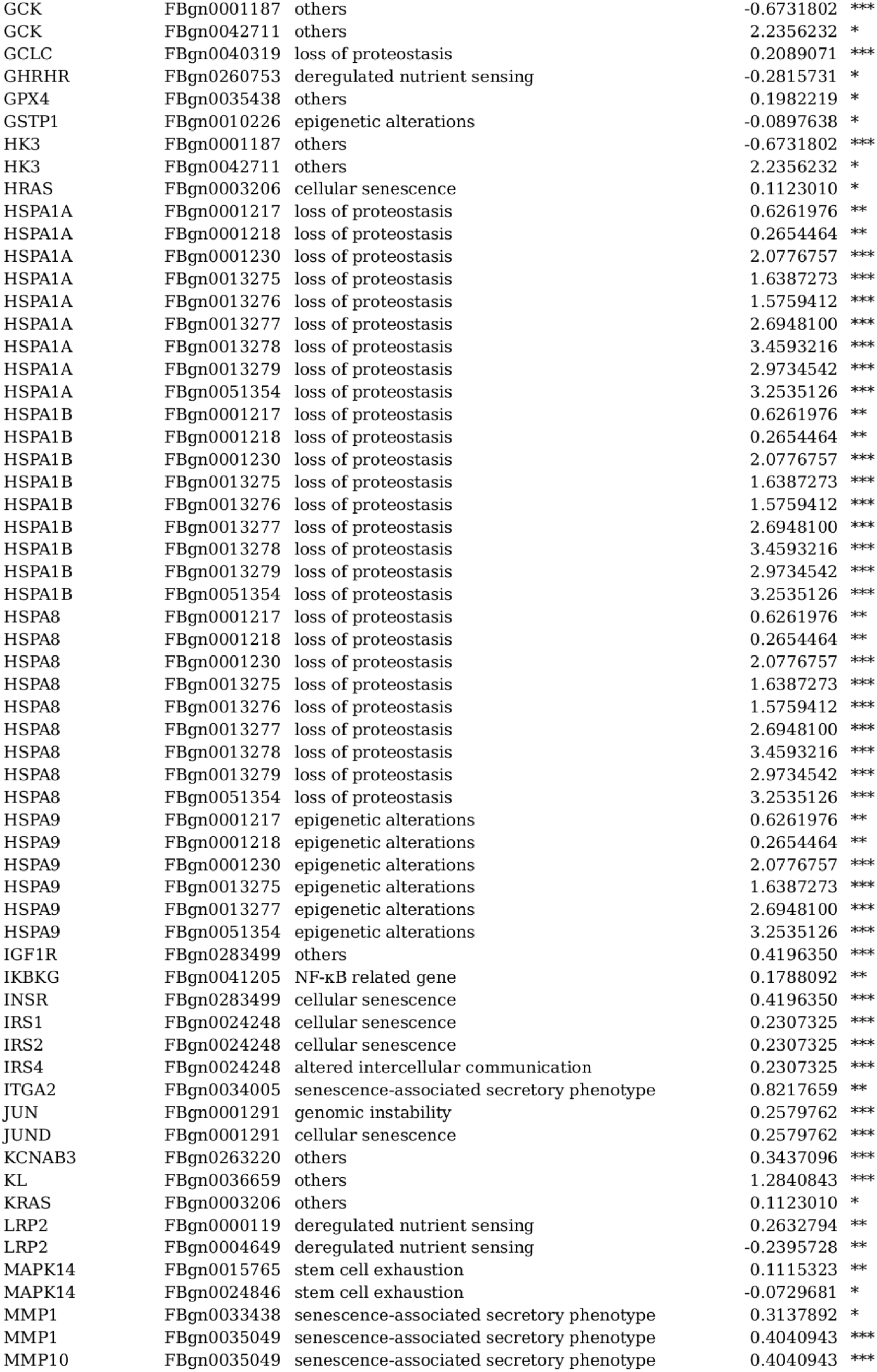
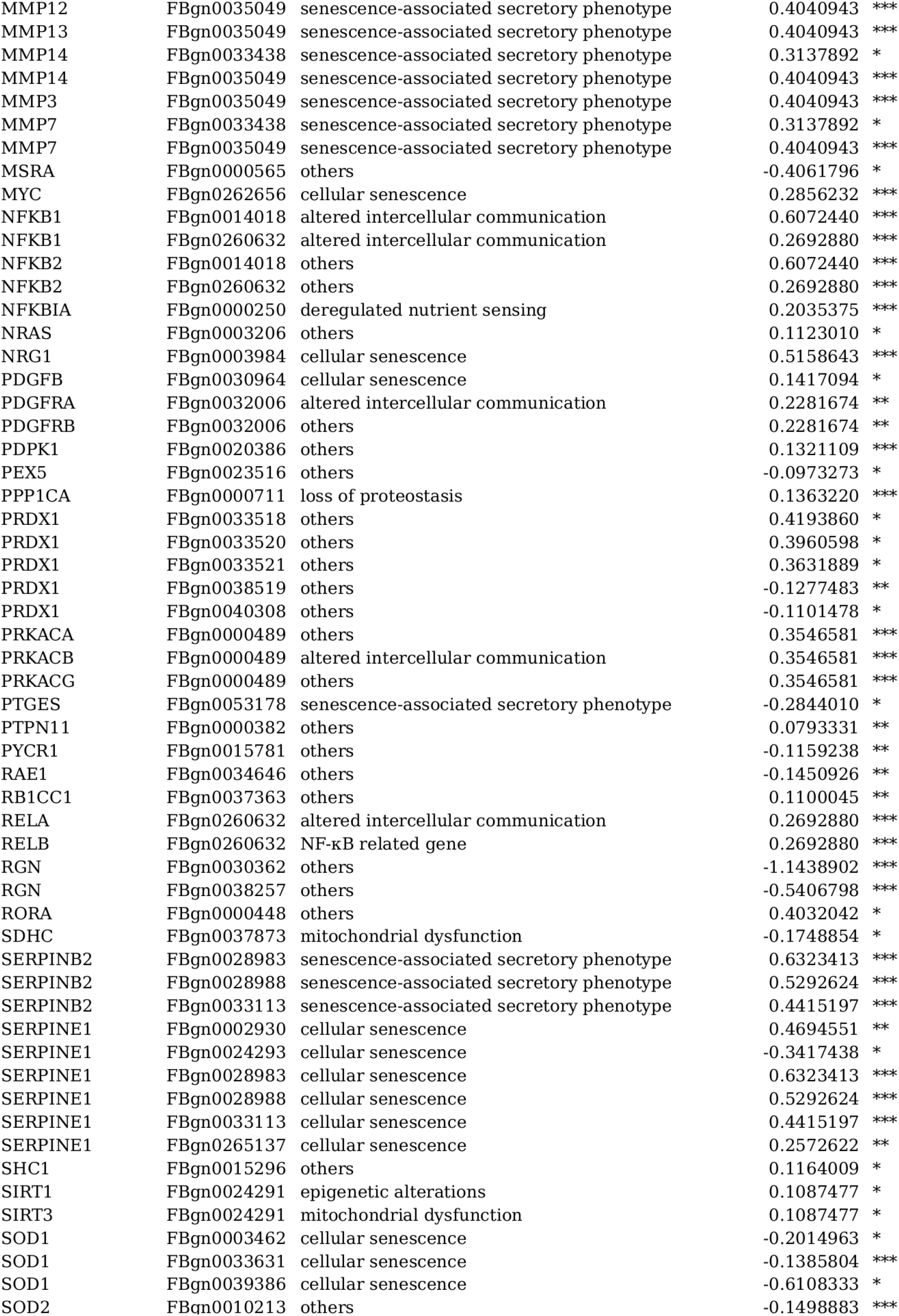
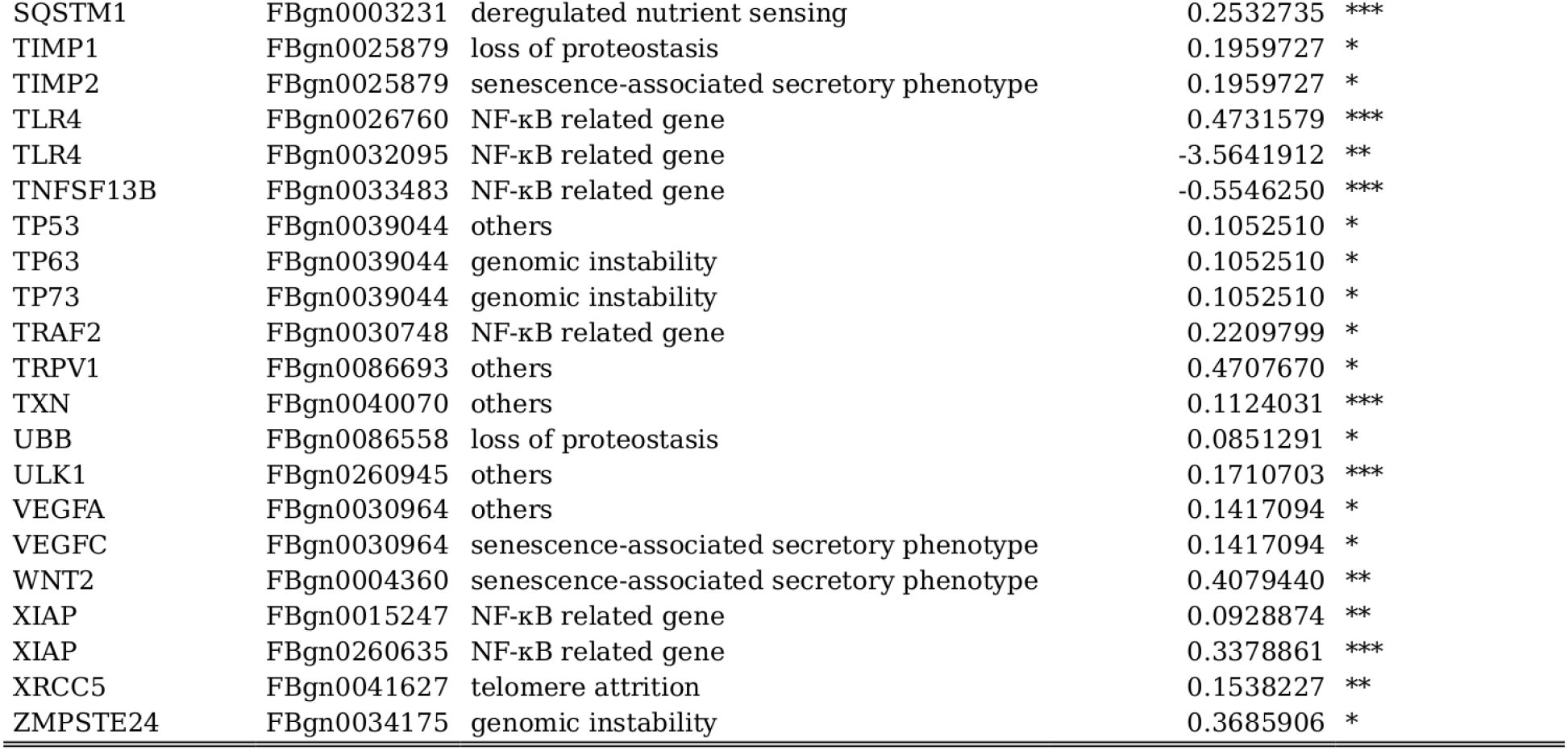
Human genes from Ageing Atlas mapping to Smurf DEGs. A total of 134 (unique) human genes are retrieved by overlapping the 500 human genes annotated in the Ageing Atlas to the Smurf DEGs. Note that in the table some human genes are “duplicated” as they map to more than one fly gene, and the opposite. In total, 121 unique fly genes are found. Human symbol: human gene name; Flybase: Drosophila gene, flybase ID; Ageing marker: ageing marker annotated to the human gene (12 in total defined); log2FC (DESeq2): log2FC estimated by DESeq2 in the Smurf/non-Smurf analysis; FDR (DESeq2): adjusted p-value, FDR method, *** FDR < 0.001, ** FDR < 0.01, * FDR < 0.05.

**Table S6.**
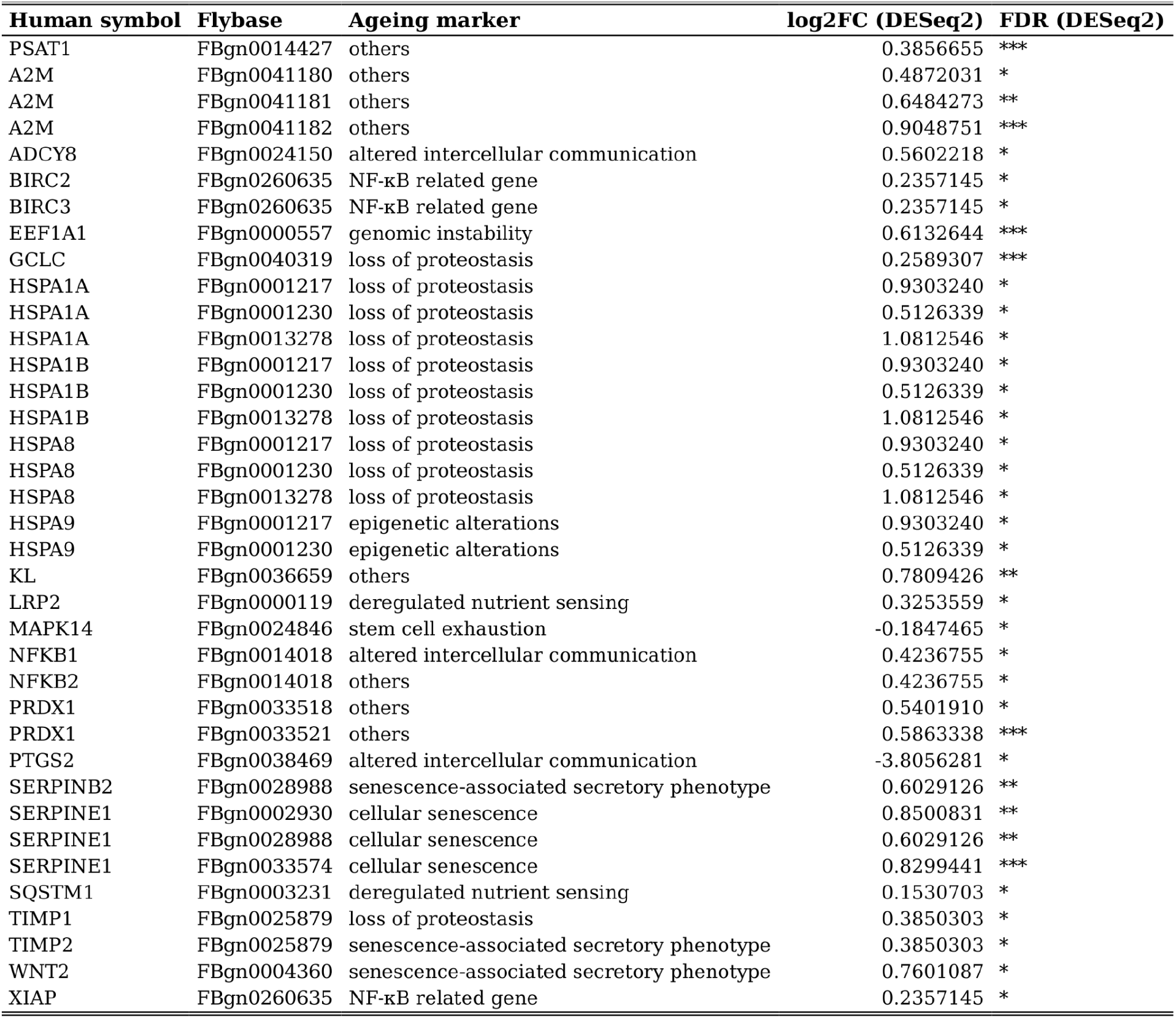
Human genes from Ageing Atlas mapping to non-Smurf DEGs. A total of 25 (unique) human genes are retrieved by overlapping the 500 human genes annotated in the Ageing Atlas to the old non-Smurf DEGs. Note that in the table some human genes are “duplicated” as they map to more than one fly gene, and the opposite. In total, 24 unique fly genes are found. Human symbol: human gene name; Flybase: Drosophila gene, flybase ID; Ageing marker: ageing marker annotated to the human gene (12 in total defined); log2FC (DESeq2): log2FC estimated by DESeq2 in the Smurf/non-Smurf analysis; FDR (DESeq2): adjusted p-value, FDR method, *** FDR < 0.001, ** FDR < 0.01, * FDR < 0.05.

**Table S7.**
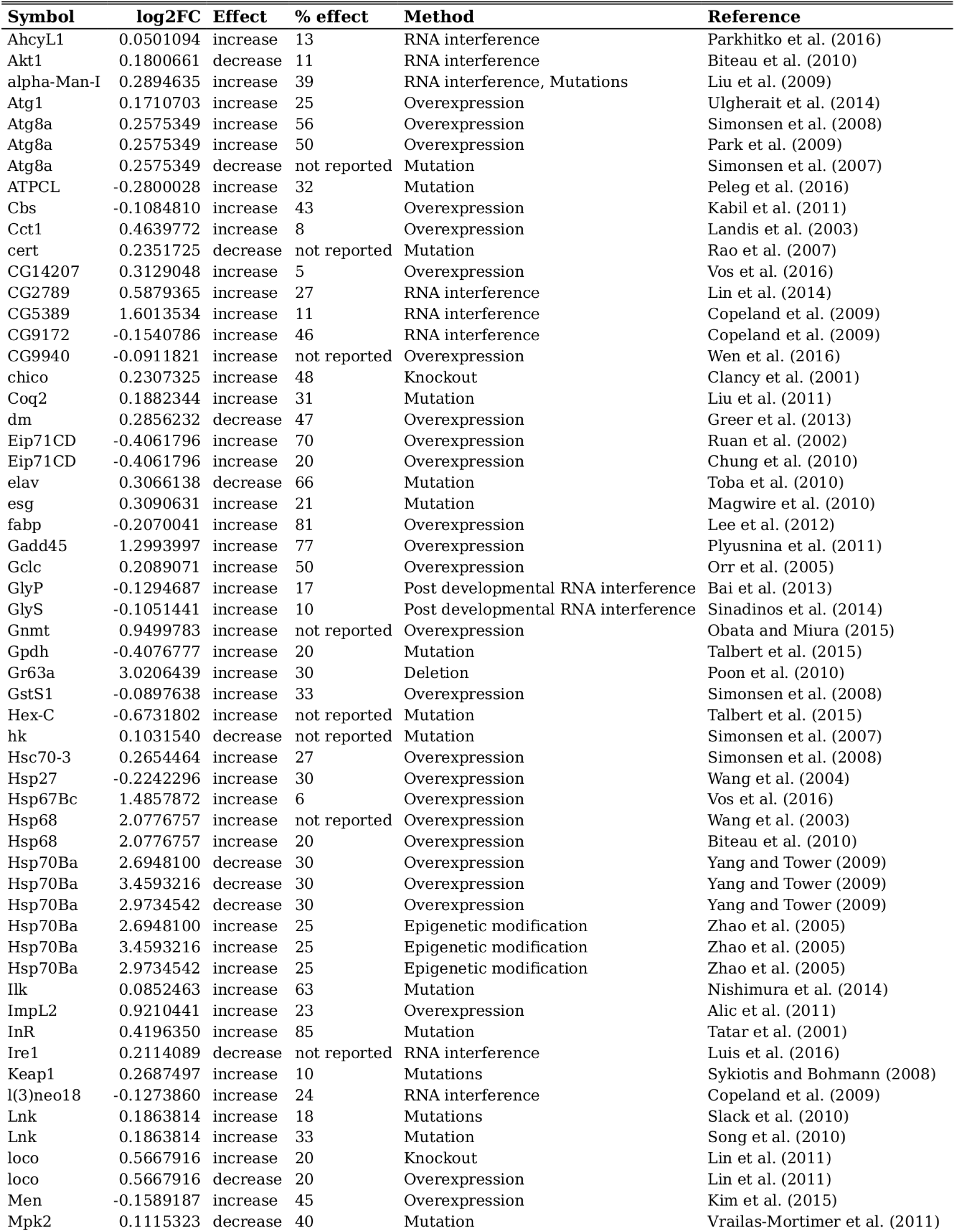
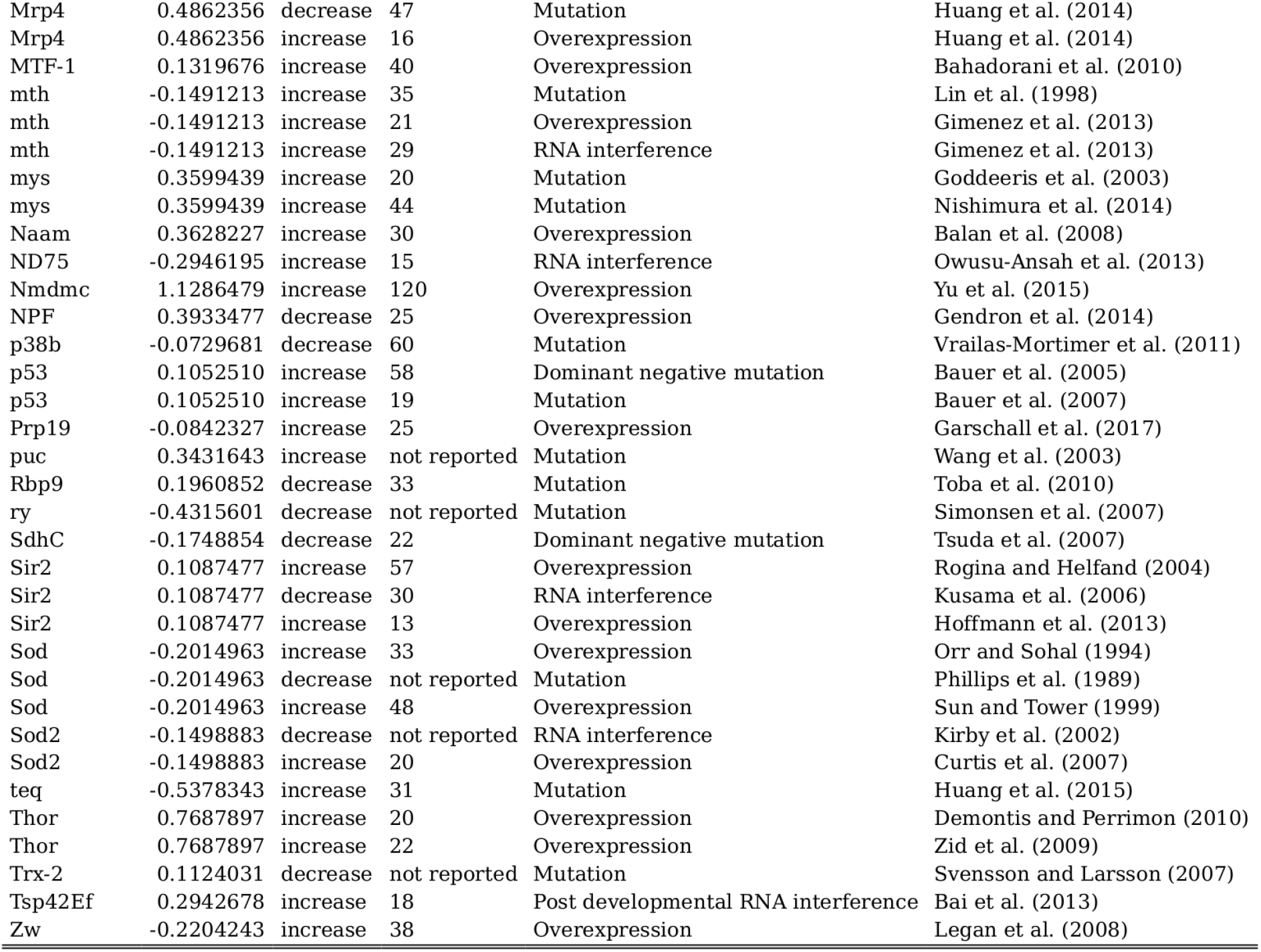
Drosophila longevity genes (GenAge) mapping to Smurf DEGs. Drosophila longevity genes (annotated in GenAge) mapping to Smurf DEGs. A total of 58 unique genes are identified. Note that the table contains duplicated gene symbols as multiple experiments can be reported for one gene. Symbol: Drosophila gene symbol; log_2_FC: log_2_FC Smurf/non-Smurfs estimated by DESeq2; effect: effect of the alteration lifespan; % effect: change in mean lifespan, in %; method: type of experiment performed; reference: reference of the study.

**Table S8.**
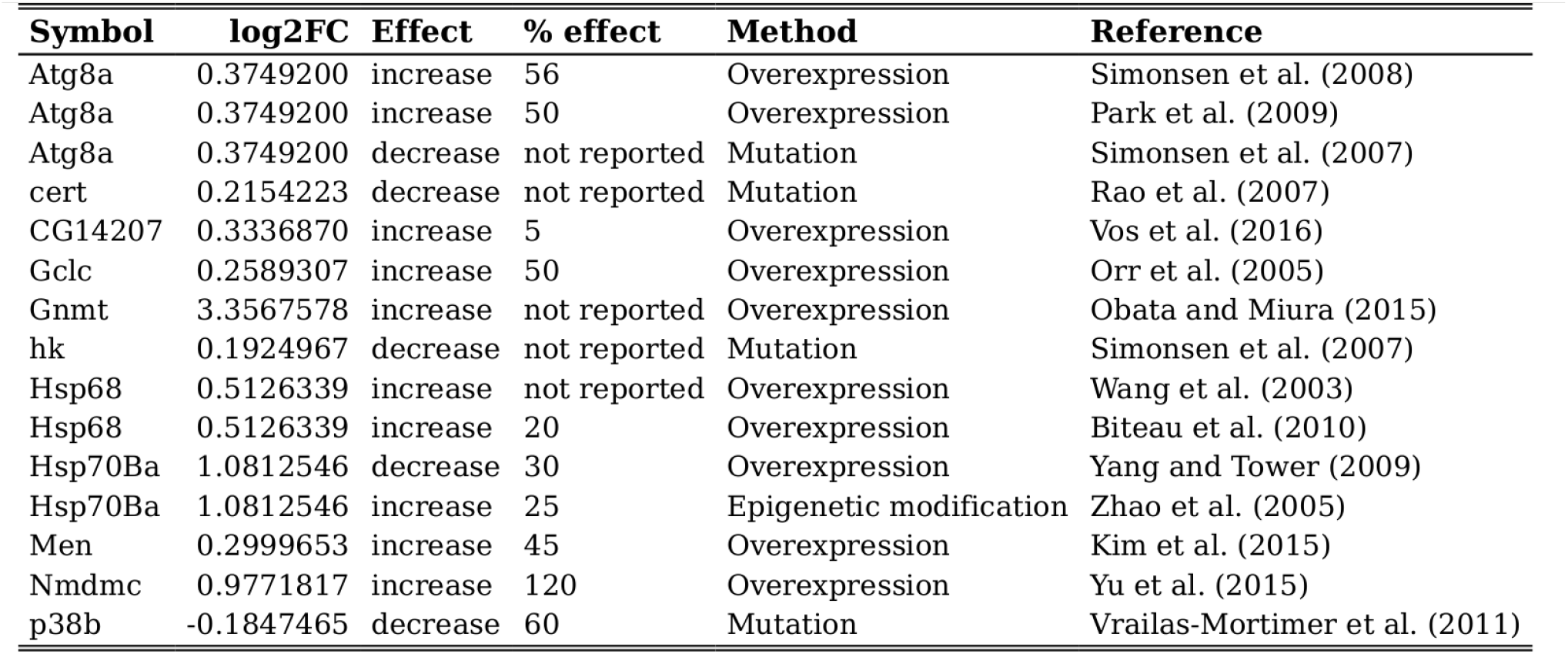
Drosophila longevity genes (GenAge) mapping to non-Smurf DEGs. Drosophila longevity genes (annotated in GenAge) mapping to old non-Smurf DEGs. A total of 11 unique genes are identified. Note that the table contains duplicated gene symbols as multiple experiments can be reported for one gene. Symbol: Drosophila gene symbol; log_2_FC: log_2_FC 20 days non-Smurf/40 days non-Smurfs estimated by DESeq2; effect: effect of the alteration lifespan; % effect: change in mean lifespan, in %; method: type of experiment performed; reference: reference of the study.

**Table S9.**
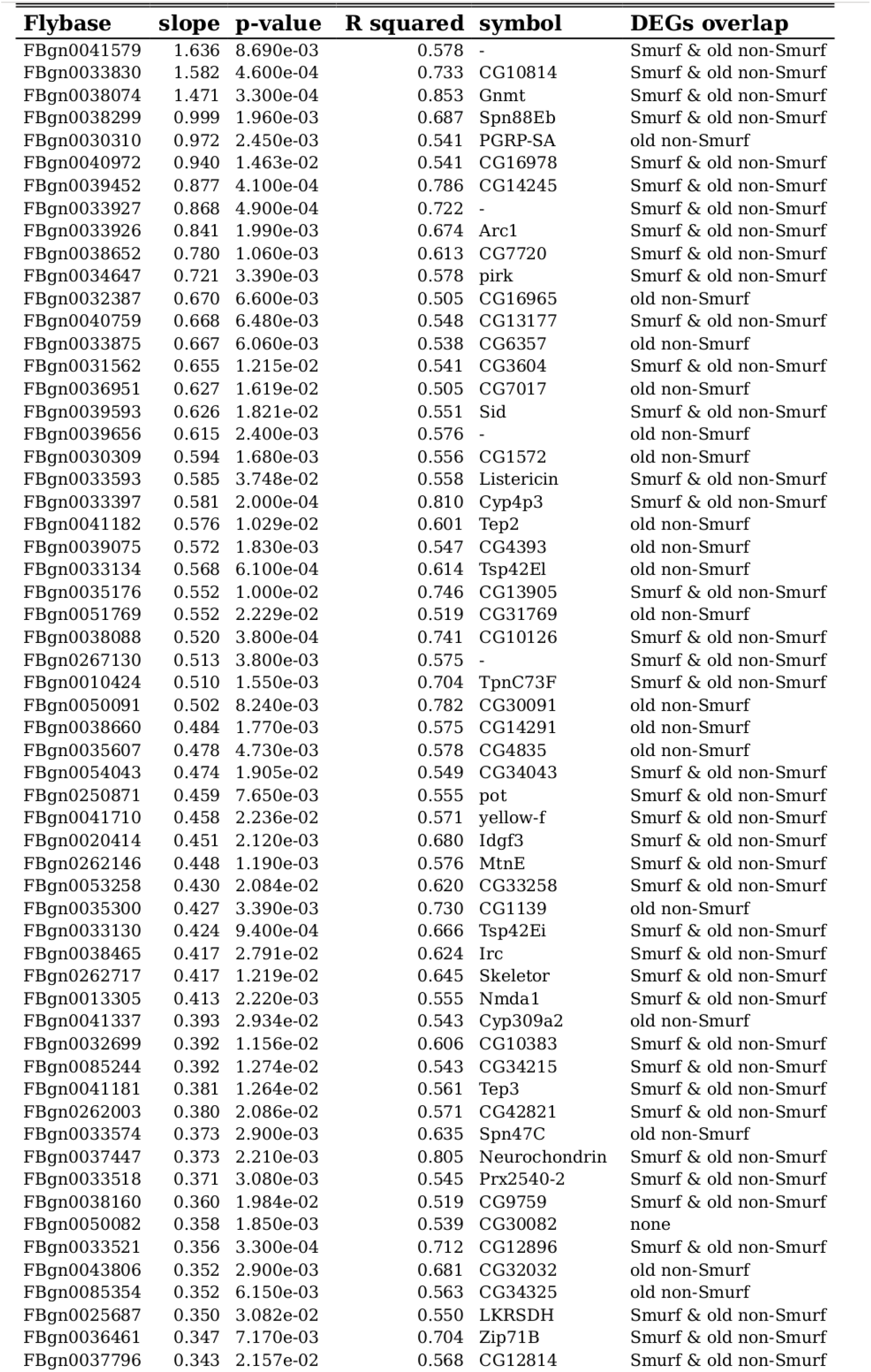
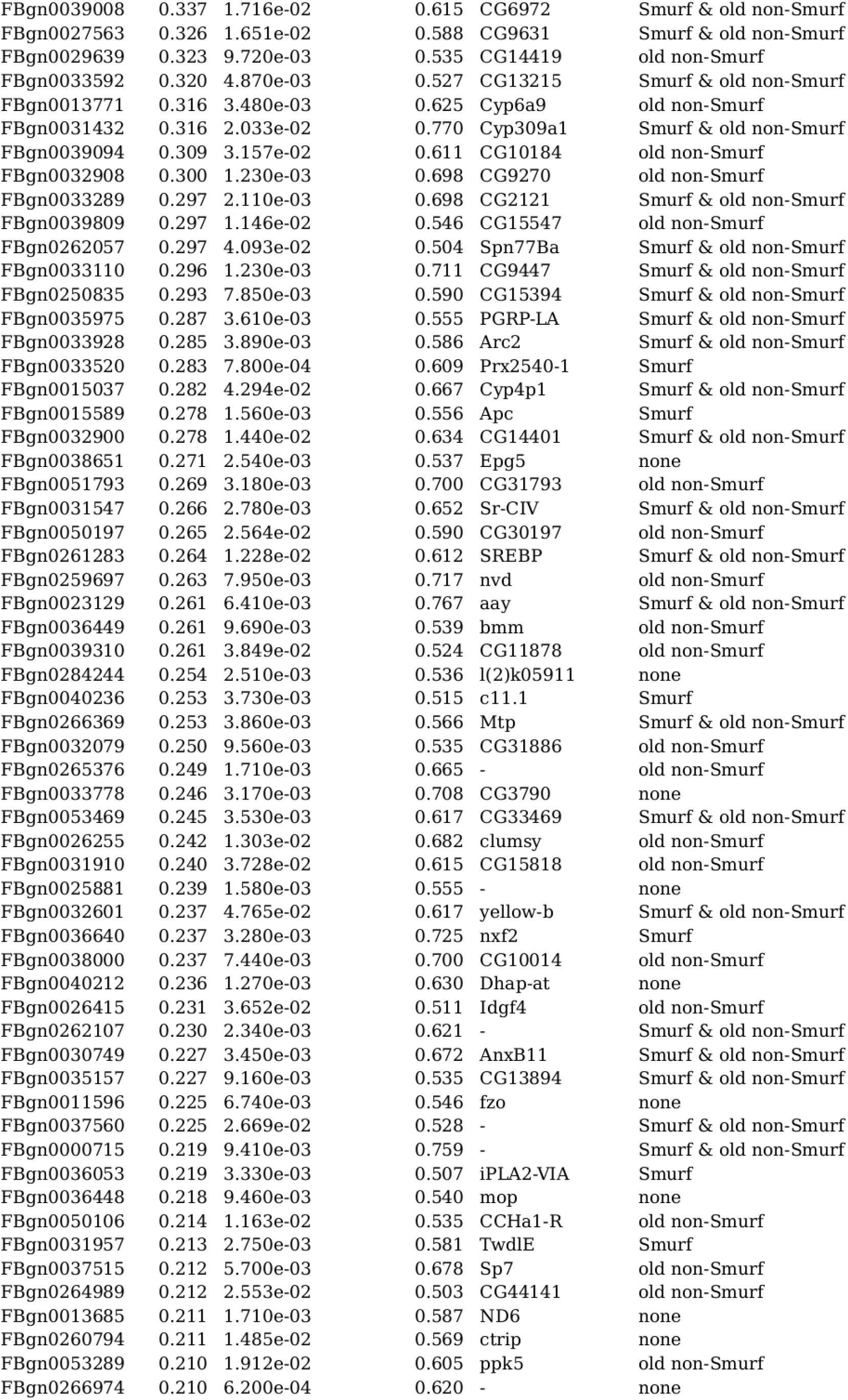
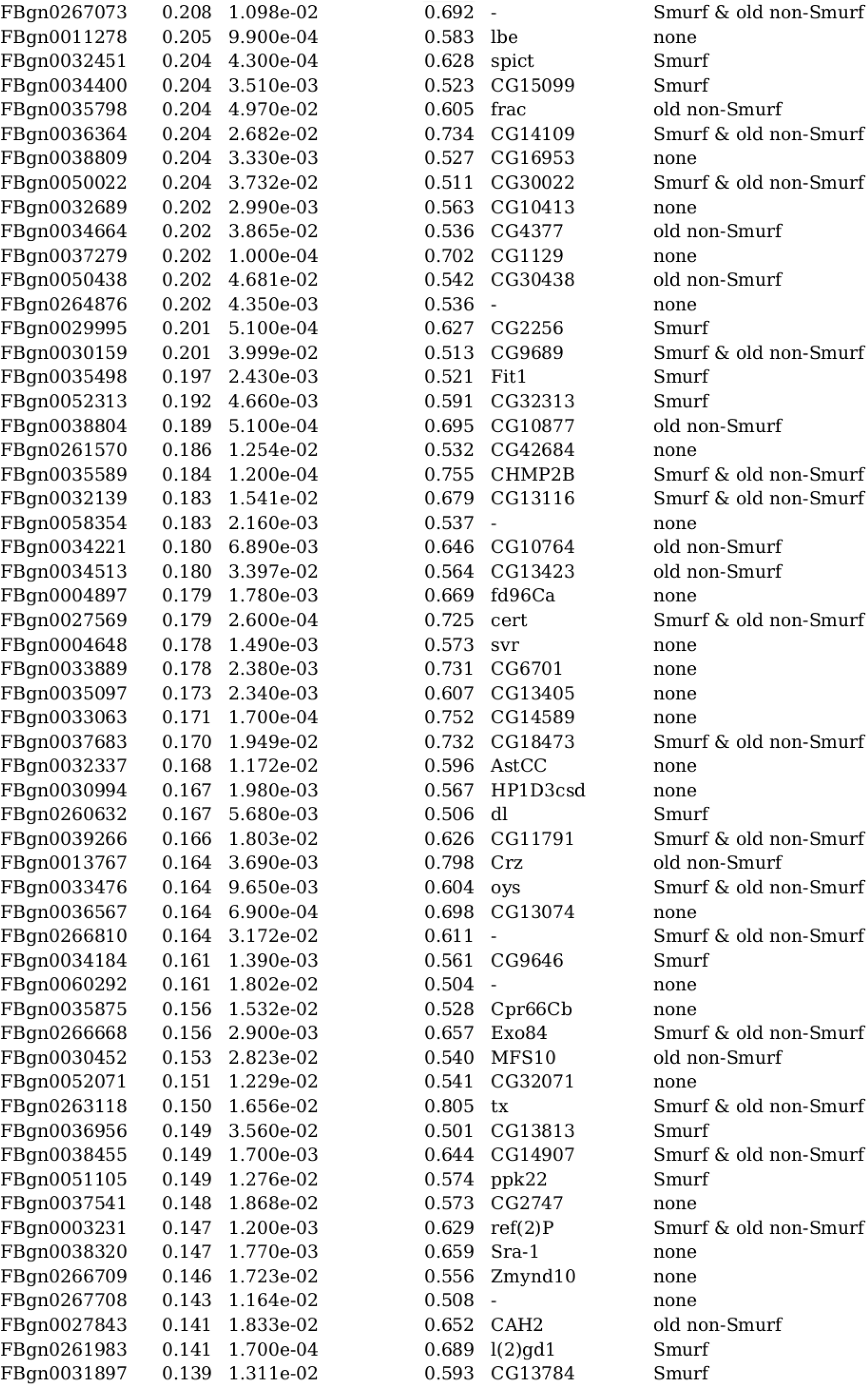
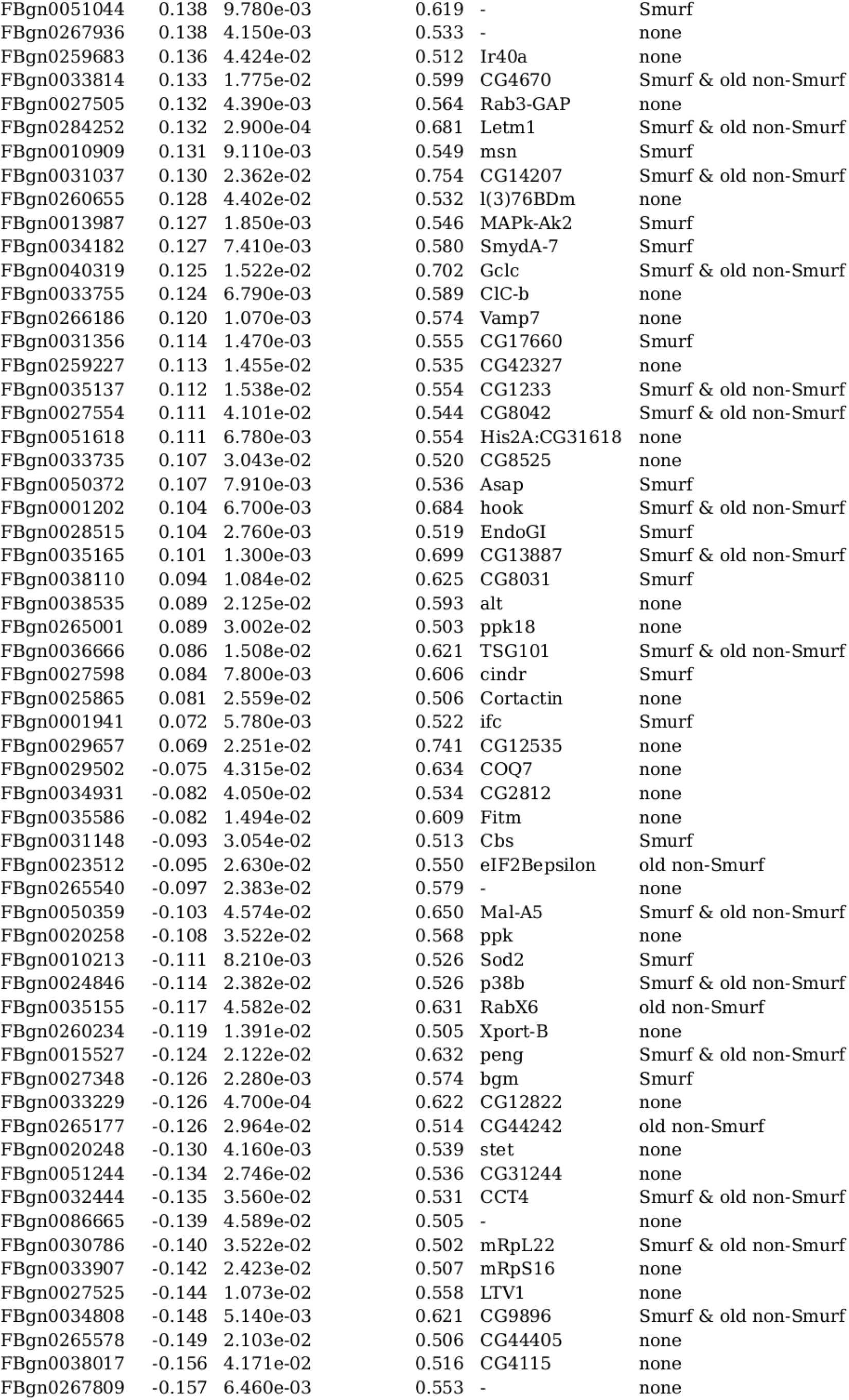
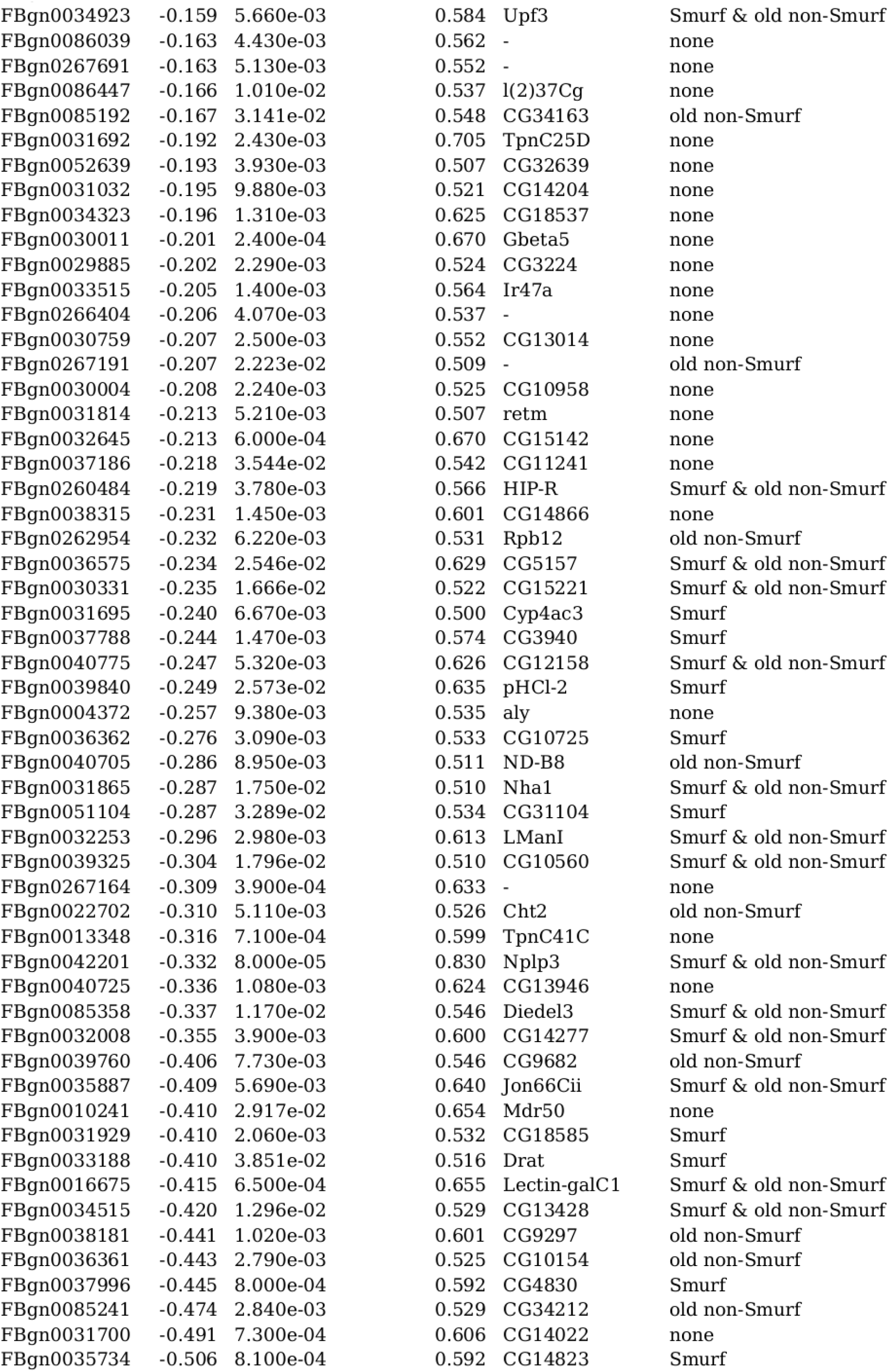
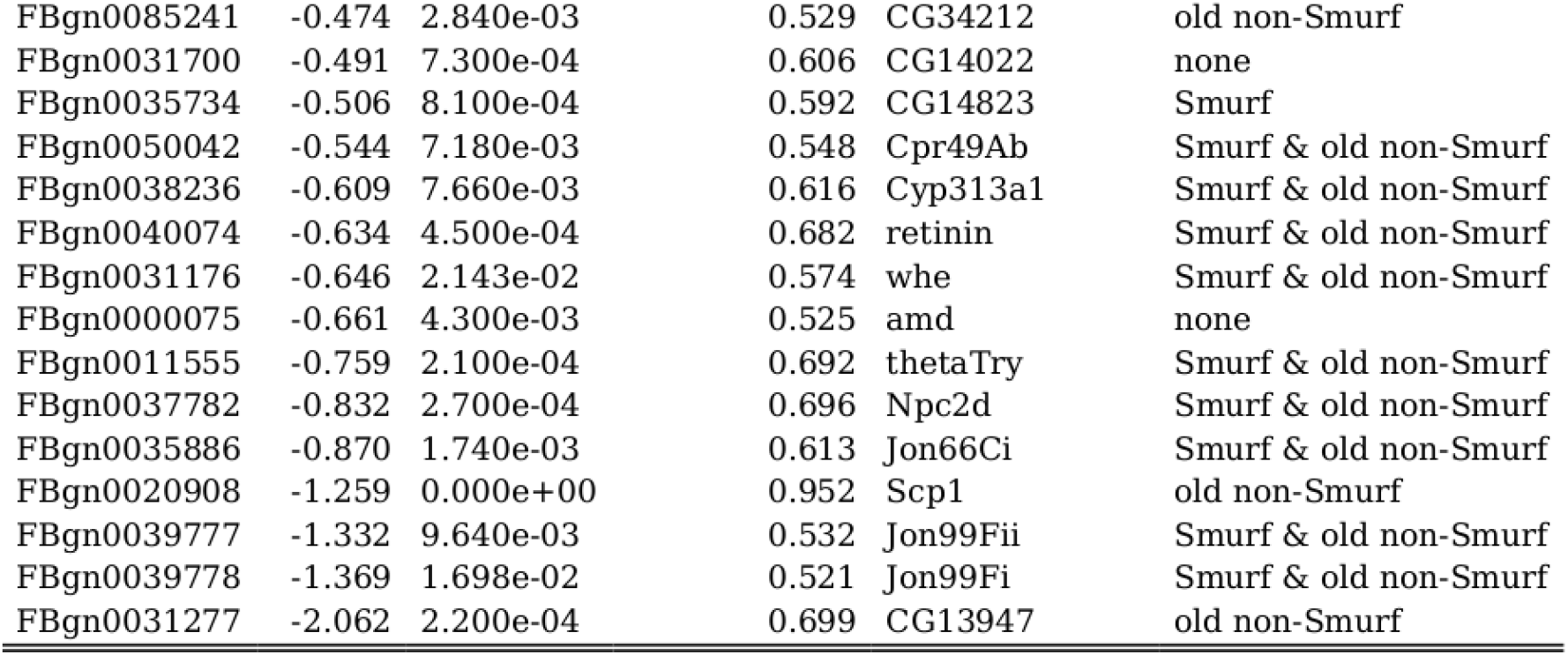
Linear regression on non-Smurfs gene expression (time dependence). The 301 genes with significant slope over time in non-Smurfs, with r^2^ > 0.5. Genes are ordered by descending slope value. Flybase: flybase ID; slope: β_1_ of the linear regression; p-value: F-statistic p-value; R squared: r^2^ of the estimated linear regression; symbol: Gene symbol; DEGs overlap: specifies if the genes has been detected as significantly deregulated in Smurfs, old non-Smurfs, both or none.

**Table S10.** KEGG pathways affected by Smurfness. The 48 pathways identified as affected more by Smurfness than chronological age according to our expression dataset. KEGG path: KEGG ID and pathway name; Avg age correlation: average gene expression correlation with chronological age on the genes belonging to the pathway; Avg Smurf correlation: average gene expression correlation with Smurf on the genes belonging to the pathway; adjust pval (Fasano-Franceschini): adjusted p-value (FDR) from the Fasano-Franceschini test.

| KEGG path | Avg age correlation | Avg Smurf correlation | adjust p-val (Fasano-Franceschini) |
| --- | --- | --- | --- |
| dme04624 Toll and Imd signaling pathway | 0.080 | 0.248 | 5.2e-06 |
| dme00565 Ether lipid metabolism | 0.096 | 0.232 | 6.3e-03 |
| dme04130 SNARE interactions in vesicular transport | -0.097 | 0.229 | 5.8e-04 |
| dme04144 Endocytosis | -0.084 | 0.207 | 1.5e-08 |
| dme04213 Longevity regulating pathway - multiple species | 0.066 | 0.166 | 9.6e-03 |
| dme04391 Hippo signaling pathway - fly | -0.023 | 0.165 | 3.5e-02 |
| dme04350 TGF-beta signaling pathway | 0.022 | 0.149 | 1.9e-02 |
| dme04070 Phosphatidylinositol signaling system | -0.025 | 0.138 | 3.9e-02 |
| dme00564 Glycerophospholipid metabolism | 0.111 | 0.134 | 3.9e-03 |
| dme04745 Phototransduction - fly | 0.081 | 0.129 | 2.8e-02 |
| dme04068 FoxO signaling pathway | -0.037 | 0.128 | 4.6e-03 |
| dme04013 MAPK signaling pathway - fly | -0.028 | 0.114 | 1.8e-03 |
| dme00480 Glutathione metabolism | 0.054 | 0.088 | 1.1e-02 |
| dme01100 Metabolic pathways | 0.083 | -0.090 | 3.7e-33 |
| dme00240 Pyrimidine metabolism | 0.032 | -0.091 | 9.1e-03 |
| dme00310 Lysine degradation | -0.065 | -0.103 | 1.7e-03 |
| dme00010 Glycolysis / Gluconeogenesis | 0.051 | -0.143 | 3.5e-03 |
| dme00620 Pyruvate metabolism | 0.053 | -0.147 | 3.8e-03 |
| dme00330 Arginine and proline metabolism | 0.073 | -0.169 | 8.3e-03 |
| dme03010 Ribosome | -0.012 | -0.191 | 9.8e-17 |
| dme03008 Ribosome biogenesis in eukaryotes | -0.159 | -0.203 | 4.0e-10 |
| dme01230 Biosynthesis of amino acids | 0.124 | -0.212 | 1.2e-08 |
| dme04146 Peroxisome | 0.065 | -0.214 | 1.9e-07 |
| dme00450 Selenocompound metabolism | -0.109 | -0.216 | 6.8e-03 |
| dme00190 Oxidative phosphorylation | 0.088 | -0.217 | 4.5e-15 |
| dme00270 Cysteine and methionine metabolism | 0.150 | -0.218 | 2.5e-05 |
| dme03020 RNA polymerase | -0.186 | -0.219 | 2.3e-04 |
| dme01200 Carbon metabolism | 0.064 | -0.222 | 1.7e-11 |
| dme00650 Butanoate metabolism | 0.013 | -0.225 | 5.0e-04 |
| dme00981 Insect hormone biosynthesis | 0.194 | -0.235 | 1.1e-04 |
| dme04512 ECM-receptor interaction | 0.156 | -0.235 | 3.8e-03 |
| dme01040 Biosynthesis of unsaturated fatty acids | 0.023 | -0.240 | 1.7e-03 |
| dme00030 Pentose phosphate pathway | 0.107 | -0.245 | 4.2e-03 |
| dme00250 Alanine, aspartate and glutamate metabolism | 0.209 | -0.249 | 3.1e-05 |
| dme00380 Tryptophan metabolism | 0.083 | -0.249 | 8.0e-05 |
| dme00062 Fatty acid elongation | -0.031 | -0.255 | 3.8e-03 |
| dme00220 Arginine biosynthesis | 0.214 | -0.263 | 4.0e-03 |
| dme00630 Glyoxylate and dicarboxylate metabolism | 0.145 | -0.267 | 2.1e-05 |
| dme00531 Glycosaminoglycan degradation | 0.136 | -0.270 | 1.5e-02 |
| dme00020 Citrate cycle (TCA cycle) | 0.062 | -0.274 | 1.5e-06 |
| dme00280 Valine, leucine and isoleucine degradation | -0.028 | -0.281 | 9.7e-07 |
| dme01212 Fatty acid metabolism | 0.035 | -0.281 | 1.5e-08 |
| dme01210 2-Oxocarboxylic acid metabolism | 0.158 | -0.299 | 2.1e-04 |
| dme00640 Propanoate metabolism | 0.005 | -0.303 | 1.1e-05 |
| dme00410 beta-Alanine metabolism | 0.024 | -0.325 | 2.2e-05 |
| dme00061 Fatty acid biosynthesis | 0.216 | -0.326 | 1.6e-03 |
| dme00511 Other glycan degradation | 0.116 | -0.369 | 1.3e-03 |
| dme00071 Fatty acid degradation | -0.063 | -0.388 | 4.3e-09 |

**Table S11.** KEGG pathways affected by chronological age. The 38 pathways identified as affected more by chronological age than Smurfness according to our expression dataset. KEGG path: KEGG ID and pathway name; Avg age correlation: average gene expression correlation with chronological age of the genes belonging to the pathway; Avg Smurf correlation: average gene expression correlation with Smurf on the genes belonging to the pathway; adjust pval (Fasano-Franceschini): adjusted p-value (FDR) from the Fasano-Franceschini test.

| KEGG path | Avg age correlation | Avg Smurf correlation | adjust p-val (Fasano-Franceschini) |
| --- | --- | --- | --- |
| dme00052 Galactose metabolism | 0.273 | -0.087 | 1.7e-04 |
| dme00670 One carbon pool by folate | 0.261 | -0.207 | 1.7e-03 |
| dme00260 Glycine, serine and threonine metabolism | 0.252 | -0.196 | 1.5e-06 |
| dme00830 Retinol metabolism | 0.240 | -0.099 | 3.3e-05 |
| dme00053 Ascorbate and aldarate metabolism | 0.232 | -0.125 | 1.6e-05 |
| dme00040 Pentose and glucuronate interconversions | 0.227 | -0.182 | 5.1e-07 |
| dme00500 Starch and sucrose metabolism | 0.219 | -0.129 | 5.5e-05 |
| dme00350 Tyrosine metabolism | 0.201 | -0.175 | 1.2e-03 |
| dme02010 ABC transporters | 0.177 | -0.004 | 5.3e-03 |
| dme04080 Neuroactive ligand-receptor interaction | 0.166 | 0.027 | 1.4e-05 |
| dme00051 Fructose and mannose metabolism | 0.165 | -0.121 | 1.1e-02 |
| dme00983 Drug metabolism - other enzymes | 0.165 | 0.005 | 2.8e-08 |
| dme00770 Pantothenate and CoA biosynthesis | 0.157 | -0.151 | 1.3e-02 |
| dme00980 Metabolism of xenobiotics by cytochrome P450 | 0.157 | 0.050 | 1.4e-07 |
| dme00561 Glycerolipid metabolism | 0.151 | -0.071 | 8.3e-04 |
| dme00982 Drug metabolism - cytochrome P450 | 0.151 | 0.069 | 1.5e-07 |
| dme04142 Lysosome | 0.148 | -0.128 | 3.6e-08 |
| dme00230 Purine metabolism | 0.144 | -0.013 | 6.9e-03 |
| dme00730 Thiamine metabolism | 0.135 | 0.015 | 1.3e-02 |
| dme00520 Amino sugar and nucleotide sugar metabolism | 0.129 | -0.051 | 7.6e-03 |
| dme00790 Folate biosynthesis | 0.123 | 0.027 | 3.5e-02 |
| dme00860 Porphyrin and chlorophyll metabolism | 0.114 | -0.062 | 2.6e-03 |
| dme04120 Ubiquitin mediated proteolysis | -0.154 | 0.012 | 1.5e-06 |
| dme03015 mRNA surveillance pathway | -0.177 | -0.066 | 1.7e-06 |
| dme03018 RNA degradation | -0.191 | -0.136 | 9.1e-09 |
| dme00970 Aminoacyl-tRNA biosynthesis | -0.238 | -0.087 | 1.3e-04 |
| dme03410 Base excision repair | -0.239 | 0.024 | 3.0e-03 |
| dme00563 Glycosylphosphatidylinositol (GPI)-anchor biosynthesis | -0.272 | -0.118 | 3.0e-04 |
| dme04330 Notch signaling pathway | -0.275 | 0.104 | 1.3e-04 |
| dme03050 Proteasome | -0.276 | -0.166 | 3.5e-09 |
| dme03013 RNA transport | -0.288 | -0.117 | 1.3e-15 |
| dme03040 Spliceosome | -0.288 | -0.124 | 1.5e-17 |
| dme03460 Fanconi anemia pathway | -0.312 | 0.020 | 1.3e-06 |
| dme03022 Basal transcription factors | -0.318 | -0.096 | 3.1e-08 |
| dme03420 Nucleotide excision repair | -0.338 | -0.073 | 1.7e-09 |
| dme03440 Homologous recombination | -0.341 | -0.005 | 1.6e-07 |
| dme03430 Mismatch repair | -0.387 | -0.081 | 1.5e-08 |
| dme03030 DNA replication | -0.393 | -0.070 | 2.2e-09 |

**Table S12.**
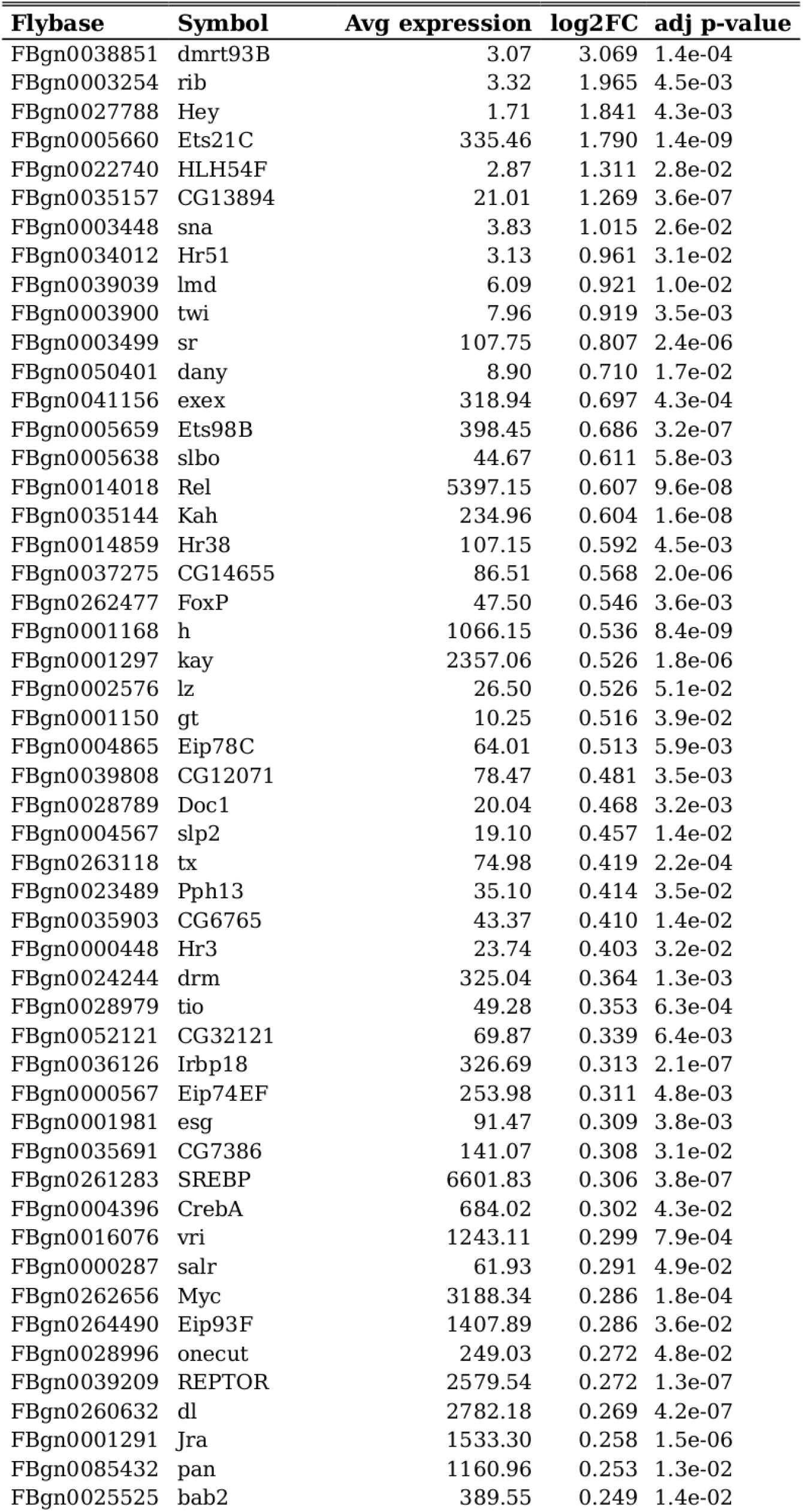
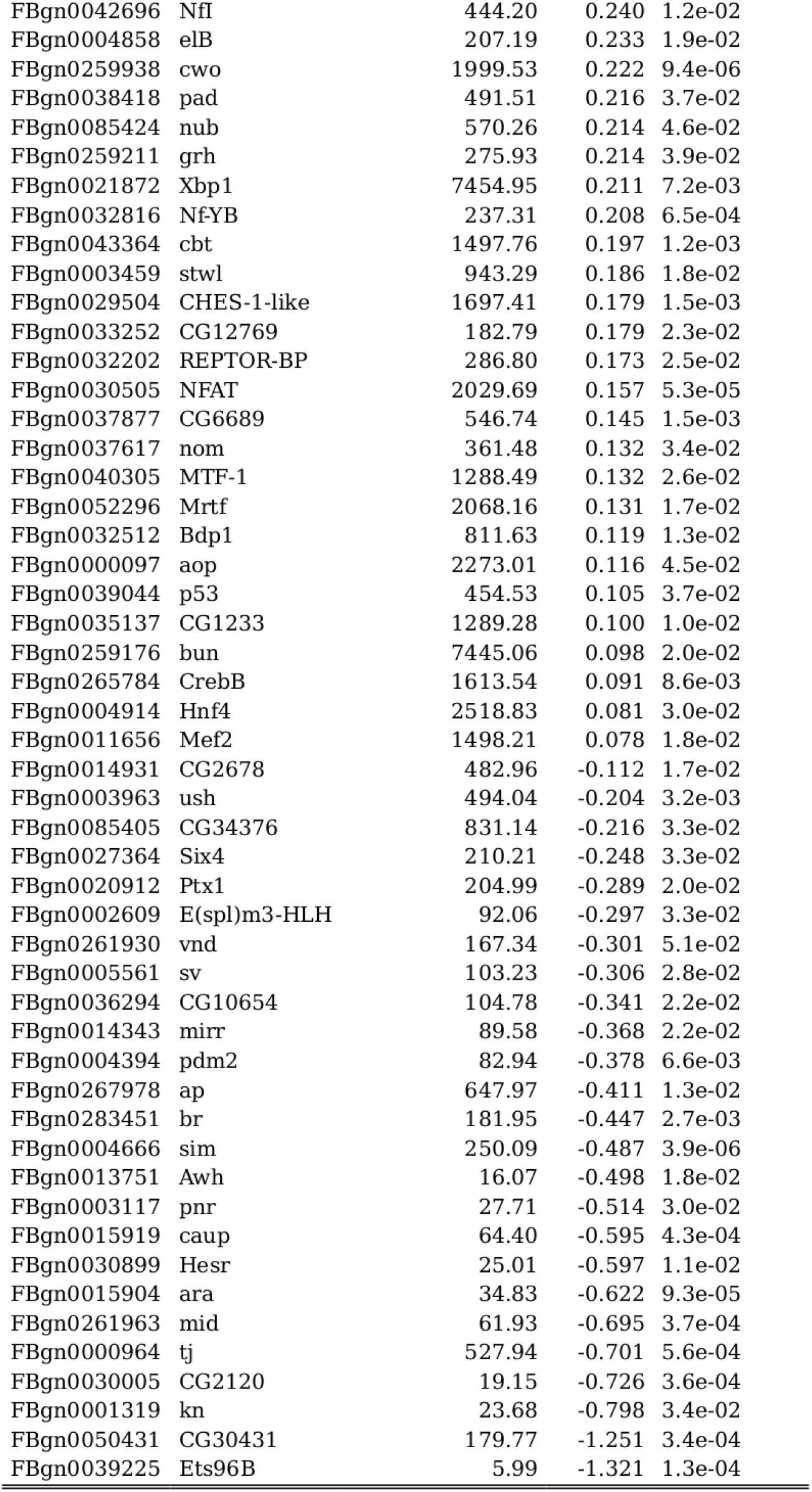
Transcription factors (TFs) deregulated in Smurfs (DESeq2). Table summarizing DESeq2 results for the transcription factors deregulated in Smurfs (ordered by log_2_FC). Flybase: Flybase gene ID; Symbol: gene symbol; Avg expression: average gene expression across samples provided by DESeq2; log_2_FC: log_2_FC estimated by DESeq2; adj p-value: FDR corrected p-value provided by DESeq2.

**Table S13.**
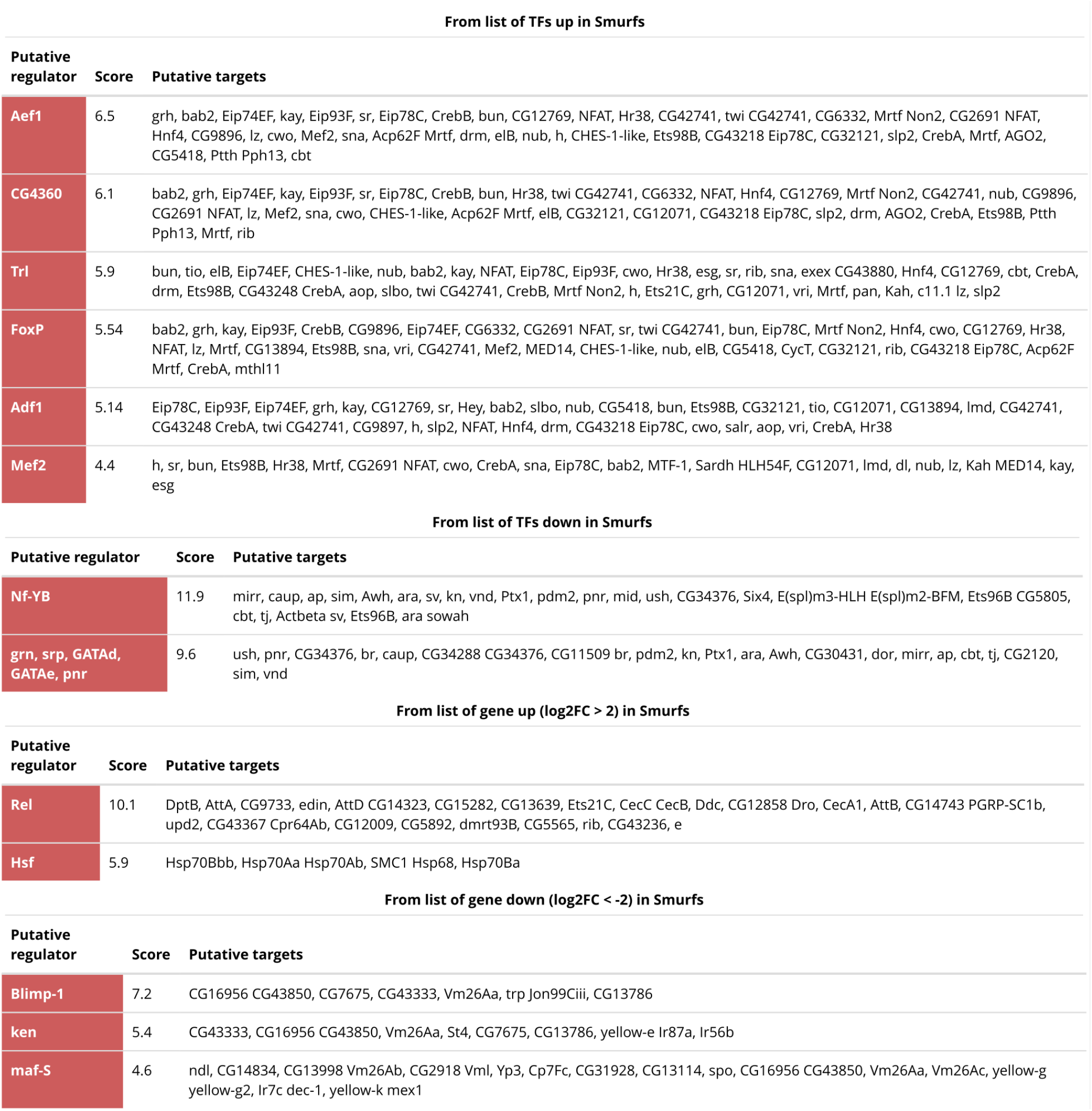
i-cisTarget results. The table reports the best hits provided by i-cisTarget when the queries are 1) TFs upregulated in Smurfs, 2) TFs downregulated in Smurfs, 3) genes upregulated in Smurfs (log_2_FC > 2), genes upregulated in Smurfs (log_2_FC < -2). In all cases the gene symbol, score and putative detected targets are reported.

**Table S14.**
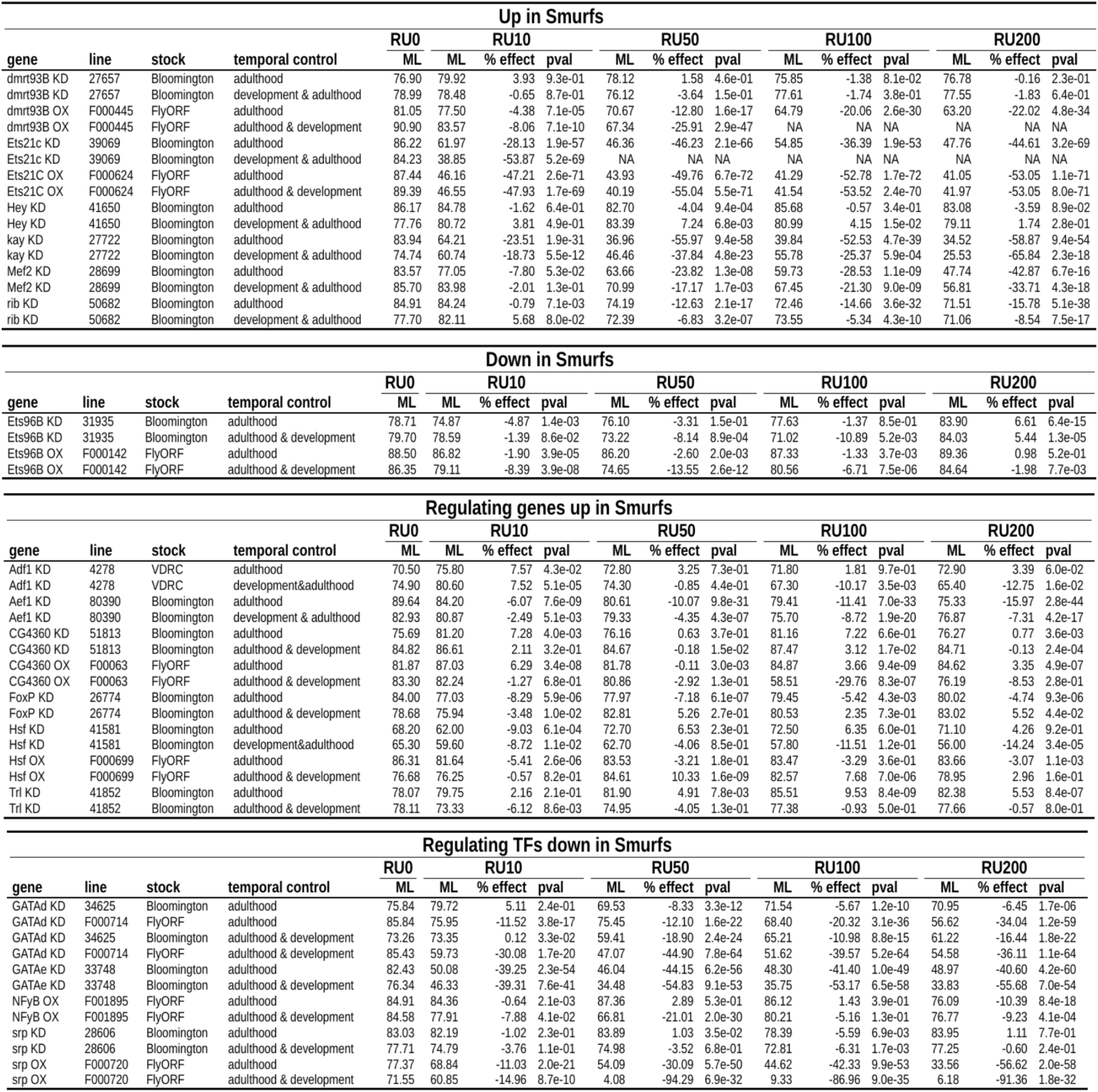
Longevity screening results. Results are organized by groups according to the way the genes were detected (DESeq2 for the first two groups - up and down in Smurfs-, and i-cisTarget for the last two groups - putative regulators of Smurf TFs). Information about the gene and its alteration (KD or OX) are provided, together with the line used and the stock center where the line was bought. Each experiment is either performed during adulthood only or adulthood & developmental (temporal control). Mean lifespan (ML), % effect (% ML change compared to controls) and log-rank p-value are provided for each RU486 condition (RU0 µg/mL - control, RU10 µg/mL, RU50 µg/mL, RU100 µg/mL and RU200 µg/mL). For visual representation of the results, see Fig. S13.

